# 7-Transmembrane Helical (7TMH) Proteins: Pseudo-Symmetry and Conformational Plasticity

**DOI:** 10.1101/465302

**Authors:** Philippe Youkharibache, Alexander Tran, Ravinder Abrol

## Abstract

Membrane proteins sharing 7 transmembrane helices (7-TMH) dominate the polytopic TMH proteome. They cannot be grouped under a monolithic fold or superfold, however, a parallel structural analysis of folds around that magic number of 7-TMH in distinct 6/7/8-TMH protein superfamilies (SWEET, PnuC, TRIC, FocA, Aquaporin, GPCRs, AND MFS), reveals a common homology, not in their structural fold, but in their systematic pseudo-symmetric construction. Our analysis leads to guiding principles of intragenic duplication and pseudo-symmetric assembly of ancestral 3 or 4 Transmembrane Helix (3/4-TMH) protodomains/protofolds. A parallel deconstruction and reconstruction of these domains provides a structural and mechanistic framework for the evolution path of current pseudo-symmetrical transmembrane helical (TMH) proteins. It highlights the conformational plasticity inherent to fold formation itself. The sequence/structure analysis of different 6/7/8-TMH superfamilies provides a unifying theme of their evolutionary process involving the intragenic duplication of protodomains with varying degrees of sequence and fold divergence under conformational and functional constraints.

## Introduction

### Structural pseudo-symmetry in protein domains

Structural pseudo-symmetry in protein domains has been observed since the early days of structural biology. Ferredoxin, Myohemerythrin, Serine and Aspartyl proteases, Immunoglobulins, the TIM (Triose-phosphate-isomerase) barrels, and the Rossmann fold were among the first crystal structures available. They all exhibit internal pseudo-symmetry (Blundell et al., 1979; Hendrickson and Ward, 1977; McLachlan, 1972, 1987)^,(Barker et al., 1978; Delhaise et al., 1980; Eck and Dayhoff, 1966; Urbain, 1969)^. As these structures appeared, they corroborated earlier sequence-based observations of possible ancestral gene duplications within today’s genes (Barker et al., 1978; Delhaise et al., 1980; Eck and Dayhoff, 1966; Urbain, 1969) and established a basis for interpreting sequence duplication with pseudo-symmetry. This defined, without naming it, what we now call *protodomains*, issued from ancestral protogenes. **A protodomain (or protofold) is a supersecondary structure that by its duplication, symmetry operations (and linkers) can generate a structural domain (fold).**

It is interesting to note that some of these pseudo-symmetric structural domains, characterized early, turned out to be today’s superfolds, some of the most diversified and prototypic folds. In the SCOP classification (Chandonia et al., 2017; Lo Conte et al., 2000), they are denoted: **a.24** (the Myohemerythrin or 4-helix bundle fold) with 28 superfamilies (SFs); **b.1** (The Immunoglobulin fold) with 28 SFs; **c.1** (the TIM barrel) with 33 SFs; and **d.58** (the Ferredoxin fold) with 59 SFs. **The fact that the most diversified folds are pseudo-symmetric suggests a strong evolutionary link between pseudo-symmetry and functional diversification**. We recently performed a census of pseudo-symmetry in the currently known universe of protein domains that shows this link for ∼ 20% of known structural domains (Myers-Turnbull et al., 2014) in the PDB database (see **Table 1**).

**Table 1.**
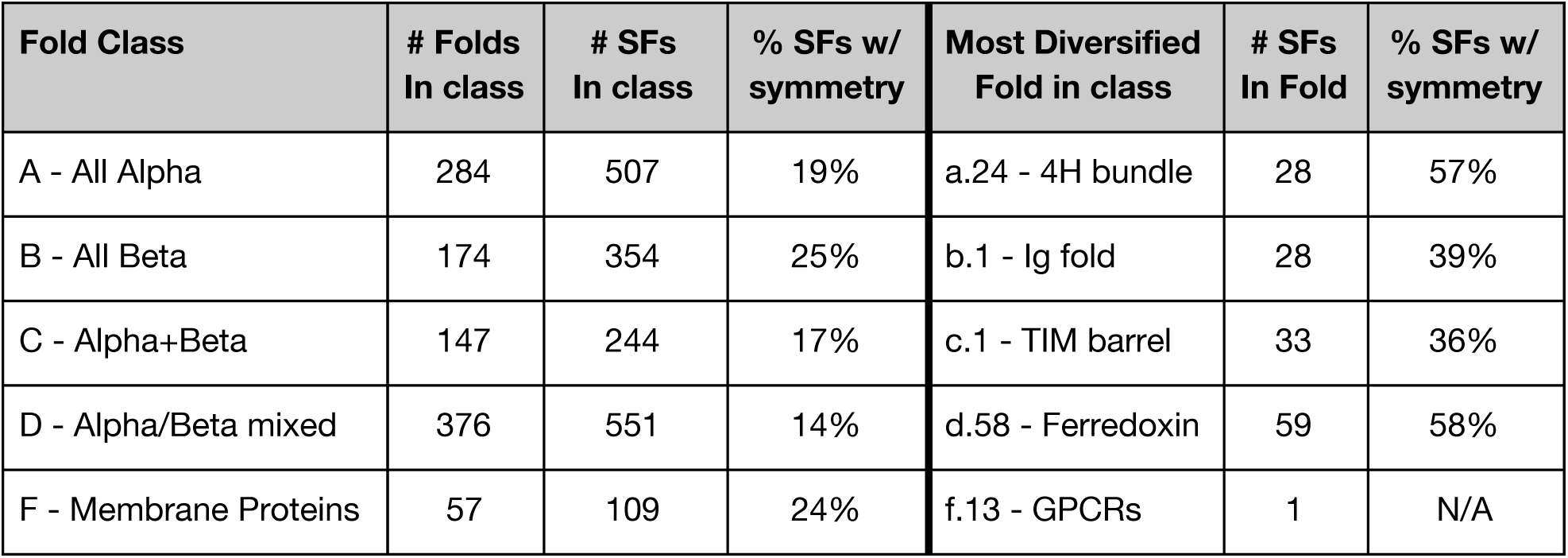
Pseudo-symmetry within Fold classes. (According to SCOP 1.75). For each of the five fold classes in SCOP, this table lists the number of folds, number of Superfamilies (SFs), percentage of SFs deemed symmetrical, most diversified fold in each class, i.e. folds with the highest number of Superfamilies in each class, and the percentage of Superfamilies exhibiting pseudo-symmetry in the most diversified fold (see Table S2 in Ref. (Myers-Turnbull et al., 2014) for details). We added GPCRs, classified as one-fold one-family in SCOP. Technically, it could also be classified as an all-alpha fold (A). The latest SCOPe 2.07 numbers are marginally higher and count 60 membrane protein folds to date(Chandonia et al., 2017).

#### Structural pseudo-symmetry in membrane protein domains

The pool of known membrane proteins is currently comprised of 71% of ɑ-helical structures, 19% of β-sheet structures and the remaining 10% being classified as monotypic. They are, however, classified as one distinct class (F) within SCOP (Chandonia et al., 2017; Lo Conte et al., 2000), regardless of their secondary structure makeup. Overall, they show a higher pseudo-symmetry rate (24%) than the all-ɑ class (A) of globular proteins (19%) (see **Table 1**). That number is likely to be a minimum, as we used a conservative approach in its determination (Myers-Turnbull et al., 2014). As stated in the original census, the criteria used underestimate the number of symmetric superfamilies in SCOP by about 27% to avoid false positives. Clearly, membrane proteins, as a class, pose a challenge for an accurate estimation of pseudo-symmetry, given the relatively low number of known superfamilies per fold, the relatively low number of structurally characterized superfamilies themselves, and simply a much lower number of known structures (with close to 5,000 membrane protein structures vs. ∼140,000 for globular proteins). This has been covered very well in an extensive review on membrane protein symmetry (Forrest, 2015). If we were to use a much less stringent criterion than in the original study to call a fold pseudo-symmetric, we could have a high end estimate of ∼40% for the class of membrane proteins, rather than the conservative 24% in **Table 1**, and closer to prior estimates (Choi et al., 2008; Forrest, 2015; Hennerdal et al., 2010).

An aim of the current analysis was to find salient evolutionary patterns, of protodomains with respect to each other in a set of pseudo-symmetric membrane protein folds covering 6/7/8-TMH protein families.

##### Structural pseudo-symmetry in GPCRs

Although pseudo-symmetry had been noticed previously in Rhodopsin (Choi et al., 2008; Youkharibache - Unpublished Results) no systematic study analyzing GPCRs’ pseudo-symmetry and corresponding protodomains alignments has been performed to date. This is mainly because GPCRs are an example of proteins where structural pseudo-symmetry is hard to detect computationally in a systematic manner with current symmetry detection programs (Kim et al., 2010; Myers-Turnbull et al., 2014) (see Methods). For example, in our original pseudo-symmetry census, pseudo-symmetry was detected computationally for only 18% of known GPCR domains. An additional issue is to get accurate protodomain alignments [with a high structural homology or low root-mean-squared deviation (RMSD), see Methods]. This is the reason we use visual inspection and interactive structural alignments as the method of choice. A careful interactive structural self-alignment of each GPCR domain, individually (and in some cases between multiple GPCRs), although tedious, leads to accurate structural protodomains alignments and a solid observation of pseudo-symmetry across all vertebrate GPCR classes (A, B, C, F). Structures of the metabotropic glutamate receptors 1 and 5 are now available (<u>4OR2</u>/<u>5CGD</u>), which belong to the oldest GPCR family (class C) and form obligatory homodimers for their metabotropic functions. They exhibit two levels of symmetry (**Figures 1** through **3** as well as **4.D** later), one at the tertiary structure level and the other at the quaternary structure level in the crystal structures. These symmetries are detected computationally and lead to accurate protodomains alignments. We use metabotropic glutamate receptor 1 as a reference to analyze GPCR structures from all classes.

**Figure 1.**
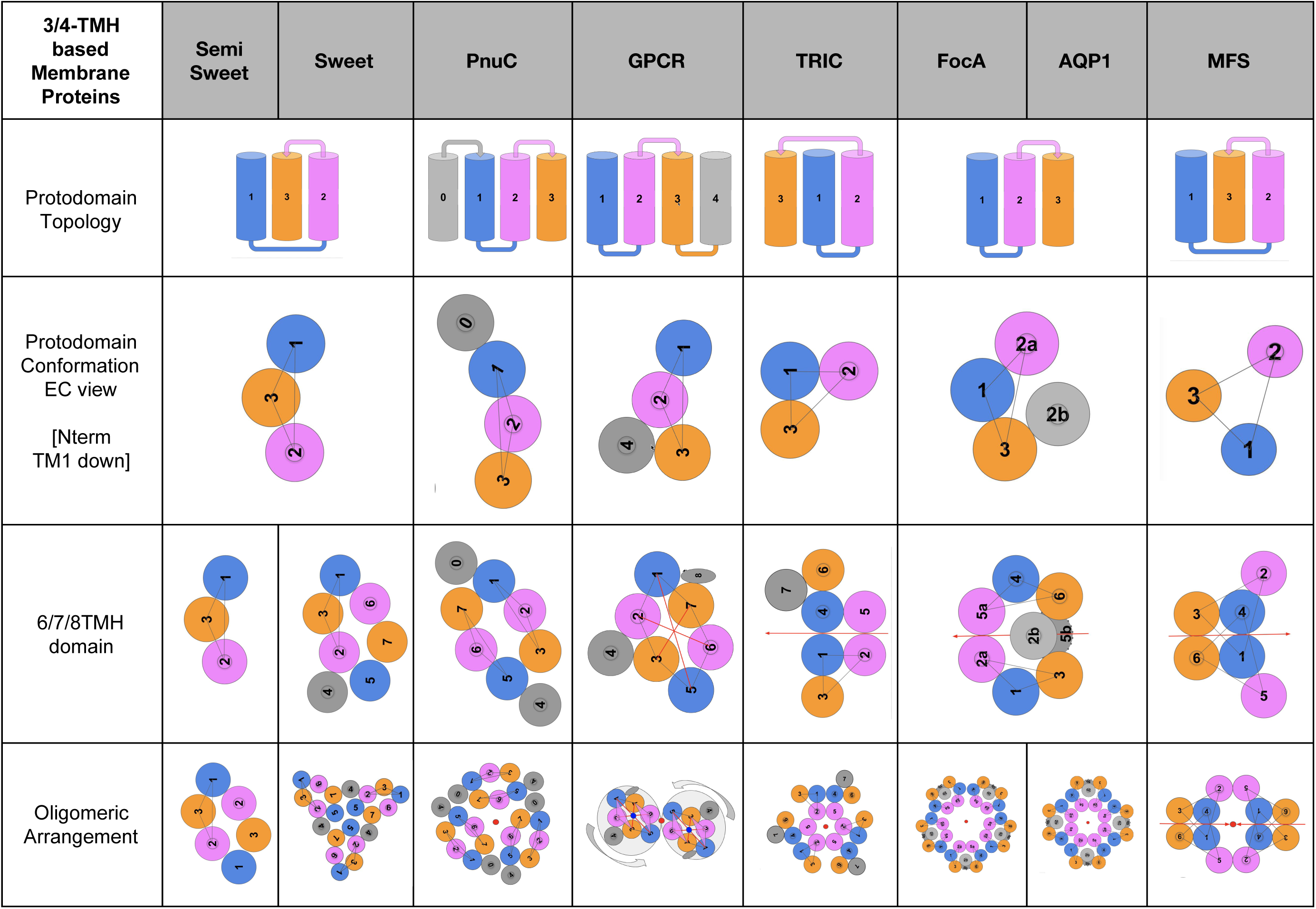
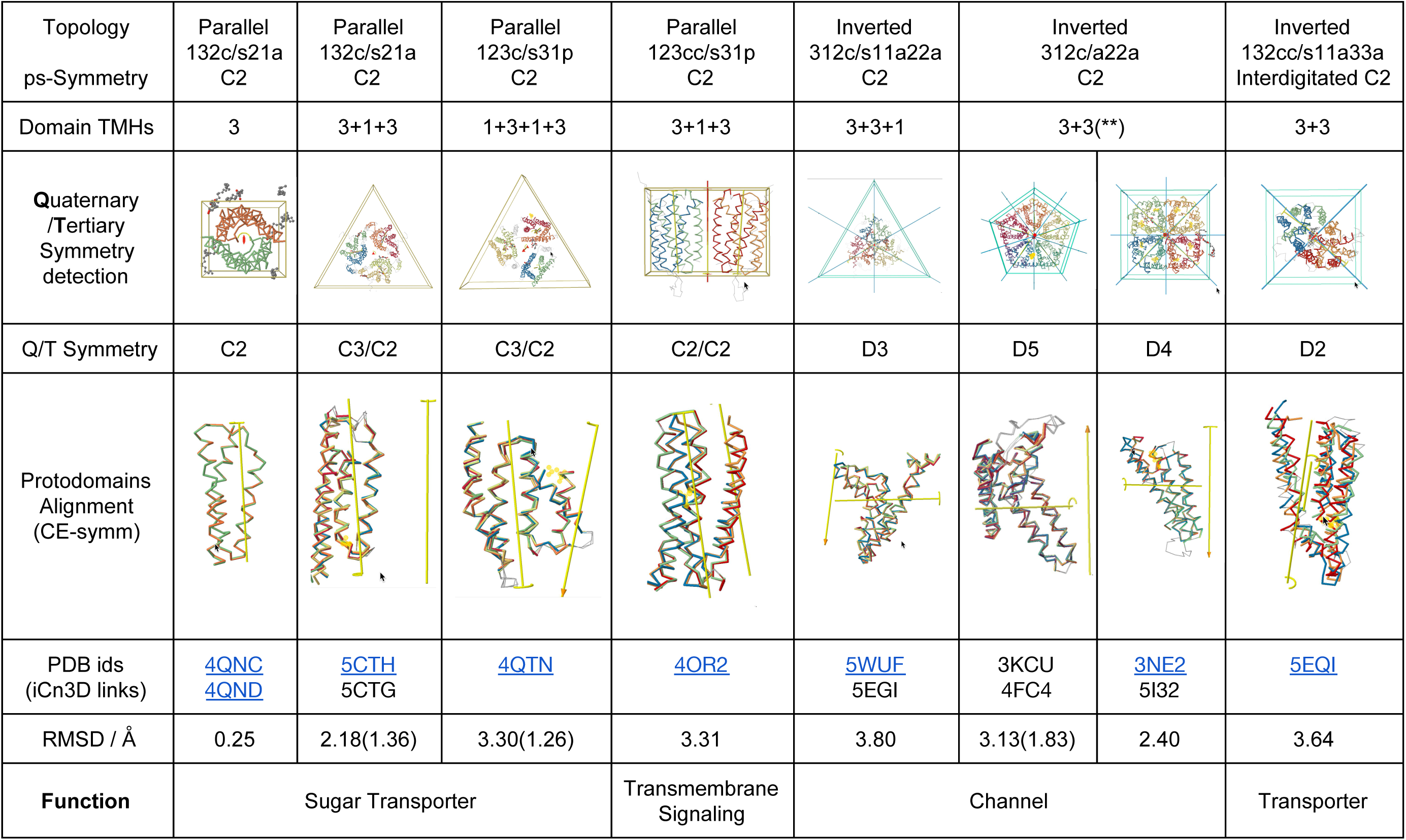
Pseudo symmetric domains formed from 3/4TMH prototomains. **Topology notation** in the example of Sweet and semiSWEET **132c/s21a** : **order 132** 3 helix bundle (1 down/2 up/3 down), **c** clockwise, **s21a** = **s symmetric contacts 21** [TM2(proto1)-TM1(proto2) and vice versa, i.e. TM2-TM5 and TM6-TM1 in SWEET], **a antiparallel helix contacts**.

It is important to note upfront, as we shall see, that GPCR structure based protodomain sequence alignment does not always show a high sequence conservation level in most cases to propose a conclusive duplication-fusion model. We use a parallel analysis of protodomains’ structural conservation within a wider set of 6/7/8-TMH pseudo-symmetric protein families, composed of 3/4-TMH protodomains, to identify generalizable evolutionary patterns. It leads to **guiding principles of protodomain assembly and to envision the role of conformational plasticity in fold formation and functional divergence.** It also provides a structural and mechanistic framework for the evolutionary deconstruction of current pseudo-symmetrical transmembrane helical (TMH) proteins.

### Protodomain Hypothesis in 7TMH Proteins

Structurally, nothing looks more like an individual transmembrane helix than another one. Some do possess breaks and tilts and can be perceived as unique, especially if structural alignments match them with a very low RMSD (see Methods for the definition; low RMSD values translate into higher structural homology).

The first step up in complexity is the smallest “supersecondary” helical structure we can envision: a Helix-Turn-Helix (2TMH) motif. In fact, at that level, we can already start seeing these 2TMH elements combining symmetrically to provide domains/folds, such as a 4-transmembrane helix (4TMH) bundle through intragenic duplication. The hemerythrin fold is a good example (Hendrickson and Ward, 1977). But in reality almost any 4-helix bundle (4TMH) can be seen as a symmetrically organized (C2 symmetry) duplicated Helix-turn Helix 2TMH “protodomain”. A pure geometrical analysis will in many cases show an even higher D2 symmetry for antiparallel (up/down) bundles. The Helix-Turn-Helix 2TMH motif can also lead to a 6TMH domain. A clear example of a C3 tertiary symmetry coming from a triplication of a 2TMH protodomain, has been observed in the case of the proton-gated urea channel (Strugatsky et al., 2013), with a parallel membrane topology, itself forming hexameric quaternary complexes (C6 symmetry).

The next step up in complexity for helical protodomains is a 3-helix motif (3TMH), or Triple-Helix-Bundle (THB), which upon intragenic duplication can lead to a 6 (or 7) helix bundle (6/7-TMH) and so on (Khan and Ghosh, 2015). **This will be “le fil rouge”, the unifying theme, in all membrane proteins examined in this study.**

Duplication (and shuffling) of domains, forming multi-domain proteins has been widely studied (Apic et al., 2001; Brenner et al., 1995; Kummerfeld and Teichmann, 2009). In fact, domain architecture of proteins is used to relate proteins through evolution (Geer et al., 2002; Scaiewicz and Levitt, 2018). Even therapeutic proteins now can be designed by architecting protein chains with domains, as immunotoxins (Alewine et al., 2015) or, now going even further, by architecting domains from functional subdomains, as in CARs (Chimeric Antigen Receptors)(Kochenderfer and Rosenberg, 2013). One could argue that linking domains together is a directed way of organizing a pseudo-quaternary arrangement of domains in proteins, usually asymmetrically within a chain. Quaternary organization itself has also been widely studied, and in that case, symmetry is pervasive (Goodsell and Olson, 2000; Levy et al., 2006; Rose et al., 2015). Compared to quaternary and pseudo-quaternary organization of proteins and protein complexes, pseudo-symmetry of tertiary structures, that can also be described as a pseudo-quaternary organization of domains themselves is not well documented, but work has started in that direction (Myers-Turnbull et al., 2014; Youkharibache, 2018 (in press)). A recent biophysical study on the ClC chloride transporter found that it is made up of two halves that fold independently as stable subunits, suggesting an evolutionary history of a stable protodomain that duplicated (Min et al., 2018). While the pseudo-symmetric organization of domains points to a clear mechanism of duplication and self assembly of protodomains, nothing at the moment can help point to an organized asymmetric assembly mechanism of diverse pre-folded subdomains in the creation of domains/folds (Alva et al., 2015; Youkharibache).

Sorting out structural pseudo-symmetry after intragenic duplication events in helical domains is not as straightforward as it may seem in the case of GPCRs, due to a lack of sequence similarity between protodomains. It is particularly true in the adaptation of the extracellular facing part of these receptors to recognize an astounding diversity of ligands as will be shown below. In order to infer a possible duplication-fusion event in an evolutionary process, we must rely on a sequence signature of some sort, where both structural/topological symmetry and sequence identity coincide.

In a parallel analysis of structurally and functionally different α-helical 6/7/8-TMH protein families, summarized in **Figure 1** (includes iCn3D (Jiyao Wang, Philippe Youkharibache, Dachuan Zhang, Chris Lanczycki, Lewis Geer, Renata Geer, Aron Marchler-Bauer, Tom Madej, Lon Phan, Minghong Ward, Shennan Lu, Gabriele Marchler, Yanli Wang, Steve Bryant, 2018 (Submitted for publication)) web links to 3D structures), we show the presence of symmetrically organized 3/4-TMH protodomains at the tertiary structure level (domains) and at the quaternary level (functional oligomers). **Supplementary Figures S1** through **S3** show the corresponding protodomain sequence alignments. Some protein families with similar functions, raise prospects of a divergent evolution at the sequence and structure levels from a common original 3TMH protodomain (called Triple Helix Bundle in the literature) or a 4TMH protodomain. These cases, however, need to be assessed critically at the sequence, the structure and the function levels simultaneously. The presence of 3/4-TMH protodomains in functionally different 6/7/8-TMH membrane proteins points to a parallel structural convergence of the gargantuan sequence space (sampled by the ancestors of these membrane proteins) into biophysically stable 3/4-TMH protodomain units (Min et al., 2018) in a number of conformations. So, the pseudo-symmetry of 6/7/8-TMH protein domains is also a geometrical property that results from a self complementary assembly of 3/4-TMH protodomains around an axis of symmetry normal to the membrane in parallel membrane topologies, or, conversely an axis of symmetry bisecting the membrane in “inverted” membrane protein topologies (as will be shown below), or both. It provides a natural functional (bi)directionality necessary for membrane-bound transporters and receptors. The conformational plasticity of 3/4-TMH protodomains seems to bring an element of structural diversity to pseudo-symmetric 6/7/8-TMH domains. Such symmetric pairing of protodomains is not limited to membrane proteins and has been observed in the β-strand globular proteins like the Ig domain and the ɑ-helical globular proteins like the 6-phosphogluconate dehydrogenase (**Supplementary Figures S4** and **S5**).

## Results and Discussion

### A. Protodomains Evidence and Evolution in 6/7/8-TMH proteins

The 6/7/8-TMH protein families analyzed in this study are presented in **Figure 1** in parallel. (Associated detailed figures are available in **Supplemental Figures S6 through S11**). They represent a variety of folds that are formed by duplication of 3/4-helix protodomains. They present either a parallel (SWEET, PnuC, GPCR) or inverted (TRIC, Aquaporin, FocA, MFS) membrane topology (see following sections for details). They all possess one axis of symmetry at the domain (tertiary) level, either parallel to the membrane planes (bisecting the membrane) or perpendicular to the membrane planes, according to that topology, respectively. Hence, their tertiary structure symmetry is C2. They all possess a higher level of quaternary symmetry that ranges from C2 to C5, where the axis of symmetry is perpendicular to the membrane. The tertiary and quaternary axes of symmetry can be parallel, leading to two levels of cyclic symmetry. They can be orthogonal to each other in the case of inverted topologies (Duran and Meiler, 2013), and combine to give dihedral symmetry groups D2, D3, D4 and D5 in the examples selected. MFS represents a special case: the domain has an inverted topology and two domains are fused, presenting D2 symmetry at the tertiary level. It can, however, be considered as a pseudo-quaternary structure from a symmetry standpoint, similar to all other cases. It is unique in this set of proteins in having an interdigitated topology.

#### Pseudo-symmetric assembly of 3TMH protodomains in an inverted topology: TRIC, Aquaporin, FocA, and MFS proteins

##### Multi-level symmetries in TRIC, Aquaporin, and FocA

A duplicated 3-helix (3TMH) protodomain can form a 6-helical C2 pseudo-symmetric membrane protein (6TMH) domain with a symmetry axis parallel to the membrane planes going through its center (TRIC, Aquaporin, and FocA in **Figure 1**). This implies a very short linker (or, if long, an extra or intracellular loop according to where the N terminus of second protodomain fuses with the C-terminus of the first protodomain). Two symmetrically related protodomains form, in that case, what is called an “inverted topology” in membrane protein terminology (Rapp et al., 2007).

Structural analysis of pseudo-symmetric TRIC domains, forming larger trimers, demonstrates the case of a 3-helix protodomain duplicated and linked through a short linker, hence with its axis of symmetry in the center of the membrane and parallel to the surface planes (Kasuya et al., 2016; Su et al., 2017). Let us note that even with the short linker, in a two inverted 3TMH protodomains symmetric arrangement as in the TRIC family architecture, the resulting 6TMH domain topology is, in fact, complemented by a 7th TMH at the C terminus (Su et al., 2017) resulting in a 7TMH protein domain.

In the parallel structural analysis of TRICs, Aquaporins and FocA, it has been noted(Su et al., 2017) that although *the topologies are similar*, their structural overlaps are highly divergent, and “if there is any evolutionary relationship among these inverted *repeat structures of different topologies*, it is lost in the sequence record”. This is consistent with our observation on numerous other sets of protein domains with 3TMH duplicated protodomains, where protodomains are highly idiosyncratic. In a nutshell, these domains show a very different 3-helix protodomain geometrical arrangement (conformation) followed by duplication with, in this case, an inverted (membrane) topology architecture (Rapp et al., 2007). It tends to demonstrate that the topological -and- geometrical (conformational) idiosyncrasy of a pseudo-symmetric domain is already built-in at the protodomain level.

Let us note in passing that the word topology, especially in membrane proteins, is used in different contexts with a different meaning, it can refer to the (tertiary) domain or protodomain, it can also relate to the membrane environment with a parallel vs inverted membrane topology, It also often refers, as in 3TMH, to the order of the transmembrane helices in the domain or protodomain, and it may also be sometimes equivocal if one considers diverse conformations. We attempt to avoid an equivocal use of the word by referring to the TMH order in the fold or protofold, to its clockwise vs. counterclockwise circular organization (**Figure 4.A**), and to a particular conformation (see **Figure 1** and **Figure 4** for definitions).

Beyond tertiary symmetry, TRIC, AQP1 domains and FocA form quaternary oligomers: trimers, tetramers and pentamers respectively. It is quite common to observe two levels (or more) of symmetry in protein complexes, whether in membrane or globular proteins. While we have not done a systematic analysis of co-existence of tertiary and quaternary levels of symmetry, it seems to be a common pattern, almost a given, that if a domain is pseudo-symmetric its propensity to form quaternary homo (or hetero) oligomers is high. One could eventually infer from this that GPCRs can probably form oligomers, as we shall see below.

Another case is Aquaporin and FocA, that share the exact same fold, with the same pattern of internal duplication of highly idiosyncratic protodomains, yet they seem to have lost a common sequence signature that can be detected (Theobald and Miller, 2010; Wang et al., 2009), leaving the door open to hypothesize a convergent evolution scenario. The structural idiosyncrasy, the uniqueness of that fold, especially at the protodomain level tells us that FocA and AQP1 should be related by divergent evolution. Effectively, if one performs a multiple protodomain alignment on both families, a sequence/structure pattern, or common signature seems to emerge on TM3/TM6 (see **Supplementary Figures S2 and S10** for details).

##### MFS: more than inverted

MFS is a very interesting fold, as it is hierarchical: a 6TMH domain is formed by duplication of 3TMH protodomains with an inverted membrane topology, followed by a domain duplication tying together two domains that dimerize through a pseudo-symmetric interface, hence a well integrated tertiary complex of dihedral symmetry (see **Figure 2.A**) emerges with 12TMH. The domain level assembly is a good example of a pseudo-quaternary association of domains, that are tethered together by a (quite long) covalent linker. The assembly is what would be expected from independent domains.

**Figure 2.**
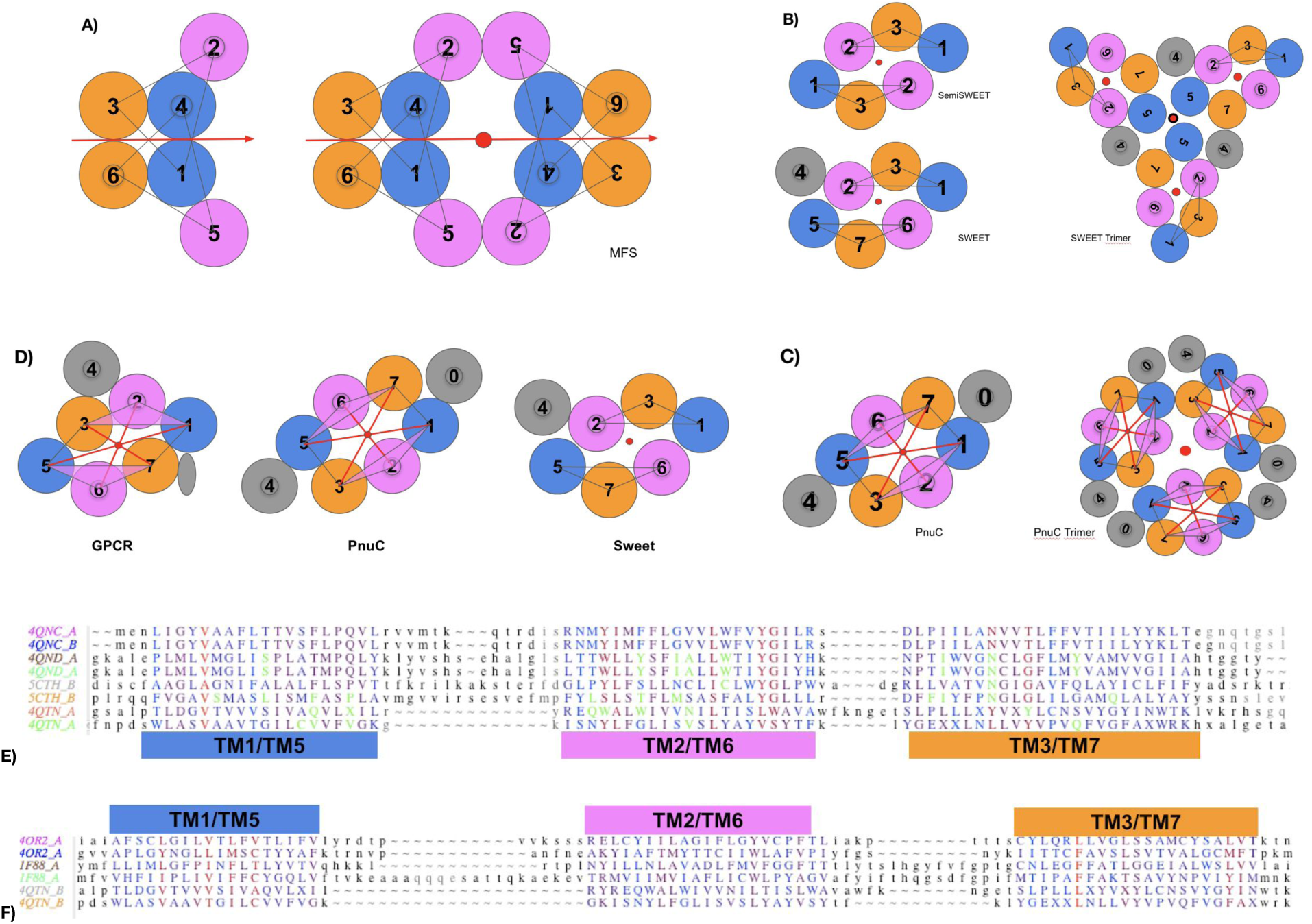
MFS, SemiSWEET, SWEET, PnuC and GPCRs. **A) MFS** - Left: **Inverted Interdigitated Protodomains forming a C2 symmetric domain**. It possesses two locally symmetric interfaces TM1-TM1 and TM3-TM3 (s11a/s33a). Right: **C2 symmetric domain packing** through a double (symmetry related) TM2-TM2 interface (2*s22a) formed by the second helix of each of the 4 protodomains. The 4-protodomains MFS protein has an overall D2 symmetry. (PDB: <u>5EQI</u>) since the internal symmetry axes of each domain colinearize and are orthogonal to the central axis (perpendicular to the membrane planes). **B) - SemiSWEET and SWEET** (left): a 6TMH dimer and a 7TMH domain respectively formed from 3TMH monomers/prototomains respectively. Sweet trimer (right) **C) PnuC 7/8TMH domain** (left; see text) **and trimer (right)**. **D) GPCR vs. PnuC vs. SWEET -** Comparison of topologies**. E) Structure based sequence alignments. SemiSWEET/SWEET/PnuC -** Structure based sequence alignment of 3TMH from SemiSWEET dimer (<u>4QNC</u>; <u>4QND</u>), SWEET (<u>5CTH</u>). PnuC (<u>4QTN</u>). The RMSD Deviation between two SemiSWEET monomers (<u>4QNC</u>) in the dimer are 0.17 Å, to another dimer binding a ligand (<u>4QND</u>) 1.29 Å, to SWEET protodomains (<u>5CTH</u>) 1.74 Å respectively. **Structural alignment** matching TMHs of PnuC (<u>4QTN</u>) with an **RMSD of 6.5 Å (alignment))**. PnuC is clearly not a structural homolog at the protodomain level. However from that very simple schematic representation, one can see that TMH167 and TM523 match the SWEET/SemiSWEET protodomain. Indeed if we ignore TM1 and TM5 the RMSD goes down to 1.71 Å and 2.47 Å respectively. For PnuC helices 167 vs SemiSWEET 3TMH (4QNC) the RMSD is 1.87 Å (see text for discussion). Ligand binding residues are in green, conserved/similar residues are in red. **F) GPCR vs. PnuC - Structure based sequence alignment** of 3TMH from a Class C GPCR (<u>4OR2</u>) vs. a class A GPCR (Rhodopsin) (<u>1F88</u>) vs. PnuC (<u>4QTN</u>) [see text] RMSD 3.63, 2.12, 3.29, 3.29, 3.34 Å respectively. However if one excludes TM2 from the structural alignment the RMSD is now 2.09, 1.17, 2.77, 2.61, 2.79 Å respectively.

The two domains, consisting of four protodomains, exhibit an overall D2 symmetry, combining two orthogonal axes of symmetry. One parallel and one perpendicular to the membrane planes. The first one is the domain level C2 symmetry axis relating 2 protodomains within a domain (**Figure 2.A** - left panel). Two domains assemble in bringing their C2 axis of symmetry as collinear, and are related by a second C2 symmetry axis perpendicular to the first one (**Figure 2.A** - right panel).

This collinearization of internal C2 symmetry axes of two domains is a typical dimerization with quaternary C2 symmetry, as in, for example, Immunoglobulins VH-VL domains (Youkharibache, 2018 (in press)), or a CD8 homo-or hetero-dimer (Youkharibache, 2018 (in press)). In fact, the inverted topology leading to an axis of symmetry in the middle of the protein, orthogonal to the longitudinal axis, is the same protodomain inversion that an immunoglobulin fold exhibits (see **Figure S4** on Ig domains symmetry). Membrane dimensions are such that this axis is a membrane-bisecting axis(Forrest, 2015), i.e. it runs in the middle of the membrane parallel to the membrane planes, depicted in the plane of the paper in our Figures (see **Figure 2.A** for MFS). In addition, the longitudinal axis of symmetry, normal to the membrane planes (the plane of the paper in our Figures, depicted as a red dot), corresponds to the communication axis (channel/transporter function) of these proteins.

To form a 6TMH domain, the two 3TMH protodomains in MFS are not just inverted, they are interdigitated. The 3 helices are not packed together, they exhibit a wide spacing between contiguous helices (Forrest, 2015) that precisely allows them to interdigitate with an image protodomain, forming a packed domain (see **Figure 1** and **Figure 2.A**). Such a protodomain, by construction or design, exhibits conformational flexibility. This flexibility in return enables the interdigitation of the helices in domain folding. The SWEET protein in contrast, presents a case with tightly packed protodomains that should form independent folding units, as we shall see next. We will come back to MFS after the sections on conformational plasticity and folding and see that the hierarchical assembly of this fold extends down to the secondary structure level (**Figure 2.A**).

#### Pseudo-symmetric assembly of 3TMH or 4TMH protodomains in a parallel topology: SWEET, PnuC, and GPCR Proteins

##### Multi-level symmetries in SWEET and PnuC

A long linker between 3TMH protodomains, long enough to form a membrane helix, enables the formation of pseudo-symmetric 7TMH proteins with the symmetry axis orthogonal to the membrane planes. The best example is the 7TMH SWEET protein. Its bacterial homolog, called appropriately SemiSWEET, is a 3-helix monomer that dimerizes to form a 6-helix quaternary structure binding a sugar in its central cavity lying on the dimer symmetry axis (**Figure 2.B**, also see **Figure 2.E** for corresponding structure-based sequence alignments). The arrangement is strictly conserved in the eukaryotic 7TMH SWEET domain, which is a pseudo-symmetric domain with its axis of symmetry, sugar binding, and local sequence patterns conserved. This provides evidence for duplication of two 3TMH protodomains, absolutely equivalent to the bacterial 3TMH SemiSWEET domain, to form the eukaryotic 7TMH SWEET membrane protein (**Figures 2.B** and **2.E**). In this case : 2*3TMH=6TMH+1TMH Linker.

The formation of helical linker is somewhat puzzling, however. In a simple duplication process of a 3 helix protodomain, one would have expected an inverted topology for the 2 protodomains, as is most often the case in membrane proteins. The most puzzling thing in this scenario is the linker forming a 4th helix. Another possibility is the duplication of a 4TMH protodomain. Yet the relationship to the bacterial SemiSWEET should then be questioned. The homology in the 3 protein dimensions: sequence, structure, and function seems to be a sign of duplication-fusion but with a linker allowing a parallel symmetric membrane topology between the two protodomains. In this case the generation of a central linker TMH would be a sign of a constraint to conserve function, *by inverting an “inverted topology”.* Inversion scenarios and comparisons of 3TMH protodomains have been reviewed extensively (Feng and Frommer, 2016; Jaehme et al., 2015; Xu et al., 2014). More recently it has been proposed from phylogenetic analyses that the eukaryotic SWEET protodomains may not have evolved by tandem duplication of an open reading frame, but rather originated by fusion between an archaeal and a bacterial SemiSWEET. This would potentially explain the asymmetry of eukaryotic SWEETs (Hu et al., 2016). In that study the 4th helix linker is ironically termed an “inversion linker” by the authors, since it inverts the repeat that would otherwise be inverted in the membrane! Regardless of the phylogenetic origin scenario, whether homo-fusion or hetero-fusion, how the TM4 linker was inserted or evolved remains elusive.

The vitamin B3 transporter PnuC Proteins represent another very interesting case of a 7/8-TMH that exhibit C2 pseudo-symmetry with evidence of a 3/4-TMH protodomain evidence (**Figure 2.E**) with a parallel topology. It also forms a trimeric quaternary structure (see **Figure 1** and **Figure 2.C**). As a family, PnuC is described as a 7TMH (e.g., Uniprot B8F8B8). The particular PnuC domain structure (PDB: 4QTN - Uniprot D2ZZC1) has 8 TMHs, that we use as an example of a possible 4TMH protodomain duplication (**Figure 2.C**). It is clear, however, that the 4th helix does not match, structurally, with its proposed symmetric counterpart in a decisive manner. The RMSD of TM0123 vs. TM5678 is 3.09 Å while if we reduce the protodomain to a 3TMH, then TM123 vs. TM567 RMSD is 1.26 Å (see **Figure 1**).

From the very simple schematic representations used in **Figure 2.B**, one can align TM1-TM6-TM7 and TM5-TM2-TM3 to SWEET/SemiSWEET 3TMH with a high structural homology (low RMSD). This is equivalent to a symmetric structural swap of TM1/TM5 as proposed (Jaehme et al., 2015), as sugar transporters PnuC and SWEET have effectively been proposed to be evolutionary related. PnuC is seen as a “full length” SWEET homolog (Feng and Frommer, 2016; Jaehme et al., 2014, 2015, 2016). This is in line with a survey(Murzin, 1998) on divergent structural folds, where the author notes: “… A simple way of altering a protein fold without a big destabilization is to change its topology while maintaining its architecture. […] This can be done by the internal swapping of similar helices and strands or by reversing the direction of some of its secondary structures….” PnuC and Sweet could effectively be considered topological isomers, or “topoisomers”(Murzin, 1998) having a similar structure despite a different topology, and having a similar function. Another explanation might be found in envisioning a permutation at the gene level between SWEET and PnuC involving a segment covering 4 helices TM2345. Permutations of secondary structure elements, commonly seen as circular permutations (CPs), conserve 3D structure, i.e. the order of secondary structure elements does not affect the folded structure (Viguera et al., 1995). To our knowledge however, CPs have not yet been observed in membrane proteins, however, circular and non-circular permutations have been engineered to show structural and functional resilience of alternate topologies (Mackin et al., 2014).

The mechanism by which such fold changes might occur is unknown. We discuss below a possible general mechanism of “conformational evolution” in the context of pseudo-symmetric folds where we will look at some similarities and differences between these transporters extending to the GPCR fold that has also been considered as a topological isomer by some authors (Saier, 2016; Yee et al., 2013).

##### Pseudo-symmetry evidence in the GPCR domain

Regardless of the evolution leading to the 7TMH GPCR fold, their geometry, the spatial arrangement of its 7 helices is such that it can be considered as formed by two 3TMH protodomains (TM123 and TM567) related by a C2 symmetry, with a TM4 “linker” (**Figure 3.A**), as in SWEET proteins. GPCRs pose a very challenging problem from an evolutionary standpoint. As new structures are coming out at an increasing pace (Ghosh et al., 2015; Thal et al., 2018), we can start to analyze these observed protodomain idiosyncrasies in the various GPCR classes, in particular GPCR class A and class C, and more recently class B and F, but also some bacterial 7TMH proteins that may be related.

**Figure 3.**
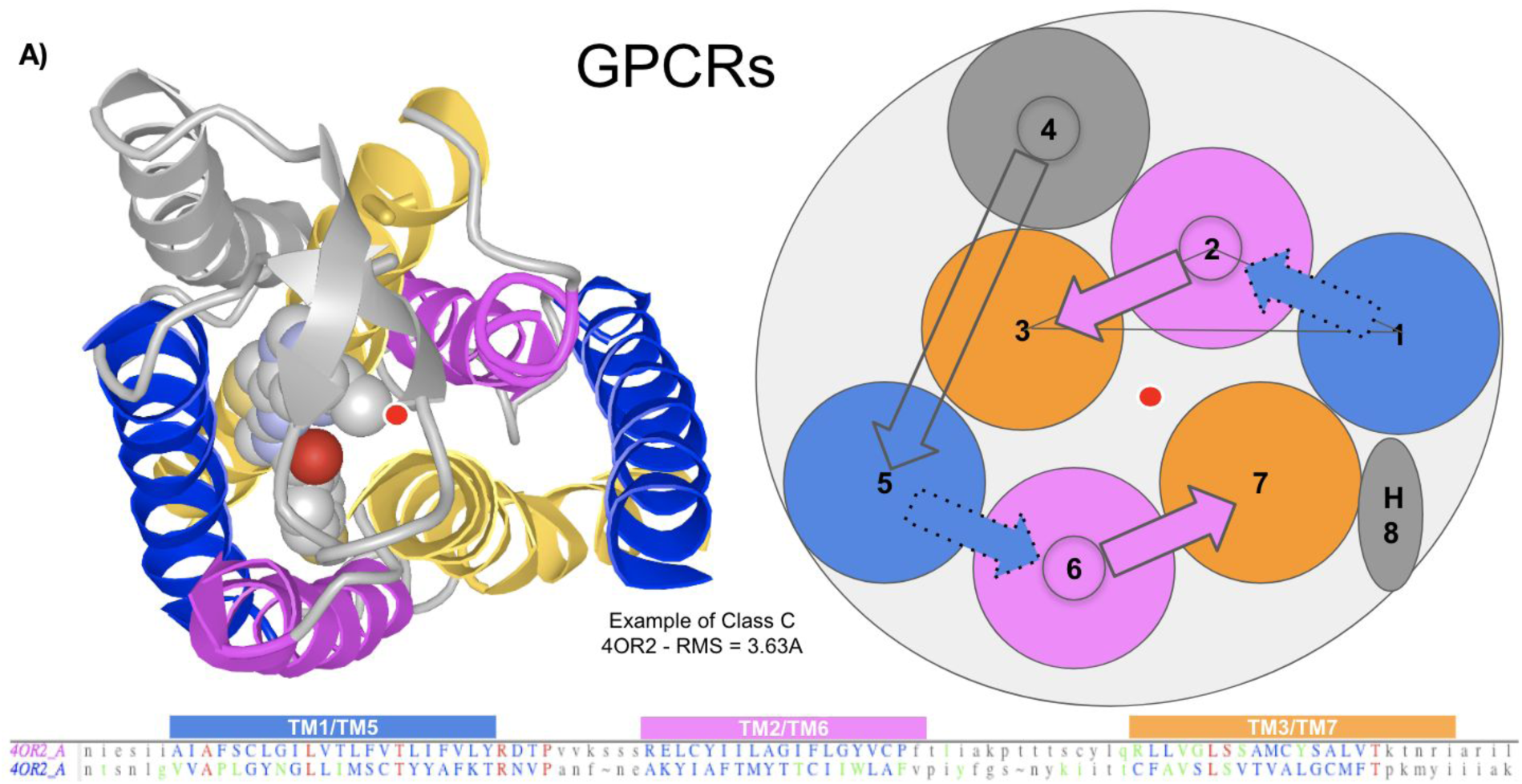
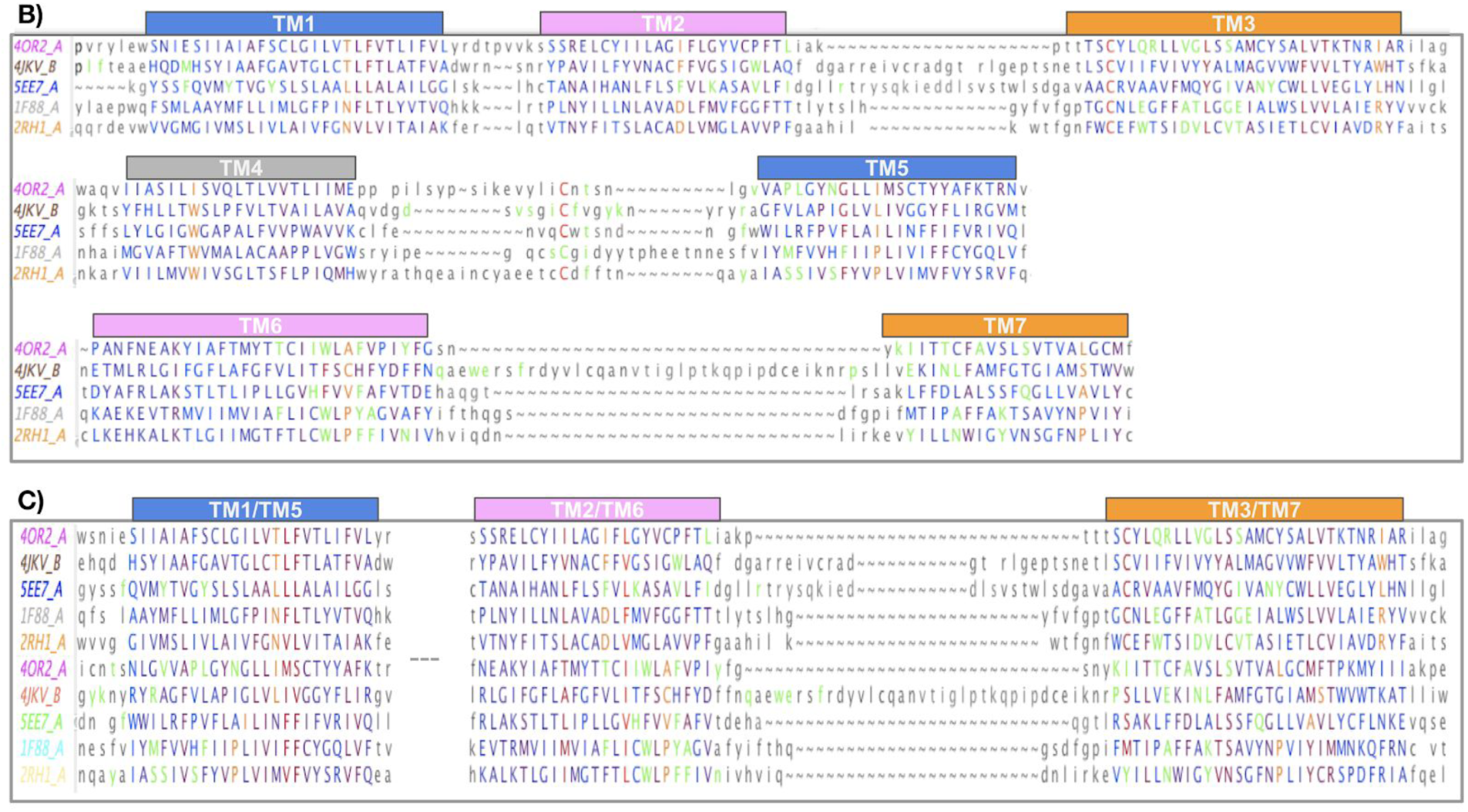
GPCR Class A, B, C, F structure alignments. **A) Structure of a GPCR** with a 3D representation and a 2D representation, seen from the extracellular side. **B) Multiple Domain Alignment** for Class C, A, C and F (RMSD relative to 4OR2.A: 4JKV.A: 3.06 Å, 5EE7.A: 2.91 Å, 1F88: 3.21 Å, 2RH1: 3.09 Å. **C) Multiple Protodomains Alignment**: TMH-123 vs. TMH-567 - RMSD relative to the first protodomain (4OR2.A-1) used as a reference structure in the alignment: 3.63 Å (4OR2.A-2), 2.12 Å (1F88.A-1), 3.29 Å (1F88.A-2), 1.73 Å (5EE7.A-1), 2.77 Å (5EE7.A-2), 1.93 Å (4JKV.A-1), 2.55 Å (4JKV.A-2), 1.79 Å (2RH1.A-1), 3.59 Å (2RH1.A-2) respectively for Class C (<u>4OR2</u>, human metabotropic glutamate receptor 1: GRM1), Class A (<u>1F88</u>, bovine rhodopsin: OPSD and <u>2RH1</u>, human β2 adrenergic receptor: ADRB2), Class B (<u>5EE7</u>, human glucagon receptor: GCGR), Class F (<u>4JKV</u>, human smoothened receptor: SMO). Similarity scale from blue to red (most similar). In green: ligand proximal residues (at less than 4 Å distance) when a ligand is present in the crystal structure. In orange the most conserved residue positions in Class A (1.50 N, 2.50 D, 3.50 R, 4.50 W not shown, 5.50 P, 6.50 P and 7.50 P) and their counterparts in other classes.

We originally came across GPCRs 3TMH pseudo-symmetry through a visual inspection and an interactive structural alignment of the first Rhodopsin structure that was crystallized (1F88 (Li et al., 2004)). Our focus has been for a long time in trying to understand protein folds, if not folding, that are formed of protodomains, i.e. repeated supersecondary structures related by structural symmetry, packing together to form protein domains. GPCRs and their protodomains are difficult to align within a very small RMSD, as for most helical structures, since helices tend to shift along their helix axis and/or move sideways (see Methods). In addition, the TM567 protodomain has conserved Prolines (classical helix breakers) in each of the TMs that cause kinks in those helices making it harder to align the TM567 protodomain to the TM123 protodomain. This translates in difficulties to accurately delineate structural protodomains, and there may be a mismatch between optimal structural alignment vs. optimal sequence conservation alignment. In the case of Rhodopsin (1F88) and a handful of class A GPCR structures we can get to a 3TMH protodomain delineation and alignments computationally, albeit with stringent alignment criteria (see Methods). In the case of known class C structures (<u>4OR2</u>/<u>5CGD</u>) the software can identify both tertiary (TM123 and TM567) and quaternary pseudo-symmetry, hence two levels of C2 symmetry, and delineate protodomains accurately (see **Figure 1** and **Figure 3**). This is quite satisfactory, since Class C GPCRs have been shown to be the most ancient GPCR class through phylogenetic analysis (Cvicek et al., 2016; Krishnan et al., 2012) and additionally form functionally obligatory homodimers.

**Figure 3** summarizes multiple structure-based sequence alignments at the domain (**Figure 3.B**) and the protodomain (**Figure 3.C**) levels, across known structure representatives of all classes of vertebrate GPCRs: A,B,C, and F (Fredriksson et al., 2003; Lagerström and Schiöth, 2008). Pairwise structure based protodomain alignments, where sequence matching patterns are easier to see, are available as **Figure S2** for a larger number of structures.

##### GPCRs vs. PnuC: Are they related?

As we have seen earlier, a question comes back in almost any discussion that deals with pseudo-symmetric membrane proteins formed by duplications of 3TMH protodomains. Are these domains formed by convergent or divergent evolution through duplication-fusion of Triple Helix Bundles (3TMH protodomains)? In the first case of TRIC vs. AQP1 or FocA, it has been noted that although the topologies are similar, their structural overlaps are highly different (Su et al., 2017). In the case of SWEET vs. PnuC (**Figures 2.B** and **2.C**), it has been postulated that PnuC (PDB: <u>4QTN</u>) may be a “full length SWEET Homolog” since they both share a sugar binding function. But, are they really true homologs? i.e., are they derived from a common ancestor, the same *protosequence*? A domain swap has been hypothesized as a possible conformational transition at the protodomain level to reconcile different topologies of the three helices (Jaehme et al., 2014, 2015, 2016). Another study goes further and groups SWEETs, PnuC, and GPCRs into a “transporter–opsin–GPCR” (TOG) superfamily (Yee et al., 2013).

If we compare the topology of PnuC vs GPCR protodomains, they share the TM1-TM2-TM3 helix order, yet PnuC presents a clockwise topology vs. a counterclockwise one for GPCRs at the protodomain as well as at the domain level (**Figures 2.D** and **2.F**). If there is any homology relation, it would imply a conformational change at the protodomain level (see later).

The comparison of protodomains of GPCRs vs. PnuC is quite instructive. It provides a critical evaluation of the homologies inferred at the sequence level, the structure level, and the function level. We have seen earlier the difference in protodomain topology which is very significant between PnuC and SWEET despite a similar function of sugar transport, raising a question on a possible common origin of these two structurally different domains. In that case, the sequence based structural alignment of the 3TMH protodomains has an RMSD of over 5 Å (weak structural homology). Here if we align GPCR vs. PnuC protodomains, they align within 3.5 Å, a number still in the range of RMSD values used to compare protodomains all along (see **Figures 2.D** and **2.F**). At the sequence level, the cross-match is about as inconclusive as any two GPCRs protodomains, hence we do not have much leeway to assess homology with this method. In fact, at the sequence level, one can sometimes get good matches between unrelated membrane proteins. All elements (sequence/structure/function) need to be considered together to make judgements about a common ancestor protein. When carefully looking at GPCR vs PnuC protodomains, we have the same 1-2-3 order for the 3TMH topology, they structurally align within 3.5 Å, but one positions the triangle apex (seen from the extracellular side) showing a clockwise protodomain topology in PnuC vs. a counterclockwise one in GPCRs, hence two different 7TMH domain topologies emerge, one clockwise and the other counterclockwise. This, on face value, raises the possibility of “conformational evolution”, where a possible sequence level homology may lead to different conformations of a protodomain and that could lead, through a duplication-fusion event, to different folds. However, for GPCRs vs PnuC, a weak structural homology in the absence of any sequence and functional similarity could be considered “parallel evolution”(Murzin, 1998).

### B. Evolution through Protodomains Duplication and Symmetric Assembly

#### Protodomains idiosyncrasy and symmetric assembly

**The question on evolution and homology** of various domains is a long standing one. For example, in the debate over divergent vs convergent evolution of Type I opsins like Bacteriorhodopsin or Sensory Rhodopsin II vs. Type II opsins like Rhodopsin (a prototypical Class A GPCR), at the sequence level a weak homology has been noticed for opsins type II between TM123 and TM567 (called ABC and EFG in that context) (Larusso et al., 2008; Taylor and Agarwal, 1993). The prevalent opinion is leaning towards the convergent evolution hypothesis. Nonetheless, some authors are strong proponents of a divergent evolution scenario (Devine et al., 2013; Larusso et al., 2008; Mackin et al., 2014; Taylor and Agarwal, 1993). Their question is rightly so: “*Given two transmembrane proteins with identical folds, yet no sequence similarity, how then could we distinguish convergence from homology?* It is effectively an unanswered question in molecular evolution (Doolittle, 1994; Murzin, 1998), a particularly acute one in the cases of pseudo-symmetric domains formed through duplication of triple helix bundles (THB, i.e. 3TMH).

Protodomains (protofolds) tend to be idiosyncratic supersecondary structures. In other words, they adopt a specific topology and conformation that is duplicated and that assembles in a complementary manner in forming a symmetric domain (fold). The *Triple helix bundles (THB, i.e. 3TMH protodomains) forming pseudo-symmetric 7TMH domains are effectively highly idiosyncratic. These repetitive supersecondary structures, or protofolds, are all different and do not align with each other through a rigid alignment.* Protodomains represent a signature of a pseudo-symmetric domain/fold. A question remains about the possibility of common *protosequences* among diverse folds that may be associated with a common function (Petrey et al., 2009), as in the case of SWEET vs. PnuC (see above).

#### Conformational Evolution?

When we consider pseudo-symmetric folds, each and every one of them is formed by distinct, idiosyncratic protodomains that exhibit a particular topology and conformation. This is especially true of all the 3TMH protodomains in this analysis. We consider that SWEET, PnuC and GPCR folds are different after careful review of protodomain sequence, topology and conformations, simultaneously. However, the recurrent discussions in the literature on possible homology of diverse pseudo-symmetric domains formed from 3TMH can be summarized as two questions:

1. Do various conformational changes within protodomains, possibly due to sequence changes and/or between protodomains due to structural constraints, enable the **formation of structurally different folds, while being sequence homologs**?
2. Conversely, does conformational convergence of unrelated sequences in protofolds followed by duplication-fusion enable **structurally similar folds**, **while not being sequence-homologs originally**?

This lead us to address the structural relation between symmetric folds of different topologies, formed by different protodomains; folds that can eventually be considered topological isomers, “topoisomers” (Murzin, 1998). This has been proposed in the case of Pnuc vs SWEET (Jaehme et al., 2015). From the structural analysis of existing topological isomers, Murzin observes that: “The close structure conversion of one protein topoisomer into another would require at least partial unfolding” but also that “the folding of different topoisomers of a protein chain is yet to be observed”(Murzin, 1998).

We consider now the conceptual mechanism that we will refer to as “conformational evolution” of 3TMH protodomains that would permit **the formation of *structurally similar pseudo-symmetric 7TMH folds from unrelated sequences*** *through:*

- *conformationally similar 3TMH protodomains with different, unrelated sequences leading to convergent 7TMH folds through a fused assembly process (****Figure 4.B****), or*
- *convergence of 7TMH folds through conformational change of fused 3TMH under a structural constraint that a fusion linker (TM4) may provide (****Figure 4.C***).

*Or, conversely, **the creation of structurally different folds, even from related sequences**:*

- *conformational changes of 3TMH protodomains for related sequences, which then, upon duplication/assembly would lead to divergent structural folds.*

**Figure 4.**
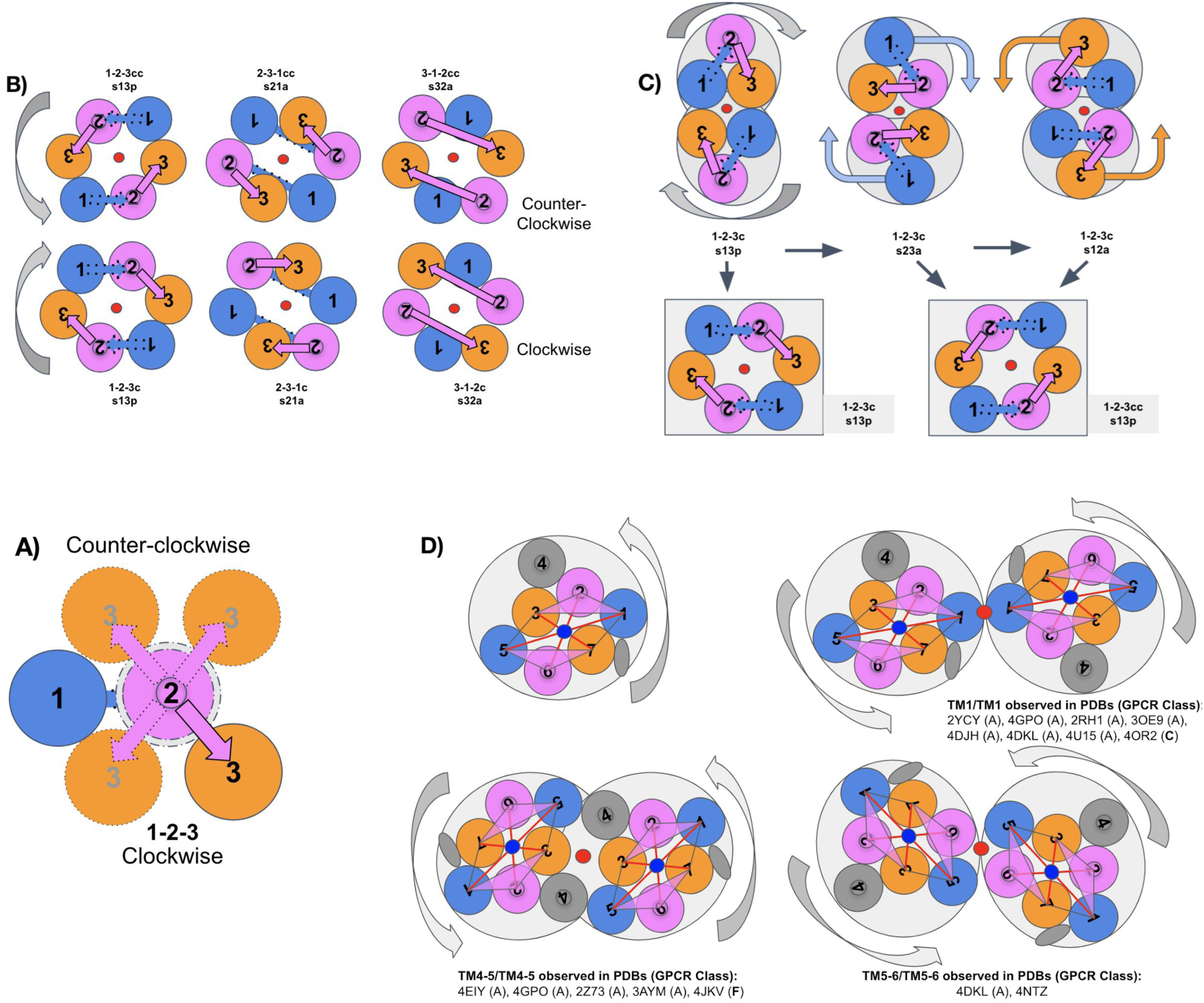
Self-Complementarity. **A)** A **3TMH** sampling clockwise and counterclockwise topologies/conformations, **B)** Duplication and **symmetric assembly of self-complementary conformations** [rigid or conformational selection model] of 3TMH protodomains with a 123 clockwise vs. a 123 counterclockwise topology, to form a 7TMH parallel membrane topology assuming a middle TMH linker (TM4) [not displayed for clarity). In the 123 topology where TM2 is central, TM1 and TM3 fromthe 2 proto domains are in contact in a symmetric manner (denoted s13/31), and so on: the 2-3-1 topology (s21/12) and the 3-1-2 (s32/23). We therefore enumerate 3 possible clockwise topologies and 3 counterclockwise. Naturally, ignoring loop connections between TMHs, any 123, 231 or 312 order could be matched with a rotation around the symmetry axis, and similarly for the counterclockwise set. So effectively, structural similarity can also be observed between folds when topology is ignored, as in the case of PnuC vs. SWEET (see text). In the 123 topology where TM2 is central, TM1 and TM3 from the 2 proto domains are in contact in a symmetric manner (denoted s13/31), and so on: the 2-3-1 topology (s21/12) and the 3-1-2 (s32/23). We therefore enumerate 3 possible clockwise topologies and 3 counterclockwise. Naturally, ignoring loop connections between TMHs, any 123, 231 or 312 order could be matched with a rotation around the symmetry axis, and similarly for the counterclockwise set. So effectively, structural similarity can also be observed between folds when topology is ignored, as in the case of PnuC vs. SWEET (see text). **C)** Duplication and **local symmetric assembly of 3TMH protodomains** in the case of a 123 clockwise topology [concerted/induced conformational model]. A rotating protodomain symmetric assembly can sample a number of pseudo-dimers. A conformational change preserving symmetry to form a 7TMH (parallel membrane topology assuming a middle TMH linker (TM4 - not shown) forming either a clockwise or counterclockwise 7TMH domain. Interestingly the two rotated protodomains on the right lead to a counterclockwise domain-level topology. Hence B and C scenarios produce the same six domain-level topologies whether a protodomain symmetric assembly starts from a given stable self-complementary 3TMH conformation or possibly involves a conformational change after a local self complementarity symmetric assembly process. **D) Rotating 7TMH GPCR dimer** sampling various symmetric homodimers observed in Crystal structures (see paragraph on oligomerization of GPCRs).

The last process corresponds to the one suggested in the case of PnuC vs. SWEET (see above). The first process however seems relevant w.r.t discussions in the literature about Aquaporin vs FocA(Theobald and Miller, 2010), where no similarity seems decisively showing a divergent evolution scenario. However, this fold has a very peculiar central helix splitted as TM2a/b (and duplicated TM5a/b), which is so unique that we would suggest a common origin. In fact, as mentioned earlier, a careful multiple protodomain alignment can let a conserved sequence/structure pattern appear in TM3/TM6, although not in TM2/TM5a/b, common to both families (see AQP1/FocA section above, and **Supplementary Figures S2 and S10** for more details).

##### Conformational Variability of 3TMH bundles forming pseudo-symmetric 7TMH domains

If there is any evolutionary relation between two or more domains formed by 3TMH, while they are topologically and/or structurally different, a conformational transition between them can be envisioned. In essence that would mean that related sequences of 3TMH protodomains (proto-sequences) can lead to different folds if they can sample different conformations that can self assemble symmetrically to form 6/7/8-TMH folds. The following analysis looks at samples of conformations of a generic 3TMH bundle and envisions, for each conformation, a symmetric assembly. This would be, in essence, a homo-dimerization process of 3TMH, where a covalent linker acts as a variable constraint to assemble, as a conceptual mechanism to generate structural fold diversity from homologous sequences with variable conformations, i.e. a combinatorial fold creation. However, while structural diversity can be associated to a given sequence with such a mechanism, possibilities of generating different pseudo-symmetric folds with duplicated 3TMH are numbered (see **Figure 4**).

##### Coincidence of Lipid membrane dimensions and Protein domain axes of symmetry

Lipid membranes reduce the dimensionality of 3D space, from 3D to 21/2D if one wants to describe it in simple terms (Theobald and Miller, 2010), hence the particular connection between symmetry and membranes. Two protein symmetry axes correspond to the membrane dimensions, and are either parallel or orthogonal to the membrane planes. Besides functional requirements of establishing communication channels (used in a general sense) between the inside and outside of a cell along one (vertical) axis, the geometry and the medium of membranes create an environment where dimensionality and directionality is 2D: vertical (in/out or up/down) and lateral (diffusion). The third dimension (for folded objects) could allow either or both of two orthogonal movements:

- **A flip between IC and EC:** This is normally prevented by the positive charge rule (von Heijne, 1992). The flip has been envisioned as a possibility in spite of that rule as a possible mechanism for inverted membrane topologies (Bowie, 2013; Min et al., 2018). That would apply to inverted protodomains, if protodomains are folded independently and behave as rigid bodies. In some cases, the whole domain topology can be controlled by a change in the lipid composition (Bogdanov and Dowhan, 2012; Bogdanov et al., 2008; Dowhan and Bogdanov, 2009; Vitrac et al., 2015).
- **A relative rigid translation and/or rotation of proteins laterally within a membrane:** The rotation axis of proteins being limited to the protein vertical axis, orthogonal to the membrane planes. This may be the biggest impact of a reduced dimensionality of a lipid environment.

The mechanism of membrane protein folding is an open and challenging problem (Cymer et al., 2015). Beside a rigid flip or inversion of independently folded protodomain, chaperones have been envisioned (Bowie, 2013). The interconnection between symmetry and folding may help get some insight.

##### Self-complementarity of protodomain conformation(s) in symmetric self-assembly

It is possible, and quite likely, that 3TMH topologies may, for some of them, sample more than one conformation, and might simply exist as conformational ensembles. One conformation that exhibits **self-complementarity** may be “selected” from an ensemble to form a symmetric domain. Alternatively such a conformation may be induced during the folding of the domain, or the tethered assembly of protodomains, if each is folding autonomously.

We are therefore lead to ask: what process may induce or select a self-complementary conformation on two protodomains assembling symmetrically. These are similar to questions raised in discussions on conformational selection vs. induced fit (Boehr et al., 2009; Changeux and Edelstein, 2011; Koshland, 1959; Koshland et al., 1966). The specificity is in *assembling the same or similar* entities in a symmetric or pseudo-symmetric manner. It is an oligomerization process under constraint (covalent linker). Whether one mechanism or another is used in reality would have to be tested experimentally. The fact that we observe numerous pseudo-symmetric folds, well beyond 3TMH (see **Table 1**) in membrane as in globular proteins, which tends to show that pseudo-symmetric structures retain favorable energetics, something that has been discussed for quaternary symmetric structures (Goodsell and Olson, 2000; Levy and Teichmann, 2013; Monod et al., 1965; Wales, 1998; Wolynes, 1996). Symmetric tertiary structures are, in effect, pseudo-quaternary structures. The difference is caused by the linker which brings two or more protodomains together in (tertiary) domain folding.

Symmetric quaternary structures show that folded domains, for most, assemble rigidly, although some conformational changes may occur, with subdomain swaps between domains (Liu and Eisenberg, 2002). Symmetric tertiary domains may well have their protodomains folding independently and assembling symmetrically. Interestingly, a circular permutation in sequence, would change such a folding pathway but would still form the same fold overall, which would then still be structurally but not topologically symmetric. This has been demonstrated experimentally in the case of globular proteins (Viguera et al., 1995), and can be observed among known circularly permuted domains, [see for example domains 2 and 3 of the elongation factor Tu (EF-Tu)(Andersen et al., 2000); SCOP folds b.43 and b.44](Chandonia et al., 2017; Lo Conte et al., 2000). Folding into a symmetric fold, especially a pseudo-symmetric one, does not necessarily imply a symmetric folding pathway, nor a symmetric topology. As far as we know, circular permutations have not yet been observed in membrane proteins, whether that is the norm of simply a limitation of our current knowledge is in question. If there are no circularly permuted membrane proteins, that would be surprising, but if real, that would be of significance.

##### Envisioning folding/assembly of conformationally flexible Protodomains

Protodomains have, in most cases, a small number of secondary structure elements and are likely to exhibit conformational flexibility. This is most likely the case in 3TMH. A pseudo-symmetric protein domain folded state implies that protodomains adopt the same conformation. That conformation presents a self-associating property: it forms a symmetric interface. We will refer to a protodomain conformation and protodomains interfaces as “self-complementary”. We may envision a number of mechanisms to relate symmetric protodomain folding/assembly within domains:

1. **Co-assembly:** Protodomains exhibit **one and the same pre-formed self-complementary conformation** that self-assemble to form a symmetric domain (**a rigid docking/assembly model**). This is similar to a domain level (monomers) symmetric oligomerization.
2. **Co-folding:** One and the same conformation of protodomains that can self-assemble symmetrically is **selected** out of an ensemble of accessible conformations (**a conformational selection model**) to form a symmetric domain. The same conformation would have to be co-opted by protodomains.
3. **Co-chaperoning:** A concerted mimetic mechanism, where protodomains converge on a self-complementary, self-assembling conformation, or where a protodomain conformation serves as a template for the shaping of a matching protodomain (**a symmetrically induced conformation model**) to form a symmetric domain.

All three mechanisms give a protein folding pathway in line with the Levinthal paradox (Levinthal, 1968, 1969), where “protein folding is speeded and guided by the rapid formation of local interactions which then determine the further folding of the peptide. **This suggests local amino acid sequences which form stable interactions and serve as nucleation points in the folding process**”(Levinthal, 1969).

Symmetric quaternary oligomerization has been amply reviewed (Goodsell and Olson, 2000; Levy and Teichmann, 2013); and so has been conformational selection in molecular recognition (Boehr et al., 2009). The induced fit (Koshland, 1959; Koshland et al., 1966) and the MWC (Monod-Wyman-Changeux) model have been postulated long ago (Changeux, 2012; Monod et al., 1965), and the debate itself between various mechanisms is as old as the models themselves (Boehr et al., 2009; Changeux and Edelstein, 2011). In all fairness there is room for more than one mechanism. Interestingly, the MWC model states that a protein composed of subunits must maintain symmetry *during* conformational changes. There is an interesting parallel, but let us stress however that our discussion is about a symmetric assembly process itself, *the co-folding and/or assembly of the same with the same (or similar with similar).* Regardless of the folding and/or assembly process itself, in the end, a protodomain self-complementary conformation and symmetric assembly leads to a structurally symmetric domain that may be a low free energy state(Wolynes, 1996), or simply a metastable state (Levinthal, 1969).

##### Combinatorial sampling of 3TMH conformations and symmetric self-assembly

A 3TMH bundle can adopt a variety of “topologies”. This word is usually used loosely to mean three sequential helices down-up-down (or vice versa) in a specific geometric order, for example 1-**2**-3, 2-**3**-1, or 3-**1**-2 as in **Figures 4.B and 4.C**. We refer to the central TMH as the apex of the triangle formed when looking from the extracellular side of the membrane protein. We can also specify if the triangle is clockwise (c) or counterclockwise (cc). This circularity will be distinctive at the domain level: GPCR domain is counterclockwise, while PnuC is clockwise (see **Figure 2.D**).

I. If we enumerate **conformations of 3TMH** (in parallel membrane topology to keep it focused), we find 6 conformations, taking circular directionality into account. If any such conformation for a given sequence is self-complementary, it can form a corresponding symmetric domain with a symmetric interface of non-apex helices. For example 1-2-3cc forms a double (symmetric) interface s31p (see **Figure 4** legend for definition), i.e. s35 and s71 in the case of GPCRs (where TMH5∼=TMH1 and TMH7∼=TMH3; see **Figure 1** and **Figure 3**). Note that the helices interfacing protodomains in this GPCRs topology are parallel (p). Naturally symmetry is not exact in GPCRs, helices are not exactly perpendicular to the membrane planes and TM4 modifies partly one interface. This is a simple schematic dimeric model, as in the case of SemiSWEET with a 1-3-2cc topology and a symmetric (double), antiparallel, s21a helical interface. In SWEET, it translates into a pseudo-symmetric interface s25a=s61a with a domain level sequential numbering, where TMH5∼=TMH1 and TMH6∼=TMH2 (see **Figure 1** and **Figure 2.B** where 1=blue, 2=magenta, 3=orange).
II. If we now take rotating compact protodomains (**Figure 4.C**) for a particular topology 1-2-3c, we can form a number of symmetric interfaces: s13p but also s12a and s23a. The latter two do not form a closed domain as such, but if we envision a concerted conformational change between protodomains preserving symmetry (could be seen as a helix swap), we do end up with a closed domain, the same for both which are now counterclockwise. Hence, overall this mechanism can form a clockwise and counterclockwise domain from a given 123 topology and, overall, six closed 7TMH domains, when considering 231 and 312 topologies, the same as in I. In this schematic example, whether a protodomain conformation is selected and fused, or induced reciprocally between protodomains, it does not change the pool of accessible 7TMH structural domains with parallel membrane topologies.

While we have just considered the parallel combination of protodomains here, they can also assemble to form domains with inverted membrane topologies, and generate further structural diversity and complexity. For example, in **Figure 4.C** (upper right), the 3TMH conformation matches the TRIC protodomain. By inversion, it forms a domain through a totally different symmetric interface TM1-TM1’(=TM4) and TM2-TM2’(=TM5) (s11a/s22a) between two protodomains (see **Figure 1**), TM2 from each of the 3 domains engage in a further domain trimerization to form a D3 symmetric complex. The TRIC protein represents a great example of structural diversity through protodomain inversion. Further diversity can also be seen for example in MFS, with wider loop connections that enable also the intertwining of 3TMH protodomains (see **Figure 2.A** and the following discussion on MFS).

In following sections, we will envision the formation of asymmetric assemblies of 4TMH protodomains and a hypothetical duplication-fusion path leading to a pseudo-symmetric 7TMH fold. We will then see that, at the domain level, symmetric dimerization/assembly is a common pattern of GPCRs (**Figure 4.D**), similar to the rotating protodomain symmetric assembly process used in **Figure 4.C**. However, before we do, let us review an additional case that may shed some light on possible folding pathways of pseudo-symmetric folds.

#### Beyond inverted topology: Interdigitation of helices between protodomains

The folding of an interdigitated domain, such as MFS (**Figure 2.A**), must involve a very complex folding pathway. Paradoxically, it may be easier to imagine than in other 3TMH based symmetric domains, and may help us in the understanding of co-folding/co-assembly. MFS reaches a self-complementary conformation on both protodomains. How does it achieve this? We single out this domain as it may provide some clues that are more difficult to capture from packed domains. The self-complementarity of MFS is striking as it involves a helix by helix self recognition/self-assembly: we observe TM1-TM1 and TM3-TM3 interfaces (s11a/s33a) in protodomain interdigitation. The TM1-TM3 pairs form an antiparallel 4-helix bundle and leaves TM2 free for an “external” interaction. The parallel assembly of two domains does effectively happen through a double, symmetric, TM2-TM2 (s22a) interface. It seems therefore that each helix may provide a locally symmetric (antiparallel) self-complementary pair interface s11a, s33a between 3TMH protodomains within a domain and s22a between domains, all in antiparallel.

In MFS we can envision a reciprocal conformational selection model, implied by local self contacting helices TM1-TM1/TM3-TM3 and then finally TM2-TM2, all in antiparallel (s11a/s33a/s22a) among four protodomains in a D2 symmetry configuration (**Figure 2.A** and **Figure 1**). The folding itself may well involve a hierarchical pairing of likewise helices, suggesting that **these provide “local amino acid sequences which form stable interactions and serve as nucleation points in the folding process**”(Levinthal, 1969), and that these local amino acid sequences, organized through locally symmetric helical interfaces form self-complementary interfaces.

Other interdigitated folds with larger protodomains such as LeuT (PDB:2A65), composed of 5TMH repeats (albeit a more complex and less symmetric structure than MFS) also exhibits self-pairing of the first four helices TM1,2,3,and 4. This is not specific to just TMH proteins. In **Supplementary Figures S4 and S5**, we show an example of a globular protein exhibiting helix by helix self-assembly. One can observe symmetric pairing of individual secondary structure elements in globular proteins, whether beta strands (**Figure S4**) or alpha-helices (**Figure S5**). Symmetric pairing of protodomains is observed in membrane and globular proteins alike.

#### Can symmetry reveal conformational selection rules?

How do protodomains adopt or select the same conformation in the formation of a pseudo-symmetric domain? We discussed some possible folding/assembly scenarios, based on conformational flexibility. The only observable however is the folded state. Folded proteins in our examples belong to C2 cyclic or D2 dihedral symmetry point groups (Kosmann-Schwarzbach, 2009). Regardless of the folding or assembly scenario, ultimately protodomains adopt the same self-complementary conformation.

Group theory has been used successfully in chemistry and, where it applies, for example in IR spectroscopy and associated vibrational motions (Guichardet, 1986; Herzberg, 1945), selection rules have been established. The symmetries we are dealing with are not exact, and the analogy may seem remote, yet we can ask the question: Are there conformational selection rules? The case of MFS may point in that direction. If we consider domains with inverted topology that from packed protodomains, i.e. non interdigitated, they do not have the luxury of local symmetry complementarity of each transmembrane helix, yet there seems to always be at least one transmembrane self-complementary helix, or helix pairs, assembling in a locally symmetric manner. In doing so, they share the symmetry axis relating protodomains. FocA/Aquaporin is an example with TM1/TM4(∼=TM1), while TRIC shows two self-associating helices TM1/TM4 and TM2/TM5(∼=TM2), with the latter two forming in addition the central trimeric quaternary interactions (See **Figure 1**). Self-complementary association is not limited to membrane proteins of course, nor is it limited to alpha structures (see **Table 1**). Beta structures, in fact, show the same patterns. In the Immunoglobulin fold that we mentioned earlier, not only the protodomains are “inverted”, they also bind together through specific secondary structure elements: strands A-E(∼=A) and C-F(∼=C) (see **Figure S4**).

We have not investigated enough cases to possibly answer this question, yet it is tempting to relate the same or similar “local amino acid sequences which form stable interactions”(Levinthal, 1969) to pseudo-symmetric assembly/folding.

#### Did GPCRs emerge from 4TMH proteins through conformational evolution?

Pushing further the idea of protodomain conformational plasticity associated with gene/protein fusion events, we ask the question: can we envision a duplication-fusion-conformational evolution process from a 4TMH protein as an ancestor of GPCRs? If we take two 4TMH monomers and roll one against the other (as in **Figures 4.C and 4.D**), this time in an arbitrary way, keeping for example one fixed and rolling the second one against it, we can sample various dimer interfaces with symmetric or asymmetric arrangements. In **Figure S12**, we look at a TM1-TM2 vs. TM3-TM2 interface (with TM2-TM2 and TM1-TM3 contacts) between monomers (asymmetric contacts a23/12(=56) using the notation in **Figure 4**).

Assuming a duplication establishing a covalent linkage between two monomers TM4-TM5, one can envision a conformational rearrangement starting from an asymmetric interface to lead to a symmetric protodomains arrangement (**Figure S12**). This conformational change could happen through a concerted asymmetric swap of TM1 on one monomer and TM7-TM8 (TM3-TM4) on the second, involving the same 4TMH domain interface. This conceptual conformational change model is a variant to the 3TMH duplication-fusion process envisioned in the preceding paragraph for a symmetric monomer-monomer interface TM1-TM2 (s12) (see **Figure 4.C**). It gives however a rationale for the presence of the TM4 “linker”. It is interesting to note that in this case an asymmetric assembly could lead to a symmetric domain 8TMH fold, through conformational rearrangement, where further evolution can lead to the 7TMH pseudo-symmetric fold with attrition of the last helix or its transformation to the H8 helix as in in GPCRs.

The pentameric ligand-gated ion channels (pLGICs) are composed of 4TMH proteins. These 4TMH monomers present the interfaces of our hypothetical duplication example. Experimental structures of GLICs reveal a cavity accessible to phospholipids from the lipid bilayer between TM1 and TM4 (shown with a black wedge in **Figure S12**), that provides an allosteric binding site for a variety of general anesthetic ligands (Changeux, 2018; Nury et al., 2011). This is the intra-subunit interface involved in the proposed asymmetric swap in 4TMH. So, their M1-M2-M3(-M4) will map, upon duplication to TM1-TM2-TM3(-TM4) and TM5-TM6-TM7(-H8) protodomains of GPCRs. Specifically, the nicotinic acetylcholine receptor Achɑ7, a pLGIC with a 4TMH domain, has recently been shown (Kabbani and Nichols, 2018; Kabbani et al., 2013; King and Kabbani, 2016; King et al., 2015) to couple to G-proteins through a RxxR motif in its M3-M4 loop. This region maps to the RxxR motif of class C GPCRs and DRY motif of class A GPCRs at the end of TM3, which are in the G protein coupling regions of GPCRs. It has not escaped our notice that Achɑ7 (and other pLGICs) could be one of the potential protodomain sources for GPCRs. However, the duplication mechanism itself needs to be investigated further before an evolutionary link can be established between pLGICs and GPCRs.

#### Oligomerization in GPCRs

As observed in all 6/7/8-TMHs analyzed in this work, they all form higher order oligomers (see **Figure 1**). GPCRs are a special case, as they sample multiple homo-dimer interfaces. Effectively, the same principle of symmetric dimerization used in sampling symmetric interfaces between protodomains (**Figure 4.C**) can be applied at the domain level to explain observed dimers in GPCRs. In fact, *dynamic* rotating symmetric dimers have recently been observed in GPCRs (Dijkman et al., 2018; Xue et al., 2015). **Figure 4.D** shows two rotating dimers synchronized on rotation to maintain a symmetric organization sampling of homodimers, which has been observed in multiple GPCR crystal structures (Dijkman et al., 2018; Xue et al., 2015).

Symmetric dimers form the majority of experimentally determined protein quaternary structures. In the PDB, among ∼144000 structures of macromolecular complexes, ∼53000 exhibit quaternary symmetry, with ∼42000 (78%) presenting a cyclic symmetry (Korkmaz et al., 2018). The C2 symmetry represents the vast majority of symmetric structures with ∼32000 representatives, of which ∼31000 are homodimers. What makes GPCR dimers so special is their ability to form dynamic symmetric dimers of variable geometry, where dimeric states *or conformations* have been shown to be sampled during the lifetime of the dimer (Dijkman et al., 2018).

### C. Ligand recognition through sequence/structure differentiation

The TMH proteins cover a very wide range of functions due to their prime location at the cellular surface, which enables them to be utilized for jobs like transport of molecules in/out of the cell and sensing of extracellular signals to trigger intracellular responses. The cell membrane’s lipid bilayer environment is inherently asymmetric, where the outer lipid leaflet faces the extracellular (EC) side and the inner lipid leaflet faces the cytoplasm on the intracellular (IC) side. This asymmetry adds a natural directionality to their transporter and receptor functions. The TMH proteins embedded in this asymmetric environment can potentially feel different evolutionary constraints on their EC-facing and IC-facing TMH halves, which can leave a distinct function-based evolutionary signature in these protein halves.

To identify these potential evolutionary signatures in each of the TMH protein families being analyzed, a diverse set of proteins were identified in each studied TMH family along with one or more experimental structures available in each family oriented in the membrane by the OPM database (Lomize et al., 2012). The list of proteins and PDB ids used for each family is provided in the <u>Supplement File SF1</u>. The sequences of proteins in each family are aligned to each other using MAFFT (Katoh and Standley, 2014) and the EC/IC-facing halves of TM regions identified for each protein in the set by utilizing the membrane orientation of the reference structure in each family (see Methods section for details). The corresponding alignments for all TM regions in each protein family are shown in <u>Supplement File SF2</u> (*h* corresponds to the hydrophobic center for each TM in the OPM oriented configuration). The EC and IC loops as well as N- and C-termini of all proteins are ignored as they are usually of highly variable lengths are are difficult to align correctly. Skipping these loop regions from the analysis is reasonable as we have shown previously (Cvicek et al., 2016) through a TM-region-only alignment of all human GPCRs that the TM regions contain enough evolutionary information to enable an accurate phylogenetic representation of different GPCR families. The sequence similarity of the EC-facing and IC-facing TM halves were calculated for each TMH protein family to look for differences in evolutionary divergence of these EC and IC facing halves.

#### Evolution of ligand binding (EC) region vs G protein binding (IC) region in GPCRs

Structures of 35 diverse GPCRs were aligned to each other utilizing the TM regions as mentioned above, which provided a corresponding (potentially more accurate) sequence alignment. This sequence alignment was used to compare the IC-facing half (G protein coupling side) vs EC-facing half (ligand binding side). **Figure 5.A** shows the sequence similarity across a diverse set of 35 GPCRs to assess the extent of divergence in each TM, in each of the two protodomains, and in each half (EC-facing and IC-facing) of the TM domains.

**Figure 5.**
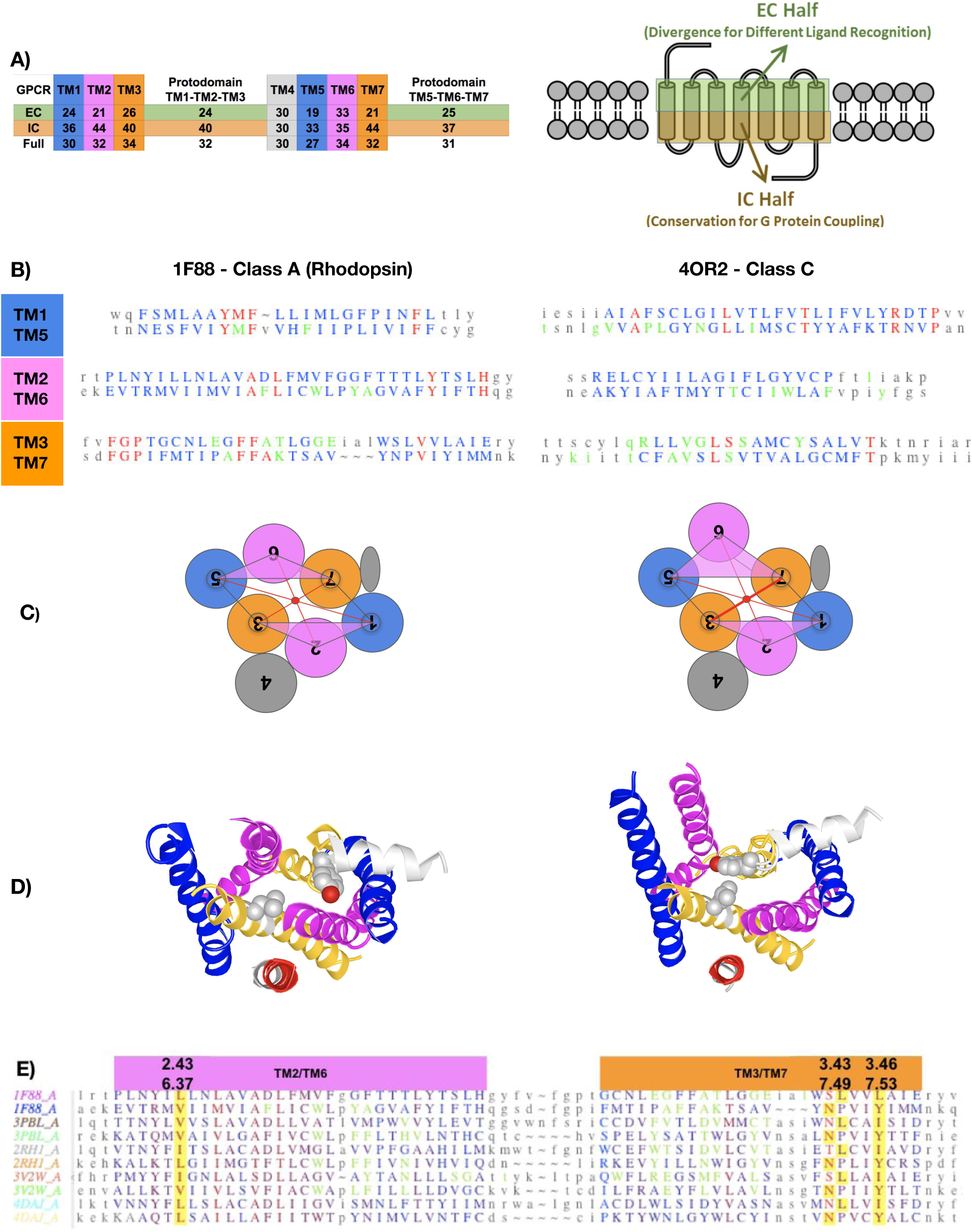
Deconstruction of GPCR Domains. **A) Right panel:** Definition of **TMH halves facing the Extracellular (EC) and Intracellular (IC) sides**. **Left Panel: Sequence similarity score** (see Methods section for details) of the aligned EC half, IC half, and Full TM sequences for each of the 7 TMs for a diverse set of GPCRs spanning all classes and subclasses. Protodomain 1 and 2 scores are also given (averages over 3TMH). Colors correspond to TMHs vertically and EC and IC halves horizontally. EC halves show a lower score for each TMH compared to IC halves in GPCRs (see text). **B) Pairwise Protodomain alignment** (RMSD: 1F88 = 3.24 Å; 4OR2 = 3.31 Å). The symmetrically conserved pattern, especially in TM3/TM7 surrounding the ligand, in each domain is idiosyncratically conserved (Red = conserved, Green indicates ligand binding/proximity residues (in less than 4 Å distance). In these examples, specifically FFA(T/K) for bovine Rhodopsin and V(G/S)LS in human metabotropic glutamate receptor. For Rhodopsin, TM1 shows a YMF pattern in TM1 and a A(D/F)L pattern in TM2 (see discussion on the D2.50/F6.44 alignment). While YMF and FFA patterns of protodomain 1 are in direct contact and move in a concerted way in an inactive (1F88) to active (2X72) transition, one cannot point to contacts between these two motifs on protodomain 2. In the metabotropic glutamate receptor, the structural homology of TM1/TM5 extends beyond into the loop regions with the RxxP pattern. In Rhodopsin, this is the case with an FGP motif before TM3/TM7. Rhodopsin is unique in that its ligand is covalently bound to the TM7 Lysine, hence this is the only case where we inserted a gap vs. TM3 to optimize the alignment. **C-D) Rhodopsin inactive vs. active conformational change** seen from the IC side (binding G-protein not shown for clarity). **Left inactive** (PDB: 1GZM)**, Right active** (PDB:6CMO). The optimum alignment as shown is obtained from VAST+(Madej et al., 2014) [see **Figure S13** for details). The **<u>iCn3D visualization</u>** (using the “alternate” command) gives a good grasp of the conformational change. The two protodomains, despite the conformational change, affecting the second protodomain mostly on TM6 still align within a 3 Å RMSD range whether active or inactive (see Supplementary Figure S3.A for detailed pairwise protodomain alignments). The conformational rearrangement is distributed on many residues/degrees of freedom. TM3-TM7 helix pair plays a central role in coupling and maintaining symmetry, and bringing the regions 7.49-7.53 with 3.43-3.46 in contact, with a noticeable change in Y 7.53 conformation and orientation. **E) Multiple alignment of protodomains of a number of Class A GPCRs** showing symmetry related residue pairs (highlighted in yellow) also involved in key contacts (in green) [See text for details]

The sequence similarities showed that EC-facing half of TM regions in GPCRs has evolved more than the IC-facing half of TM regions for all seven TMs, consistent with the fact that GPCRs sense a huge chemically diverse set of ligands using their EC facing half, but they couple to only a small family of G proteins using the IC facing half. The sequence similarities also show that TMs 5, 6, and 7 (protodomain 2) have evolved to the same extent as the TMs 1, 2, and 3 (protodomain 1) (31% vs 32% respectively, as seen in **Figure 5.A**).

These results show that functionally GPCRs live in a highly asymmetric environment due to G protein coupling on one side and ligand binding on the other side, which is captured in higher sequence similarity in the IC region for both protodomains (40% and 37% respectively vs 24% and 25%).

##### Comparing EC vs IC regions for other TMH proteins

The **Supplementary Figure S14** shows the sequence similarity of the EC facing and IC facing TM halves of selected other TMH proteins: Aquaporins, Foca, PnuC, TRIC, and MFS. Some of the features that emerge are as follows. The two protodomains in each of these families have diverged to the same extent, similar to the GPCRs. Aquaporins have the NPA motif in TMs 2B and 5B that impart high conservation to those segments. PnuC has a conserved WxxW in the IC half of TM6 that binds to its substrate, hence it is more conserved than other TMs. TRIC has a conserved GG motif in TMs 2/5, and since it is an ion conduction channel it contains conserved residues along the whole TM length, so no EC vs IC patterns emerge like in GPCRs.

#### Symmetrically related TM3/TM7 ligand binding in GPCRs

The architecture of GPCR domains clearly separates the ligand binding on the EC side from G-protein (or Arrestin) coupling on the IC side. There is a low sequence conservation (high sequence variability) on the ligand binding region and a high sequence conservation (low sequence variability) on the G-protein coupling side, as was shown above.

Considering only the transmembrane helices, ligand binding residues can be distributed on all TMHs, however most ligands bind effectively to the 3 helices (TM5, TM6, TM7) in protodomain 2 and TM3 in protodomain 1. This can be seen in multiple examples of class A GPCRs (green residues in **Figure 5.B** and **Supplementary Figure S3**). A few cases, however, may involve residues in TM1 and TM2. Hence in terms of pseudo-symmetry and ligand binding, it involves essentially the TM3/TM7 pair, with anchor residues for ligand binding positioned symmetrically. In TM3, ligand binding residues are mostly in positions 3.28, 3.32-3.33, 3.36-3.37 vs. 7.35, 7.39, 7.42-7.43 in TM7 (using the Ballesteros-Weinstein numbering of TM residues for GPCRs (Visiers et al., 2002)). TM3-TM7 is the only obligatory transmembrane helix pair for ligand binding, and their two binding regions are symmetrically related.

In the second protodomain, TM6 residue W6.48 is a highly noticeable ligand binding residue, along with residues 6.51-6.52 around the conserved residue P6.50. The residue F6.44 (in prodomain 2) can also sometimes be involved in ligand binding. Its symmetry related, highly conserved residue D 2.50 on TM2 (in protodomain 1) does not bind to the ligand. In fact, D2.50 binds to Na^+^ that has been shown to correlate with the functional state of the receptors(White et al., 2018). It can be symmetrically paired with either F5.44 or W5.48, both highly conserved. This points to a co-evolution of D2.50 vs. F6.44 and W6.48. In fact, in pairwise protodomain alignments (see **Supplementary Figure S3.A**), we have alternative protodomains alignements where D(2.50) can be equivalenced to W(6.48) instead of F(6.44). It is a common feature in helical proteins alignements to see translations of helices along their helical axis, shifting residues in positions +/-4 (see note in section Methods - Protodomains delineation).

#### G-protein binding and the TM3/TM7 paired interactions

Additional functional significance emerges around the TM3/TM7 paired interactions. Just below the ligand binding area, the highly conserved class A GPCR residues S3.39 and N/S7.45 (or S7.46) match symmetrically across the protodomains. They are Na^+^ binding residues (White et al., 2018). Below the Na^+^ binding area, when one looks at helix-helix contacts that change upon activation (Cvicek et al., 2016; Venkatakrishnan et al., 2016), TM3 and TM7 form contacts between residues at positions 3.43-7.49, 3.43-7.53, 3.46-7.53 in the active conformations **but not in the inactive conformation,** as observed previously (Cvicek et al., 2016; Venkatakrishnan et al., 2016).

The two regions in TM3 and TM7 match symmetrically whether in the active or inactive state, yet they form direct contacts in the active state. In **Figures 5.C and 5.D**, we show the case of Rhodopsin, seen from the intracellular side where two Leucine residues in position 3.43 and 3.46 (and similar I/L/M residues in other class A GPCRs) form contacts with the highly conserved Tyrosine 7.53 (see protodomain alignments and highlights in **Figure 5.E**). Interestingly both protodomains in active and inactive states align within a 3 Å RMSD (see **Figures 5.C, 5.D, 5.E**, and also **Supplementary Figure S3** for examples).

These observations show that the EC-facing halves of TM3/TM7 provide endogenous ligand contacts along with Na^+^ mediated contacts, and their IC-facing halves provide direct contact with each other upon activation. **This places the TM3/TM7 interface at the heart of GPCR functional action**. Previous studies (Venkatakrishnan et al., 2013) have made the case for the central role of TM3 in GPCR function, however, our protodomain hypothesis suggests that TM3 and its pseudo-symmetric partner TM7, together, play a “pivotal” role in that function.

##### Each pseudo-symmetric protein shows a unique protodomain co-evolution pattern

Each GPCR domain has its own evolution history, however, each maintains an internal homology and an identity pattern for some symmetrically equivalent residues. Such identity patterns are idiosyncratic, they are different between various GPCRs. We observe such symmetric sequence pattern “coincidences”, or identities, in a number of GPCR protodomain pairwise alignments.

In **Figure 5.E** and in **Supplementary Figure S3**, we present pairwise alignments for a variety of GPCRs showing pseudo-symmetric sequence/structure similarity patterns. No sequence pattern is shared by any two GPCRs, unless they are highly related. This is a puzzling observation: in a number of GPCRs, a *seemingly conserved* sequence pattern emerges in the ligand binding region of TM3 vs. TM7 when aligning protodomains. It is the case in Rhodopsin or Metabotropic glutamate receptors 1 and 5. In the case of Rhodopsin (1F88), for example, the FFA(T) pattern in TM3 matches FFA(K) preceding the Lysine residue in TM7 (that binds to retinal); in the case of Metabotropic glutamate receptor 1 (4OR2) the pattern is VxLS, as shown in **Figure 5.B** (see also in **Supplementary Figure S3**). CB1, another class A GPCR (PDB:5U09) has a VFxF pattern. Given the fact that sequence identity is usually low when comparing protodomains (see Methods - Protodomain delineation) this has attracted our attention. Whether this is of evolutionary significance in the classical sense is unclear.

Such patterns are specific to a given protein and close relatives, in one word they are idiosyncratic, and do not extend to distant homologs. For example, Squid Rhodopsin does not present the pattern on both protodomains (PDB:2ZIY), but the human (PDB:6CMO) and bovine (PDB:1F88) Rhodopsin do. Such patterns can also be common in the highly hydrophobic region of TM1 vs. TM5 (see for example chemokine CXCR4 PDB:3OE9 or histamine H1 PDB:3RZE) in **Supplementary Figure S3**). These “internal conservation” patterns cannot be looked at in the same way as domain level patterns across families as we are used to, since families have conserved residues for folding or functional reasons in separate, distinct, parallel, independent evolutionary histories. Internal conservation is of a completely different type in most cases, and especially in GPCRs. Protodomains have a common co-evolution history, where it is quite unlikely that symmetrically equivalent pairs are conserved, unless there is a reason, be it folding or functional, to maintain identity and this should show beyond a single or a set of highly related proteins. The pattern, when it occurs, seems more like a convergence towards identity or coevolution of identity rather than a conservation. Another element we can notice is that in such patterns, such as FFA motif in Rhodopsin, the residues involved, some facing the lipid, are framing the ligand binding pocket.

##### Co-evolutionary conserved residues and Function

The diversity and complexity of GPCRs is such that it is extremely difficult to infer co-evolution patterns between protodomains through a simple observation of a sequence alignment. While the D(2.50)/F(6.44)-W(6.48) pattern is relatively easy to pick (see earlier), there are certainly other co-evolved pairs (or larger sets of residues) in GPCRs to detect, as we usually find in pseudo-symmetric domains (Youkharibache, 2018 (in press)). Another set of residues that are related in function are naturally the sodium binding residues D(2.50), and the symmetrically related pairs in TM3 and TM7: S(3.39) and N/S(7.45) [or S(7.46)]. Here the co-evolution involves the two protodomains and the Na^+^ ion. Systematic studies would be needed, and can now be envisioned to identify co-evolution patterns, between protodomains, especially in the case of GPCRs with ligands. The relations are certainly very complex, and we are not used to or trained to capture such patterns. A symmetric decomposition of domains into protodomains may help.

Let us note that sets of conserved co-evolved residues, or in general, sets of residue positions interrelated in a pattern, will help in the development of a method to align protodomains from sequence down the road. For example we observe that if one aligns, structurally, the regions in TM2 and TM6 surrounding the D(2.50) - F(6.44)/W(6.48) anchor, the other helices are automatically positioned for the whole protodomain structural alignment. With a few more patterns interrelating residue positions in GPCRs, a constrained sequence alignment may well give results comparable to structural alignments, as performed in this study. We have observed this in small beta barrels (Youkharibache et al., 2018) that, like the pseudo-symmetric membrane proteins, exhibit a very low sequence conservation at the domain and at the protodomain level, yet both similarity and complementarity patterns emerge. Sequence alignment has been based on similarity until now. It needs to be extended to include complementarity. Self-complementarity of protodomains and their interrelated patterns should provide constraints for accurate sequence alignment of protodomains in pseudo-symmetric domains. Coupling this information with co-evolutionary contacts (Adhikari and Cheng, 2018; Adhikari et al., 2018) aimed at predicting 3D structure from sequence should improve the structure prediction accuracies for such domains.

## Conclusions

In this study, we have established a parallel between diverse 6/7/8-TMH proteins that shows a parallel evolutionary path of duplication-fusion and symmetric assembly of 3/4-TMH protodomains. This parallel path does not necessarily imply that these proteins have a common origin in sequence space. The formation of pseudo-symmetric membrane proteins is similar to what is observed in globular proteins. What stands out however among 6/7/8-TMH proteins, is the formation of a diverse set of folds from conformationally variable 3/4-TMH protofolds. This should also be put in perspective with a significant overrepresentation of 7TMH proteins in the surfaceome (Bausch-Fluck et al., 2018).

A reason for the evolutionary success of 7-TMH proteins may well be a structural one. The creation of an almost cylindrical unit provides a molecular device with a natural directionality to channel/transport molecules or ions across a membrane, or for transmembrane signaling. Duplication and symmetric assembly of a 3/4-TMH around an axis normal to the cell membrane looks like a simple mechanism to get to the minimum size cylinder with a directional function. In the examples selected, some folds may share a function, such as transporting sugar, yet they may or may not have evolved from the same ancestor (SWEET vs PnuC). Conversely, functional diversification has been obtained from common ancestors (FocA vs. Aquaporin). In addition, 3/4-TMH protodomains provide cohesive energetically stable supersecondary structural units that can self assemble. The biophysical evidence is only now beginning to emerge (Min et al., 2018).

We have a general molecular self assembly principle at work, in membrane and globular proteins alike, forming symmetric oligomers as well as pseudo-symmetric tertiary structures (domains) and even pseudo-quaternary domain assemblies (as in the case of MFS). The coincidence between symmetry axes in helical membrane proteins and a lipid membrane axis system, and for many of their functions, tends to imply that a large number of membrane proteins should be symmetric. GPCRs, whose function does not seem to require symmetry, nevertheless exhibit pseudo-symmetry, where the second protodomain (TM5-TM6-TM7) undergoes conformational change upon receptor activation to accommodate the G protein.

We have reviewed possible scenarios for the parallel evolution of a variety of 3/4-TMH protodomains, that we termed *conformational evolution*. Naturally, this work calls for further research to test ideas through experiments where related sequences may lead to diverse folds, and/or folding experiments to understand the mechanism(s) of protodomain assembly and membrane protein folding. In particular, the local pairing of transmembrane helices in inverted, interdigitated protodomains that, as far as we know, have not been found in oligomers, deserves further attention.

We may need to revisit the methodology developed earlier for a more systematic and accurate symmetry determination of membrane proteins. This study also highlights a need for a more systematic study of co-evolution of protodomains, especially of GPCRs. This should be possible as the number of membrane proteins structures are now growing at the same exponential pace as globular proteins (https://www.rcsb.org/stats/growth/overall), with the GPCRs leading the charge.

## Methods

### Pseudo-symmetry Protodomain Analysis (PSPA) Method

We have reviewed the method in detail elsewhere (Youkharibache, 2018 (in press)). The method is simple and involves 2 initial steps: **Symmetry detection** and **Protodomain delineation**. Symmetry detection gives the symmetry point group and a first delineation of protodomains. Both tertiary and quaternary structural symmetries can be determined at the same time. The second step is usually required to optimize protodomain boundaries and structural alignment for accurate protodomains delineation. This then opens the door to any desired analysis that a structural alignment of tertiary but also quaternary structures may enable.

Matching sequence patterns between protodomains resulting from their structural alignment is similar, at first sight, with any domain level analysis. However, one should note that an alignment of protodomains is different from a classical domain alignment leading to families and superfamilies. One should not expect “internal” sequence conservation in the same sense. Protodomains have not evolved separately and conserved residues for functional or folding reasons in the same way. Protodomains in a domain have co-evolved jointly to reach a differentiated idiosyncratic function while maintaining a pseudo-symmetric fold.

#### Symmetry detection

In many cases, recently developed computer programs allow the detection of internal pseudo-symmetry in tertiary structure (Kim et al., 2010; Myers-Turnbull et al., 2014) (See **Table 1**). We use the program CE-symm to that effect. A newer version of the software allows quaternary symmetry analysis of multi domain complexes at the same time (https://github.com/rcsb/symmetry). There are many cases where we have to adapt program parameters to detect symmetry and obtain structurally aligned protodomains, depending on departure from perfect symmetry and structure quality. In some cases, one has to use interactive alignment software to align a domain onto itself, that requires a necessary visual inspection at each and every step. This is particularly true for GPCRs that present a wide range of structures, resolution, co-crystallization domains, and conformational states. In all cases we optimize protodomains’ delineation through interactive structural alignment for accuracy.

#### Protodomains’ delineation: Optimization through structural alignment

Protodomains’ alignment may highlight key residues that may be internally conserved for a structural reason (folding/assembly) or for a functional reason, such as ligand binding. In most cases, the degree of overall internal conservation is very low. This is a hallmark of most pseudo-symmetric domains, unlike most domain/family level sequence-structure conservation, except for some clear duplication cases where homology between protodomains may be as high as 40%. In the examples considered in this paper, TRICs show such a clear duplication pattern of protodomains with 29% identity and no insertion/deletion between the duplicated protodomains (Kasuya et al., 2016; Su et al., 2017) (see alignments in **Figure S1**). The low level of “internal conservation” observed in most cases however is likely due to a long protodomain-protodomain coevolution within each and every protein domain but also at quaternary interfaces.

In order to call internally conserved residues, an accurate structural alignment of protodomains constituting a domain is required, and each and every protodomain pair within a family or superfamily will be different. The few invariant (“internally conserved”) residues at equivalent positions, may not be invariant across domains in a family or superfamily, but if they are, they will no doubt bear a particular significance. These cases are rare, and it may be best to talk about coincidence than conservation until further evidence is gathered. More often, residues (that may or may not be in contact) at symmetrically equivalent positions, may be conserved as pairs across families, they would show as individually conserved in domain level alignments (see section “sequence patterns in pseudo-symmetric domains”).

Using pseudo-symmetry provides a framework for a hierarchical structural analysis. It allows a deconstruction of protein domains in well-defined parts that may also lead into evolutionary and/or functional analysis. The reconstruction of a domain from its parts (protodomains) leads into a co-evolutionary analysis of the parts and their interfaces, and in a better understanding of molecular self-assembly. This opens perspectives in developing the analysis method further in that direction. We focus essentially on a high level structural analysis in this paper. The current method should be seen as providing a rigorous data organization framework to identify co-evolution patterns that may help in the understanding of various functional levels across GPCRs. The set of known GPCRs should provide a rich dataset to envision the use of machine learning to identify co-evolution patterns within that pseudo-symmetric framework.

The Root Mean Square Deviation (RMSD) is the quantitative measure of structural similarity we are using all along. A small RMSD denotes a strong structural similarity. This criterion is being used systematically across all structural comparisons in this paper as is done across the literature. In comparing helical structures, we consider a RMSD lower than 3.5 Å to describe a good structural match. However this is a unique number with many contributions between two sets of residues, two substructures. Hence, it does not capture the details at the residue level, but is a good overall match criterion.

The majority of our domain or protodomain alignments on GPCRs lie between 2.5-3.5 Å RMSD. A note of importance is that within this range, we can have alternative sequence alignments. It is common in helical alignments to see translations of helices along their helix axis, shifting residues in positions +/-4. Most helical protein structural alignments match structures within ∼3-4 Angstroms in backbone RMSD, even when sequence matching is indicative of high homology, and a helical turn translation of a helix may not drastically change such an overall RMSD.

#### Software programs

Two programs allow the automatic detection of pseudo-symmetry in protein domains, SYMD(Kim et al., 2010) and CE-symm(Myers-Turnbull et al., 2014). We use the latter for automatic detection. The program does a good job at capturing both quaternary and tertiary levels of symmetry. Some results are summarized in the **Figure 1** for all the proteins studied in this paper. They all show two levels of symmetry, one tertiary and one quaternary that in the case of inverted membrane topologies do combine to give dihedral symmetries. GPCR pseudo-symmetry is not detected, however, by the software, except in the case of <u>4OR2</u> (see above). Interactive structure alignments (optimization) for all structures in this paper are performed with the Cn3D software (Madej et al., 2012; Wang et al., 2000). All GPCR protodomain pairwise and multiple sequence/structure alignments are optimized with Cn3D. Domain level structural alignments can be readily available from NCBI VAST structural database (Madej et al., 2012), and from VAST+(Madej et al., 2014) for quaternary assemblies alignments. VAST+ alignments capture conformational changes in tertiary domains and quaternary assemblies, in particular for GPCR conformational changes. All these structures can now be visualized with web based visualization and analysis software iCn3D(Jiyao Wang, Philippe Youkharibache, Dachuan Zhang, Chris Lanczycki, Lewis Geer, Renata Geer, Aron Marchler-Bauer, Tom Madej, Lon Phan, Minghong Ward, Shennan Lu, Gabriele Marchler, Yanli Wang, Steve Bryant, 2018 (Submitted for publication)) developed at NCBI and available as open source (https://github.com/ncbi/icn3d). iCn3D allows the creation and exchange of annotated 3D structure visualizations in parallel with sequence (1D) in particular. 3D visualization links are given in **Figure 1**.

#### Notation, Coloring and Visualization

Naming of secondary structure elements with repeats can be confusing, as elements are numbered in sequence, within a protodomain, within a domain, and within a multidomain protein (such as MFS). Hence that notation has to vary depending on context. We cannot avoid a certain imprecision due to the multiple numbering of a given element. Also numbering in sequence may not match what is used in naming different topologies adopted by 3TMH, such as 123, 231, 312.

Color is the most important element of distinction and recognition used in Figures. In a 3TMH protodomain we use sequential colors BLUE, MAGENTA, and ORANGE for transmembrane helices that we name TM1, TM2, TM3 respectively. Whatever number a protodomain’s TMH ends up having, such as TM5, TM6 and TM7 in 7TMH domains, they will be colored blue, magenta and orange.

We use schematic 2D projections from the extracellular (EC) side for visualization of protein transmembrane domains, except where specified. These are idealized as if TMHs would be exactly perpendicular to the membrane, while in reality they could vary by as much as 45 degrees each, hence in some cases we could have two neighboring helices that are orthogonal to each other in the membrane. This could result in principle in different views from the EC and IC, yet it is not the case in the proteins examined in this study. Let us note however that a clockwise arrangement of helices seen from the EC side would appear as counterclockwise from the IC side, its mirror image. 3D visualization is available through iCn3D(Jiyao Wang, Philippe Youkharibache, Dachuan Zhang, Chris Lanczycki, Lewis Geer, Renata Geer, Aron Marchler-Bauer, Tom Madej, Lon Phan, Minghong Ward, Shennan Lu, Gabriele Marchler, Yanli Wang, Steve Bryant, 2018 (Submitted for publication)) web links in the **Figure 1**.

#### Protein structure classification

Two major classifications SCOP(Chandonia et al., 2017; Lo Conte et al., 2000; Murzin et al., 1995), and CATH(Dawson et al., 2017; Orengo et al., 1997), have been used for a long time, and more recently ECOD(Cheng et al., 2014; Schaeffer et al., 2017). We chose SCOP for its fold classification, based on geometrical criteria and manual curation, as “The method used to construct the protein classification in scop is essentially the visual inspection and comparison of structures, though various automatic tools are used to make the task manageable”(Murzin et al., 1995). This, in essence, has been our simple but accurate approach to pseudo-symmetry analysis and protodomain delineation through self protein alignment (see earlier), especially for GPCRs, for two reasons. First, at this stage, and even more when we started, no automatic tool will identify their symmetry and protodomains alignments accurately and systematically (see earlier). Second, evolution of membrane proteins may have, precisely, a particular geometrical drive that transcends current views of sequence based evolutionary paths. Hence the notion of fold and protofold in our context, constitutes a central geometrical Element(Euclid, 300 BC) of major importance.

### Evolutionary Structural Analysis of Different Transmembrane Protein Families

Structures related to the specific protein family being analyzed are pulled from NCBI’s Conserved Domains Database (CDD)(Marchler-Bauer et al., 2003, 2015, 2017). The corresponding fasta sequences are pulled from Uniprot (UniProt Consortium, 2018) input into our PredicTM program (Goddard et al., 2010) that implements the hydropathy analysis (von Heijne, 1992) to generate modified-fasta (mfta) files containing hydrophobic TM domains. Each protein’s TM regions are extended based on the protein data bank (PDB)(Burley et al., 2017) structures. The hydrophobic centers are determined by looking at the PDB structures aligned by the OPM database (Lomize et al., 2012) to an implicit membrane (membrane middle defined by z=0 plane) and selecting one residue as a hydrophobic center in each TM domain with the C *α z*-coordinate closest to zero (designated *h* in <u>Supplementary File SF2</u>). The TM regions and hydrophobic centers are recorded in the protein’s mfta file. Then, the CDD protein family’s sequence alignments are downloaded from NCBI’s CDD website. This sequence alignment consists of the ten most diverse proteins in the family. Using HMMER, the proteins with PDB structures are aligned to the existing CDD multi-sequence alignment. Based on the alignment, a consensus for the TM regions and hydrophobic centers is reached using the proteins with PDB structures. This consensus is used to assign the TM regions and hydrophobic centers in the respective mfta files for all proteins in the new CDD multi-sequence alignment. All proteins include both with and without structures. After using a custom script to cut up the sequences using the TM regions and hydrophobic centers, the resulting fasta files are consolidated into combined fasta files for each TM as well as each TM’s intracellular facing half and extracellular facing half determined from the hydrophobic centers determined earlier (**Figure 5.A**). Finally, the consolidated multi-sequence fasta files are run through a custom scoring function defined below based on the blosum62 matrix and the scores are recorded for each TM and for each EC and IC half. The TM lengths of the corresponding TMs across the two protodomains were not matched to each other to capture the true divergence of the TM regions at this stage. A handful of TM regions were not present in the CDD alignments, in which case those were aligned manually.

#### Scoring of Structure-Based Intracellular and Extracellular facing Halves’ Sequence Alignments

The scoring program divides the sequences in an alignment into vertical columns. Each amino acid in a column is compared to others in the same column and given a score. The scores are derived from custom scores based on the blosum62 matrix. A blosum62 matrix score of 4 or greater results in a score of 1. A blosum62 matrix score of 3 results in a score of 0.75. A blosum62 matrix score of 2 results in a score of 0.50. A blosum62 matrix score of 1 results in a score of 0.25. A blosum62 matrix score of 0 results in a score of 0.125. All negative scores result in a score of 0. This scoring is repeated for every column in the alignment. The scores are then added together and divided by the number of comparisons. After multiplying the result by 100, we get the similarity percentage between all sequences in the alignment, which captures how much each respective domain has diverged, smaller the number higher the divergence.

## Supporting information

## Acknowledgements

We would like to thanks NCBI collaborators: Tom Madej and Jiyao Wang for their assistance with the use of VAST+ and the targeted development of iCn3D respectively, Aron Marchler-Bauer and James Song for their support in the use of NCBI’s Conserved Domains Database (CDD) used for evolutionary analyses and the newly updated GPCR Conserved Domains; and Antoniya Aleksandrova for her help in getting membrane datasets. This work was supported in part by the NIH intramural program (PY) and by the startup funds at CSUN (RA). AT would like to thank the Abrol lab members for helpful discussions.

## Author Contributions

PY and RA wrote the main manuscript text. PY, AT, and RA generated the data. All authors analyzed the data and reviewed the manuscript.

## Competing interests

The authors declare no competing interests.

## Notes

#### Summary of Updates

Title shortened Abstract/Conclusion updated Figures/Tables renumbered

