## Supplementary material for "7-Transmembrane Helical (7TMH) Proteins: Pseudo-Symmetry and Conformational Plasticity"

### SUPPLEMENTARY INFORMATION

| | | |
|---|---|---|
| MFS | 5EQI | https://d55qc.app.goo.gl/wcE7nXES2yhE4CQr6 |
| TRIC | 5WUF | https://d55qc.app.goo.gl/XUkPY7iPmR8LVJ8c9 |
| AQP1 | 3NE2 | https://d55qc.app.goo.gl/ZpabKYatPRrpnWF39<br>https://d55qc.app.goo.gl/pN9hZKMbiS4VML5H6 << new |
| SemiSWEET Apo vs. Ligand | 4QNC 4QND | https://d55qc.app.goo.gl/i9vWJfzXEcL8q2A69 |
| SWEET vs. SemiSWEET | 5CTH 4QND | https://d55qc.app.goo.gl/9NfEK6KmideLHYt1A<br>Use keyboard “a” letter or “Alternate” command to visualize alternatively the 2 aligned structures of SWEET (PDB:5CTH) vs. 3TMH monomer A of SemiSWEET (PDB:4QND) (aligned only on 3TMH protodomain 1 )<br>Use “a” alternate command to alternate between structures |
| PnuC | 4QTN | https://d55qc.app.goo.gl/S1atQgdGptdt7bCs6 |
| GPCR C | 4OR2 | https://d55qc.app.goo.gl/ZFChMCsBRER3uSbD7 |
| GPCR Aα | 5G53 | https://d55qc.app.goo.gl/igoFR7sh9hZdjK1L9 |
| Rhodopsin | 1GZM | https://d55qc.app.goo.gl/KtAe6nkkSJ7fm3aHA |
| Rhodopsin active-inactive | 6CMO 1GZM | https://d55qc.app.goo.gl/KLJg7G6Zx3g9hju4A |
| GLIC | 5VDI | https://d55qc.app.goo.gl/keoUw5UrpHs9wKY77 |

**Table S1 - 3D visualization links** - using iCn3D

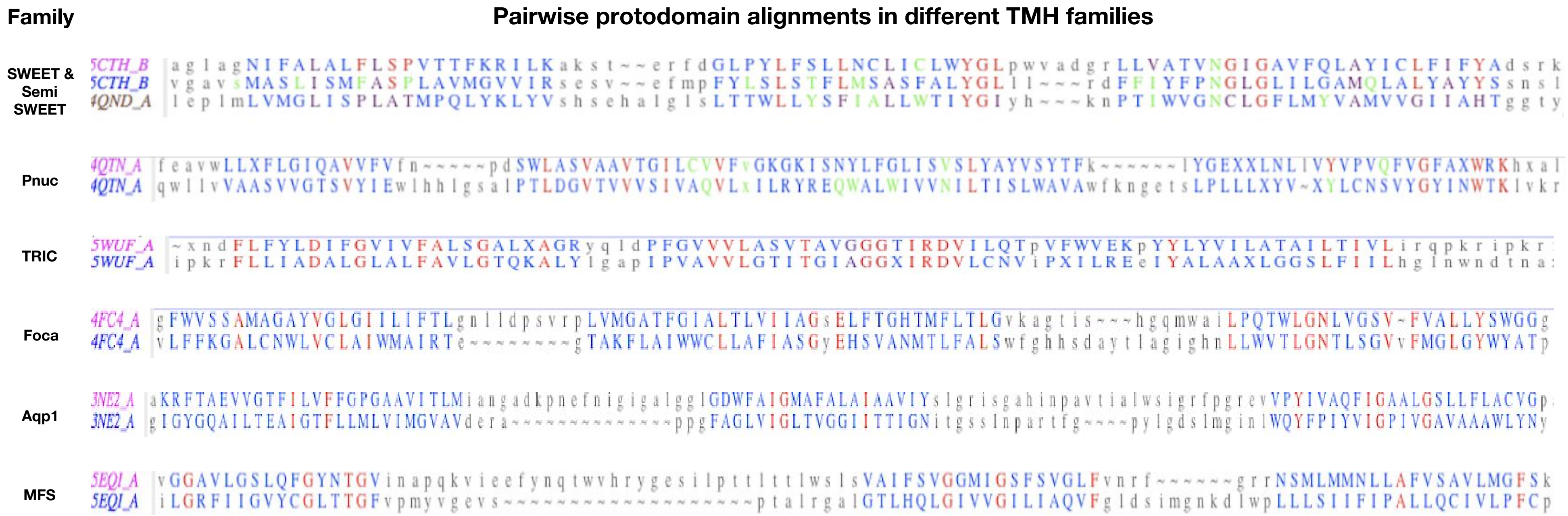

**Figure S1 - Protodomains pairwise alignment of:** Sweet protodomains 1.36 Å 5CTH vs SemiSweet 1.98 Å 4QND, Pnuc: 3.09 Å/1.26 Å 4QTN, Tric: 1.88 Å/1.53 Å, FocA: 4FC4 1.83 Å, Aqp1 3NE2 1.94 Å, MFS 1.70 Å - after optimisation of protodomain boundaries for best structural match. Notice the lower RMSD than in Table 2 when considering solely a structure match with optimized structural optimization (see Methods) - for example for Sweet 1.36 Å vs 2.18 Å when factoring in the axis of symmetry offset used in CE-symm [**Figure 1**] .

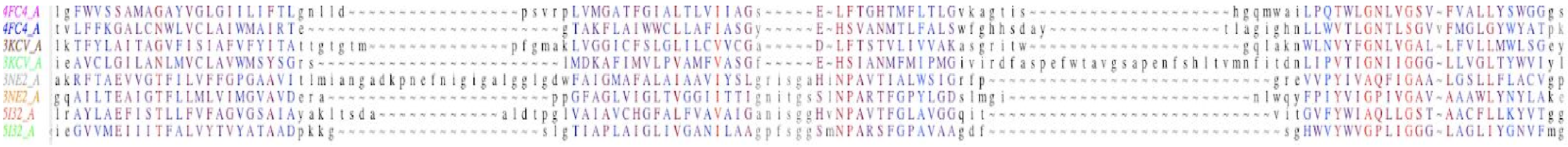

**Figure S2 - Protodomains multiple alignment of FocA vs AQP1.** FocA **(**PDB: 4FC4, 3KCV), AQP1 (PDB: 3NE2, 5I32) Protodomains RMSD vs. first protodomain: **FocA** 1.91A, 1.55A/2.05A; **AQP1** 2.33A/2.57A, 2.26A/2.98A. The sequence match is poor between FocA and AQP1, apart from in TM3 with motif involving a [G/A]xxx[G/S][G/A/S] motif, especially in the protodomain2 of 3KCV and 5I32 with the close sequence match **GxxxIGGGL** of FoCA and AQP1**.**

Class

#### Pairwise GPCR protodomain alignments

```
A   1F88_A  qFSMLAAYMF~LLIMLGFPINFLtlyvtvq~~~~~~~~~~~~~~hkklrtPLNYILLNLAVADLFMVFGGFTTTLYTSLHgyfvFGPTGCNLEGFFATLGGEialWSLVVLAIEry
    1F88_A  nNESFVIYMFvVHFIIPLIVIFFcygqlvftvkeaaaqqqesattqkaekEVTRMVIIMVIAFLICWLPYAGVAFYIFTHqgsdFGPIFMTIPAFFAKTSAV~~~YNPVIYIMMnk

Aa  2X72_A  tnNESFVIYMFvVHFIIPLIVIFFCYgqlvftvkeaaaqqqesattqkaekevtrmVIIMVIAFLICWLPYAGVafyifthqgscFGPIFMTIPAFFAktsavynpviyimmnkqfr
    2X72_A  wqFSMLAAYMF~LLIMLGFPINFLTLyvtvqhkk~~~~~~~~~~~~~~lrtplnyiLLNLAVADLFMVFGGFTTtlytslhgyfvFGPTGCNLQGFFAtlggeialwslvvlaiery

    6CMO_R  pwqFSMLAAYMFLLIVLGf~PINFLTLYVTVQhkk~~~~~~~~~~~~~~lrTPLNYILLNLAVADLFMVLGGFTSTLYTSLHgyfvFGPTGCNLQGFFATLGGEIALWSLVVLaiery
    6CMO_R  evnNESFVIYMFVVHFTIpmIIIFFCYGQLVFtvkeaaaqqqesattqkaeKEVTRMVIIYVIAFLICWVPYASVAFYIFTHqgscFGPIFMTIPAFFAKSAAIYNPVIYIMMnkqfr

B   5EE7_A  nmgfwWILRFPVFLAILINFFIFVRIVQLLvaklrarqmhhtdyafRLAKSTLTLIPLLGVHFVVFAFVtdeha~~~~~~~~~~~~~~~~~~~~qgtIRSAKLFFDLALSSFQ~GLLVAVLYCFInk
    5EE7_A  yssfqVMYTVGYSLSLAALLLALAILGGLSkl~~~~~~~~~~~~hcTANAIHANLFLSFVLKASAVLFIdgllrtrysqkieddlsvstwlsdgavaACRVAAVFMQYGIVAnYCWLLVEGLYlhn

C   4OR2_A  sniesIIAIAFSCLGILVTLFVTLIFVLYRDTPvvkssSRELCYIILAGIFLGYVCPFTLIAkptttscylqRLLVGLSSAMCYSALVTKTNRiaril
    4OR2_A  cntsnLGVVAPLGYNGLLIMSCTYYAFKTRNVPan~fnEAKYIAFTMYTTCIIWLAFVPIYFgsnyk~iittCFAVSLSVTVALGCMFTPKMYiiiak

F   4JKV_A  hqdMHSYIAAFg~AVTGLCTLFTLATFVAdwr~~~~~~~~~~~~~~~~~~~~~~nsnRYPAVILFYVNACFFVGSIGWLAQFMdgarreivcradg~~~~~~~~~~~~tmrlgeptsnetlSCVIIFviVYYALMAGVVWFVVlty
    4JKV_A  gykNYRYRAGFvlAPIGLVLIVGGYFLIRgvmtlfsiksnhpgllsekaaskinetmLRLGIFGFLAFGFVLITFSCHFYDFFnqaewersfrdyvlcqanvtiglptkqpipdceiknrpSLLVEKi~NLFAMFGTGIAMSTwvw

Aa  4MQT_A  evvfIVLVAGSLSLVTIIGNILVMVSIKVNrhlqtv~~~~~~~~~~~~~~~~~~~~~~~~~~~~~~~~~nnyflFSLACADLIIGVFSMNLytlytvigywplgpvvcdlWLALDYVVSNASVMNLLIIsfdry
    4MQT_A  ffsnAAVTFGTAIAAFYLPVIIMTVLYWHIsrasksrikkdkkepvanqdpvstrkkpppsrekkvtrtiLAILLAFIITWAPYNVMvlintfca~~~~pcipntvWTIGYWLCYINSTINPACYalcna

    4MQT_A  evvfIVLVAGSLSLVTIIGNILVMVSIKVNrhlqtvn~~~~~~~~~~~~~~~~~~~~~~~~~~~~~~~~nyflfslACADLIIGVFSMNlytlytvigywplgpvvcdlwlALDYVVSNASVMNLLIIsfdry
    4MQT_A  naavTFGTAIAAFYLPVIIMTVLYWHISRAsksrikkdkkepvanqdpvstrkkpppsrekkvtrtilaiLLAFIITWAPYNVmvlintfcapc~ipntvwtigyWLCYINSTINPACYALCnatfkl

A   5U09_A  qqlaiAVLSLTLGTFTVLENLLVLCVILHSrsl~   ~rcrPSYHFIGSLAVADLLGSVIFVYSFIDFHVFhrk~dsRNVFLFKLGGVTASFTASVGSLFLAaidry
    5U09_A  fphidETYLMFWIGVTSVLLLFIVYAYMYIlwka   rmdiRLAKTLVLILVVLIICWGPLLAIMVYDVFGkmnkliKTVFAFCSMLCLLNSTVNPIIYALRskdlr

A   4U15_A  vvfIAFLTGFLALV~TIIGNILVIVAFKVnkq~   ~lkTVNNYFLLsl~acADLIIGVismnlfttyiimnrwalgnlacDLWLSIDYVasNASVMNLLVIsfdry
    4U15_A  flsEPTITFGTAIAaFYMPVTIMTILYWRiyke   kkaAQTLSAILlafiiTWTPYNImvlvntf~~~~~~~~cdscipkTYWNLGYWLc~YINSTVNPVCyalcn

A   4MBS_A  yqfwkNFQTLKIVILGLVLPLLVMVIcysgilk   rdvrliFTIMIVYFLFWAPYNIVLLLNtfqeffglnncsssNRLDQAMQVTETLGMTH~CCINPIIyafvg
    4MBS_A  aarllPPLYSLVFIFGFVGNMLVILIlinykr~   ~lksmtDIYLLNLAISDLFFLLTVPFWahyaaa~~~qwdfgNTMCQLLTGLYFIGFFSgIFFIILLtidry

A   4IB4_A  tcvltkerfGDFMLFGSLAAFFTPLAIMIVTYFltiha   raskVLGIVFFLFLLMWCPFFItnitlvl~~~~~~cdscnqTTLQMLLEIFVWIGYVSSGVNPLVYTLFnkt
    4IB4_A  eeqgnklhwAALLILMVIIPTIGGNTLVILAVSlekk~   ~lqyATNYFLMSLAVADLLVGLfvmpialltimfeamwplpLVLCPAWLFLDVLFSTASIWHLCAISVDryi

A   4GRV_A  ysKVLVTAIYLALFVVGTVGNSVTLFTLARKksl~   ~qsLQSTVHYHLGSLALSDLLILLLAMPVELYNfiwvh~~hpwafGDAGCRgYYFLRDACTYATALNVASLSVAry
    4GRV_A  tvKVVIQVNTFMSFLFPMLVISILNTVIANKltvm   svqALRHGVLVARAVVIAFVVCWLPYHVRRLMFcyisdeqwttflFDFYHY~FYMLTNALAYASSAINPILYNLvs
```

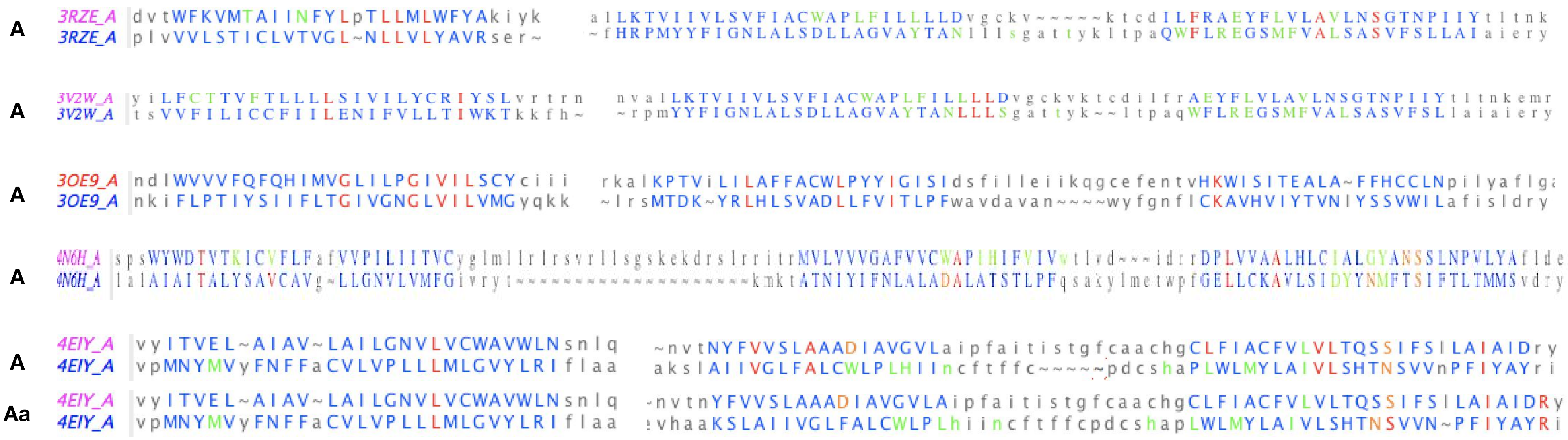

**Figure S3.A - Protodomains pairwise alignment (TM123 vs. TM567) of GPCRs.**

(Class A) Rhodopsin - 1F88: 3.24 Å; (A) Rhodopsin active 2X72: 2.59 Å; Rhodopsin active 6CMO: 3.26 Å; (B) 5EE7: 2.45 Å, (C) 4OR2: 3.36 Å, (F) 4JKV: 2.45 Å; (A) active 4MQT: 2.73 Å - human M2 muscarinic acetylcholine receptor; (A) 4U15: 2.6 Å M3 muscarinic receptor bound to tiotropium; (A) 4MBS: 2.62 Å CCR5 chemokine receptor-HIV entry inhibitor maraviroc complex; (A) 4IB4: 2.71 Å; (A) 4GRV: 3.31 Å; (A) 3RZE: 3.01 Å, (A) 3V2W: 2.73 Å (FD), alternate 2.68 Å (WD) ; (A) 3OE9: 3.38 Å; (A) 4N6H: 2.87 Å human delta-opioid receptor (delta-OR) bound to Sodium (orange) and Naltrindole (green); (A) 4EIY: 2.68 Å (WD), alternate 2.89 Å (FD). Conserved residues are colored in red. Ligand binding/proximity (less than 4 Å) colored in green; Sodium binding residues colored in orange. (FD) means the alignment matches F6.44 with D2.50. (WD) matches W6.48 with D2.50. In most cases both alignments can be performed, as a rigid body translation along the symmetry axis of one protodomain vs. the other, in some cases TMH2/6 translation alone. Examples 3V2W and 4EIY are two cases where RMSD is low and similar for both alignments. In the case of Active vs. Inactive Rhodopsin, and alignment of protodomains gets them all within the same RMSD range: 6CMO protodomain 1 vs. 1F88 protodomain 1 : RMSD=1.78 Å; protodomain 2 RMSD=2.88 Å.

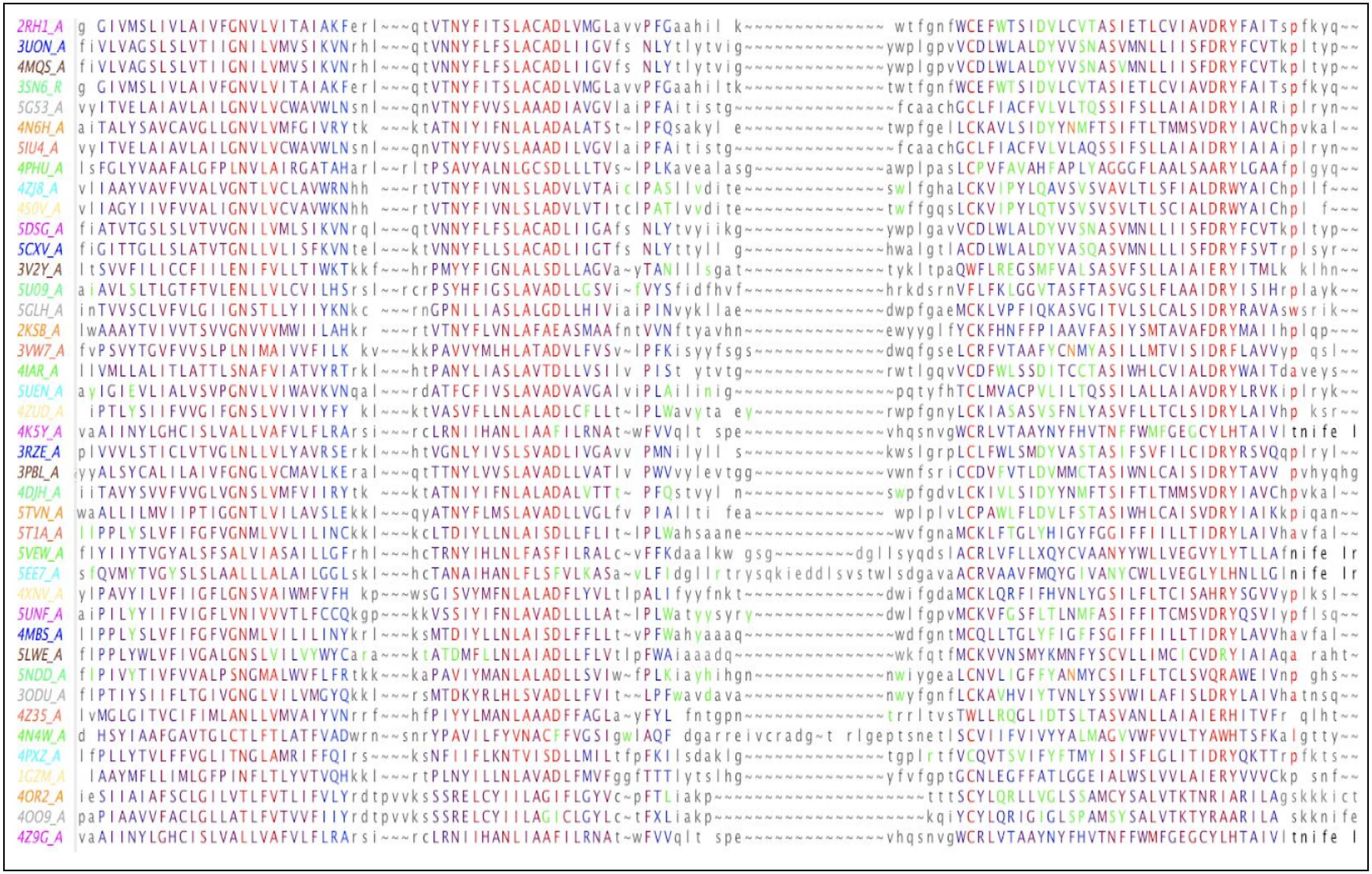
2RH1_A
3UON_A
4MQS_A
3SN6_R
5G53_A
4N6H_A
5IU4_A
4PHU_A
4ZJ8_A
4S0V_A
5DSG_A
5CXV_A
3V2Y_A
5U09_A
5GLH_A
2KSB_A
3VW7_A
4IAR_A
5UEN_A
4ZUD_A
4K5Y_A
3RZE_A
3PBL_A
4DJH_A
5TVN_A
5T1A_A
5VEW_A
5EE7_A
4XNV_A
5UNF_A
4MBS_A
5LWE_A
5NDD_A
3ODU_A
4Z35_A
4N4W_A
4PXZ_A
1GZM_A
4OR2_A
4OO9_A
4Z9G_A

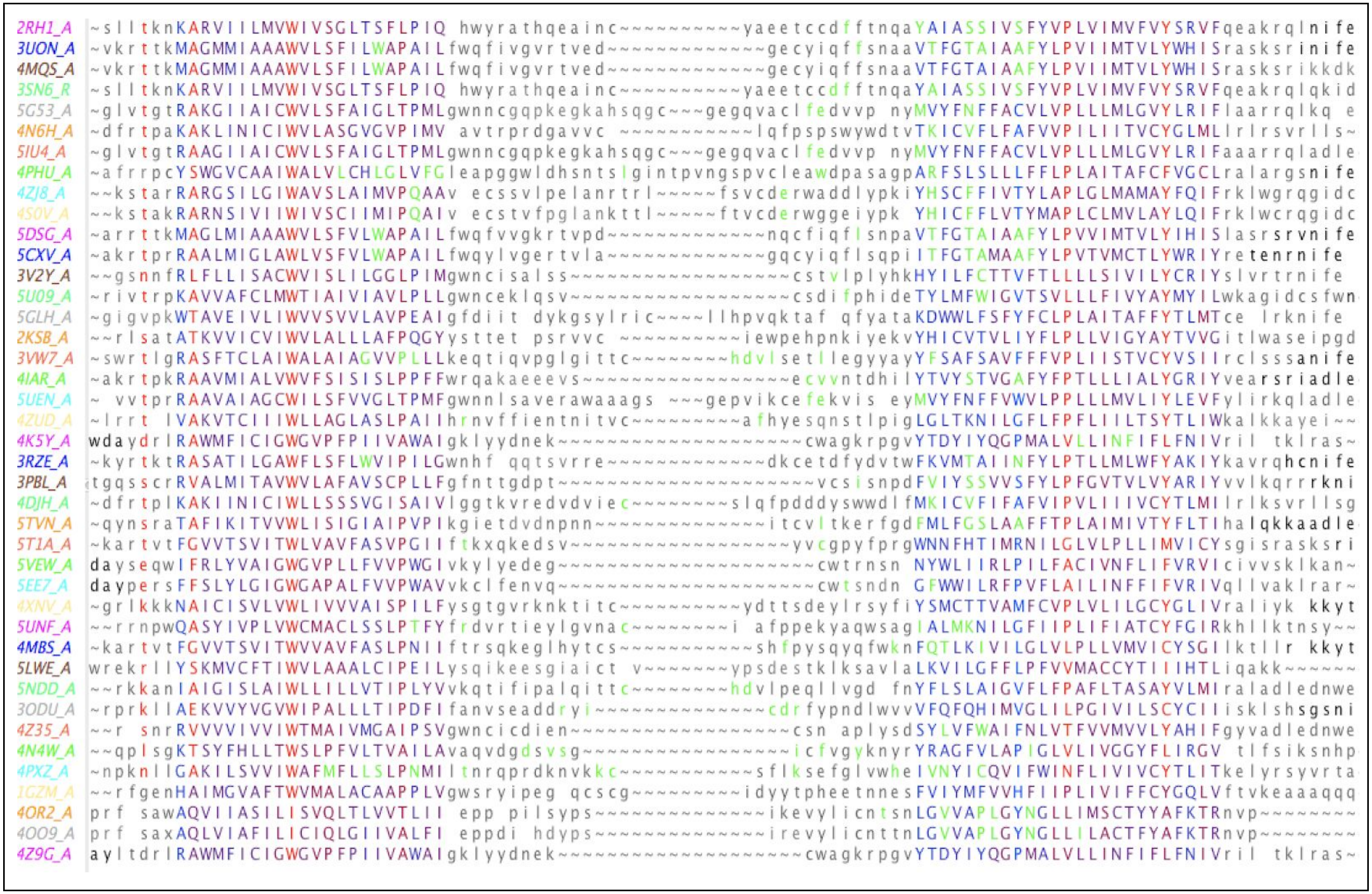
2RH1_A
3UON_A
4MQS_A
3SN6_R
5G53_A
4N6H_A
5IU4_A
4PHU_A
4ZJ8_A
4S0V_A
5DSG_A
5CXV_A
3V2Y_A
5U09_A
5GLH_A
2KSB_A
3VW7_A
4IAR_A
5UEN_A
4ZUD_A
4K5Y_A
3RZE_A
3PBL_A
4DJH_A
5TVN_A
5T1A_A
5VEW_A
5EE7_A
4XNV_A
5UNF_A
4MBS_A
5LWE_A
5NDD_A
3ODU_A
4Z35_A
4N4W_A
4PXZ_A
1GZM_A
4OR2_A
4OO9_A
4Z9G_A

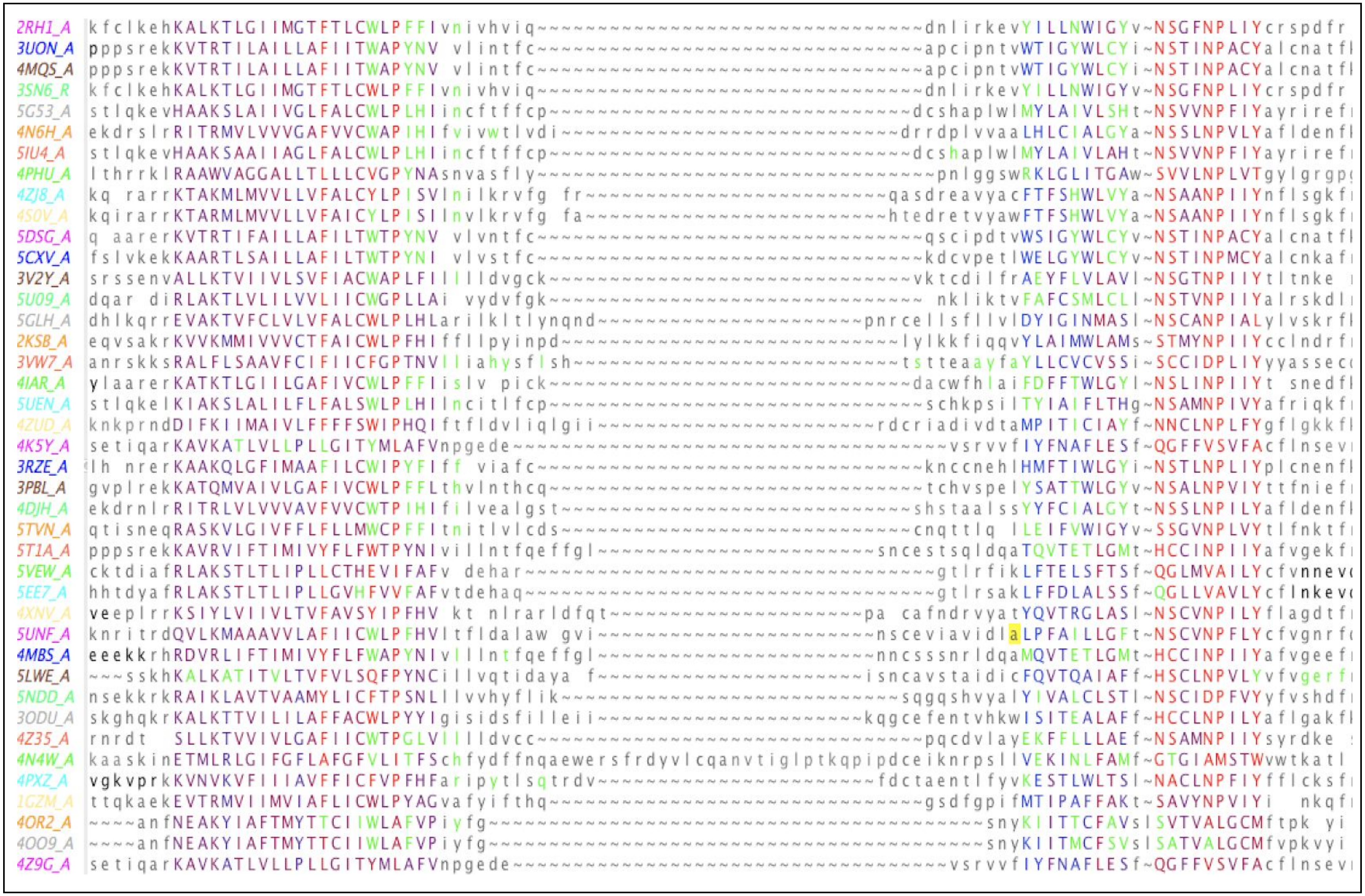

**Figure S3.B - GPCR domains TM1-TM7 multiple alignment - (mostly Class A).** Red = conserved, Green = ligand binding

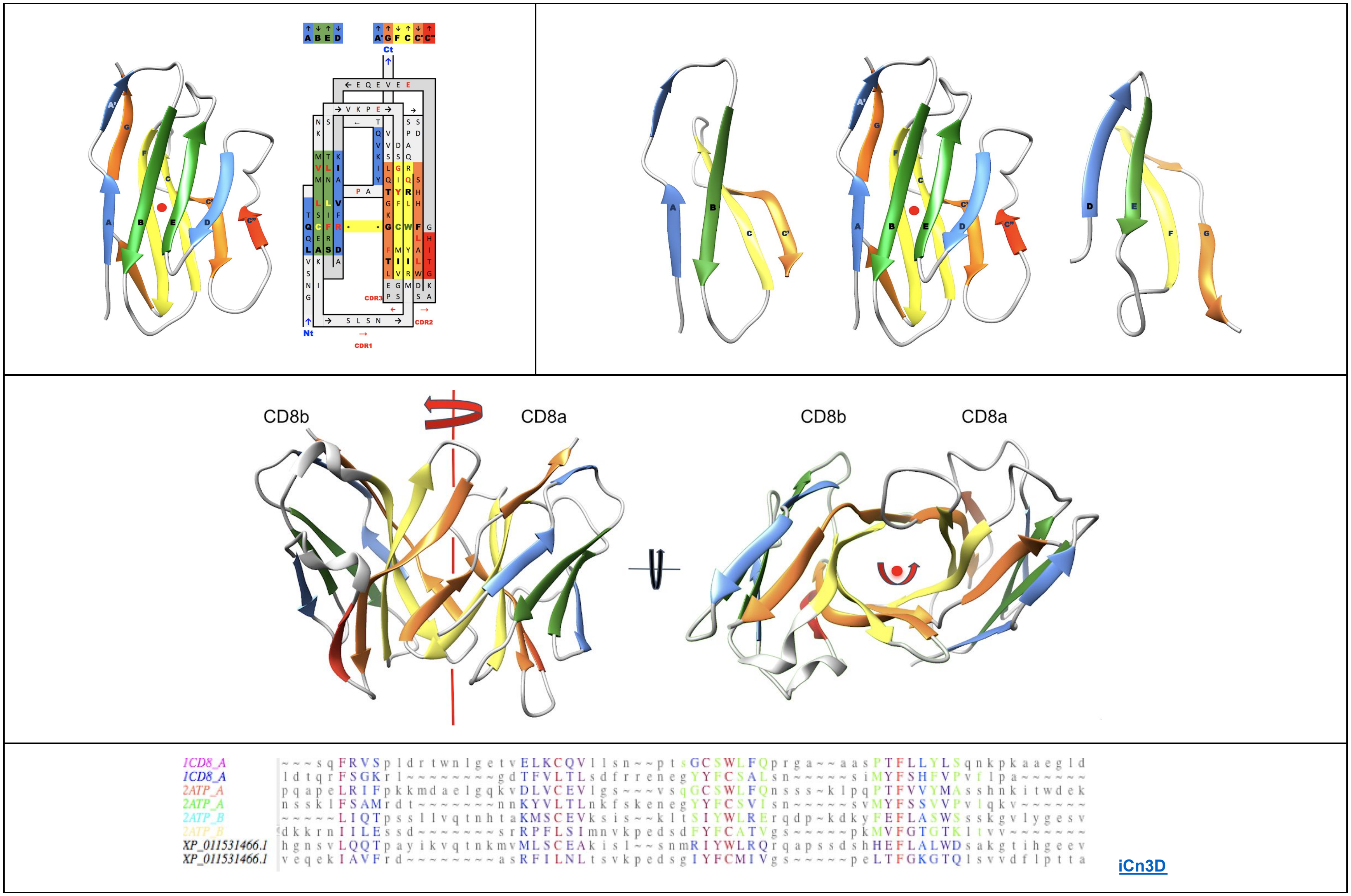

**Figure S4 - Pseudo-symmetry and Protodomains self-complementarity in Immunoglobulin domains and dimers -** Example of CD8ab.

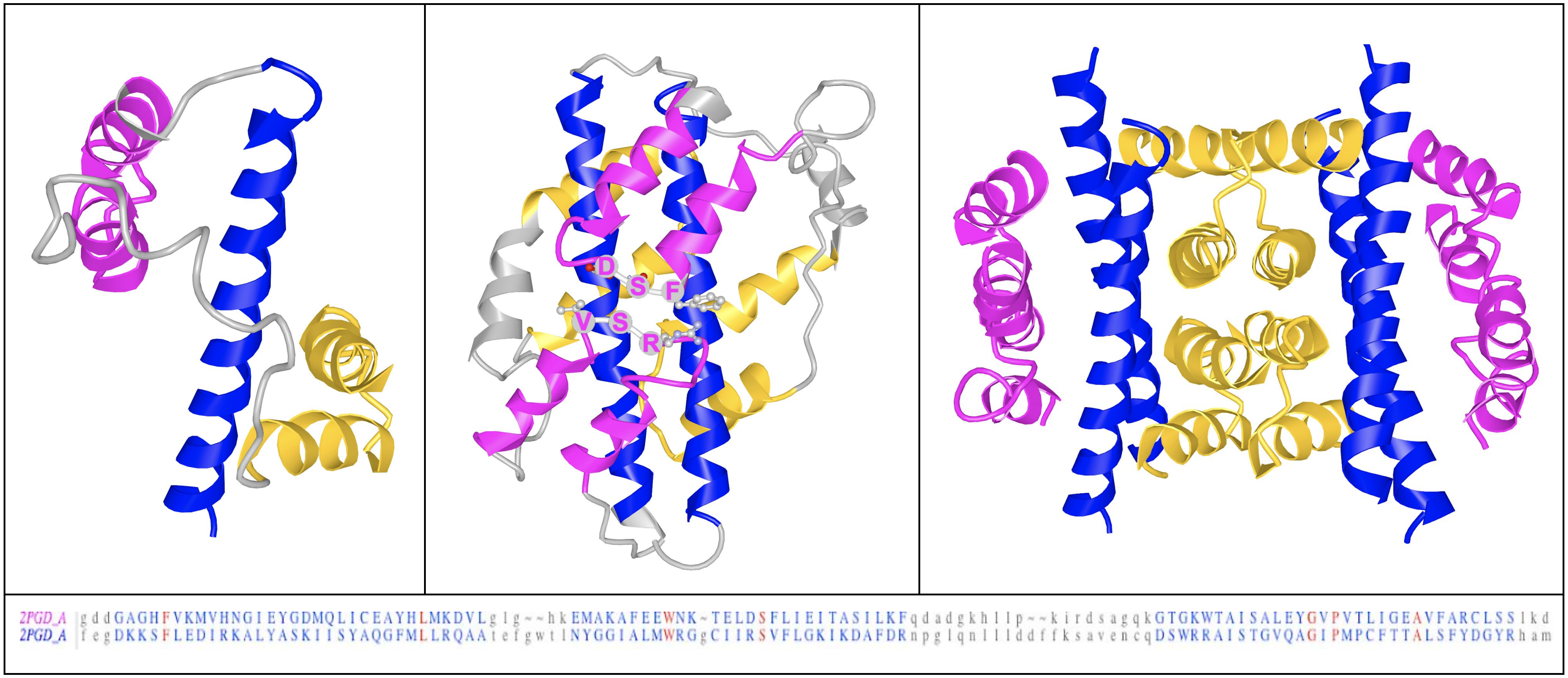

**Figure S5 - Protodomain alignment of an alpha-helical globular protein. (**3D visualization link**:** iCn3D) Example of 6-phosphogluconate dehydrogenase (6PGD) (PDB:2PGD; SCOP a.100). Protodomain (left) , domain (middle) , dimer (right) without loops and linkers for clarity. This example shows first the protodomain supersecondary structure with the first central Helix in blue, followed by a helix turn helix in magenta and another helix turn helix motif in orange. The domain is formed by 2 complementary protodomains where each elementary motif is complementary to its image blue, magenta, orange. Finally two domains dimerize through a complementary surface formed by 2x2 orange motif forming a 4x4 helical interface, where each individual helix is paired with its image. This structure exhibits multiple local self-complementary motifs. Note that the pairing of magenta helix turn helix motifs is performed by the turn motif labeled in the central picture, with the DSF and RSV structurally homologous at the backbone level. Such a motif would correspond in membrane proteins with “reentrant” helical motifs, that may resemble the magenta pairing pattern.

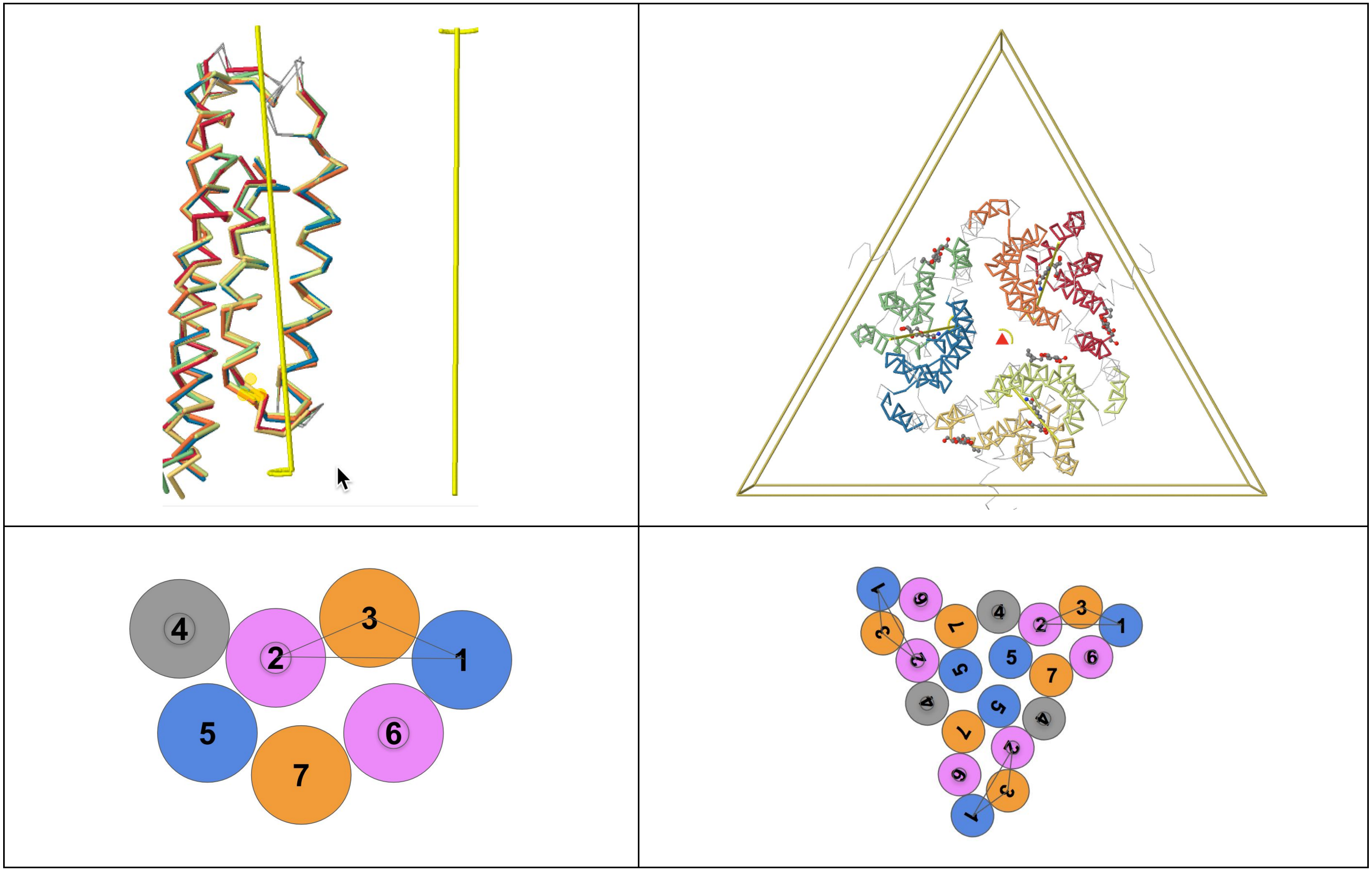

**Figure S6 - Protodomains, Domain Pseudo-symmetry and Quaternary symmetry - SWEET** Protodomain topology parallel C2 132c/s21a

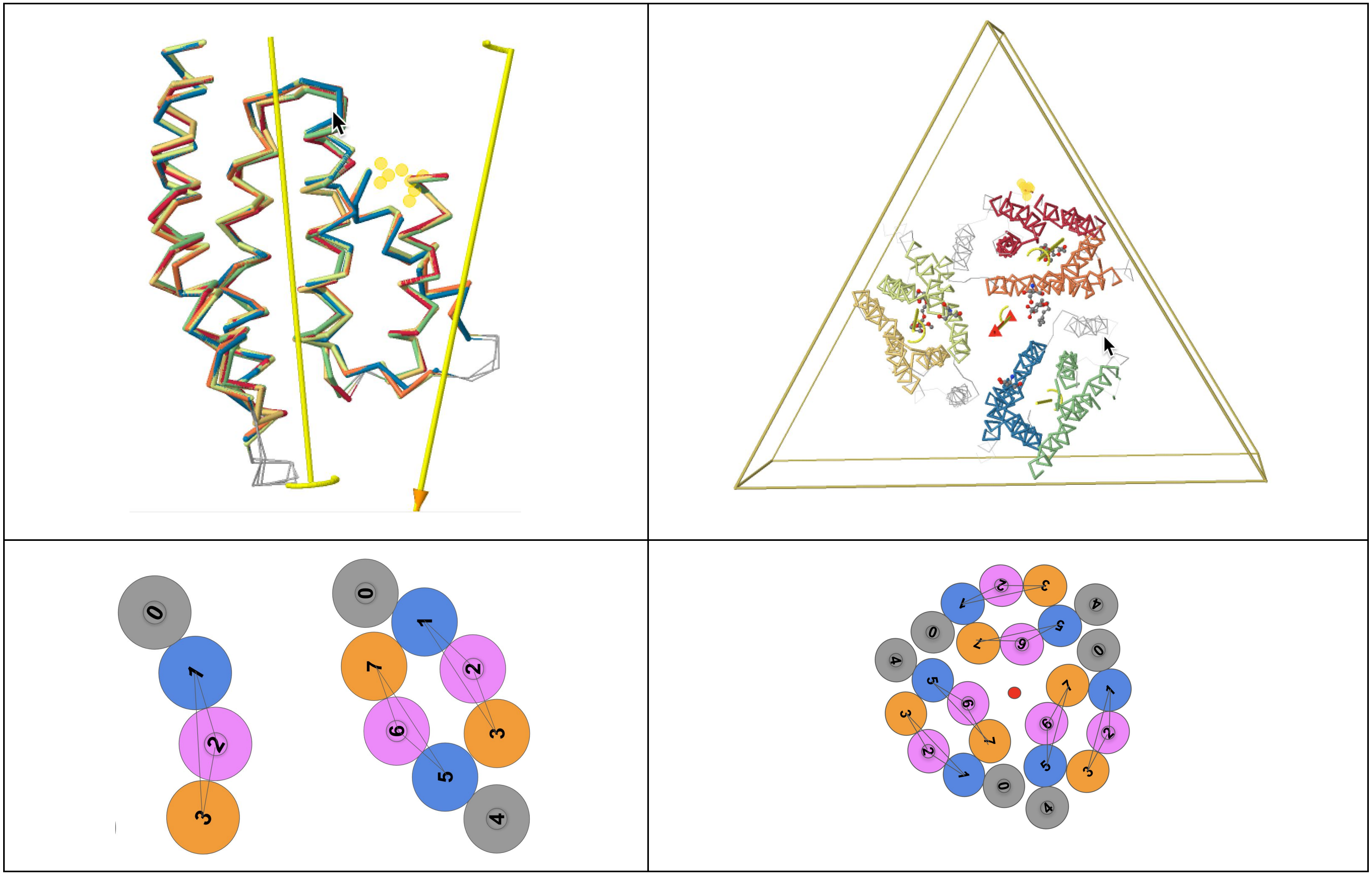

**Figure S7 - Protodomains, Domain Pseudo-symmetry and Quaternary symmetry - PnuC** Protodomain topology parallel C2 123c/s31p

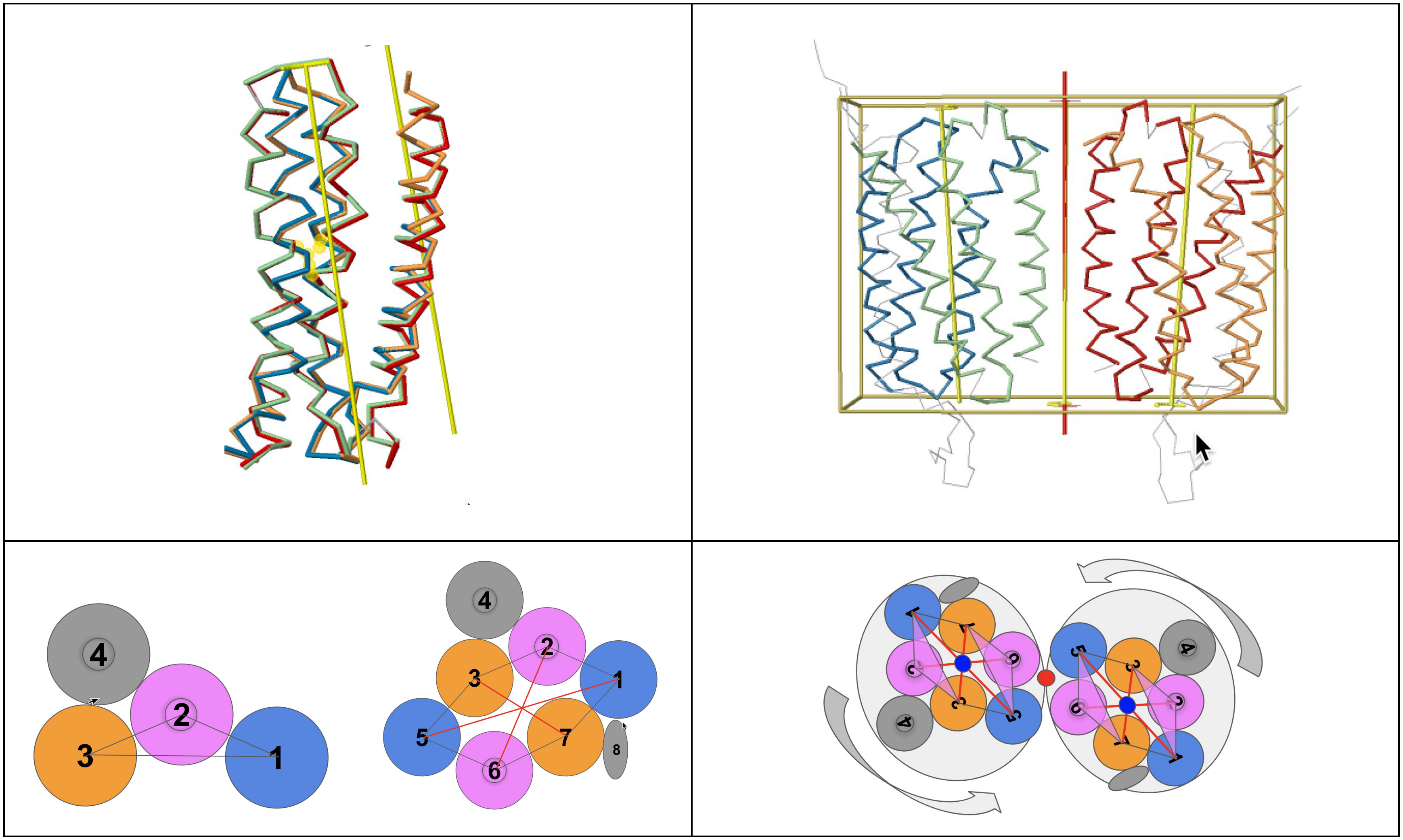

**Figure S8 - Protodomains, Domain Pseudo-symmetry and Quaternary symmetry -** GPCR Protodomain topology parallel C2 123cc/s31p.

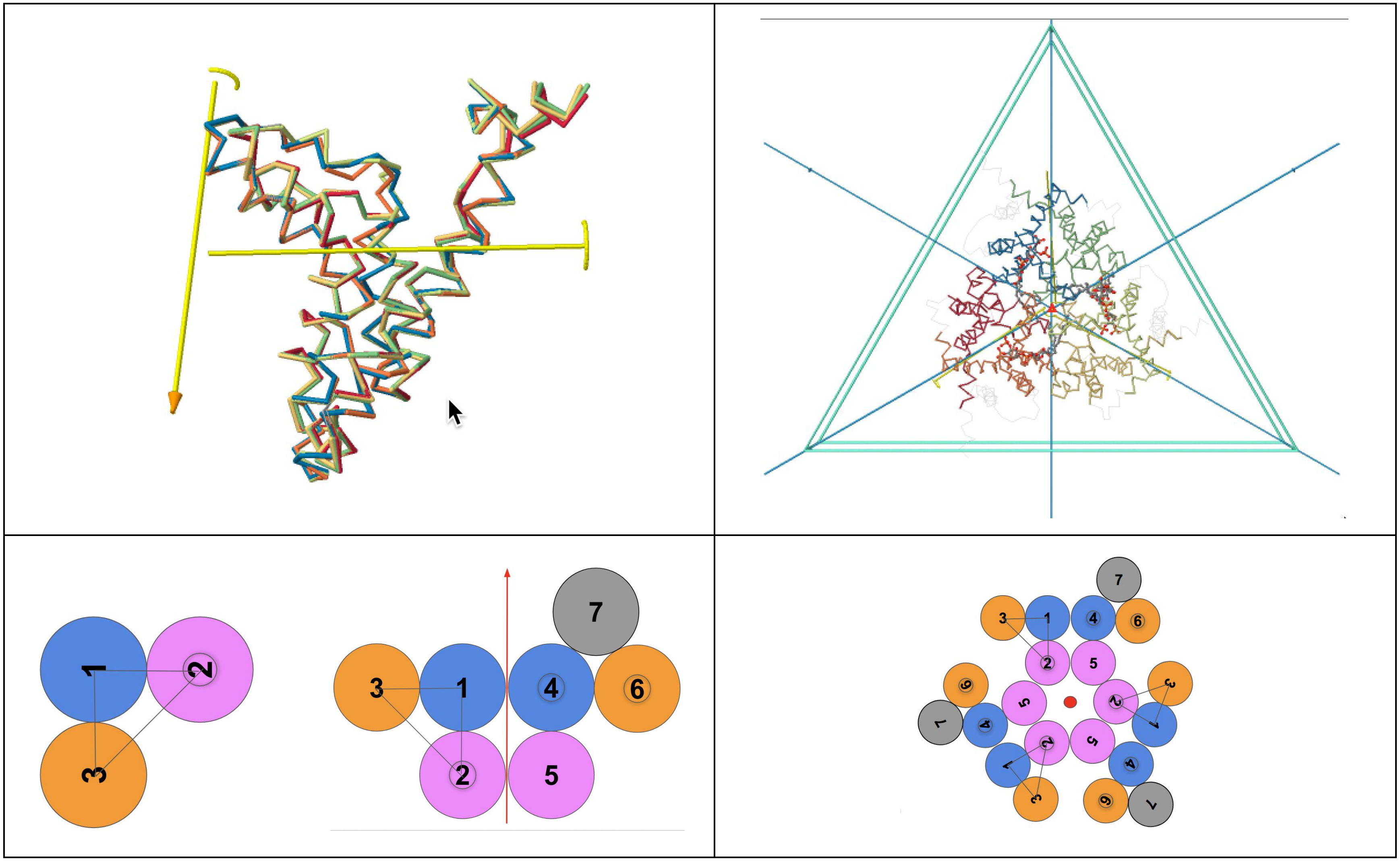

**Figure S9 - Protodomains, Domain Pseudo-symmetry and Quaternary symmetry -** TRIC Protodomain topology inverted 312c/s11a22a

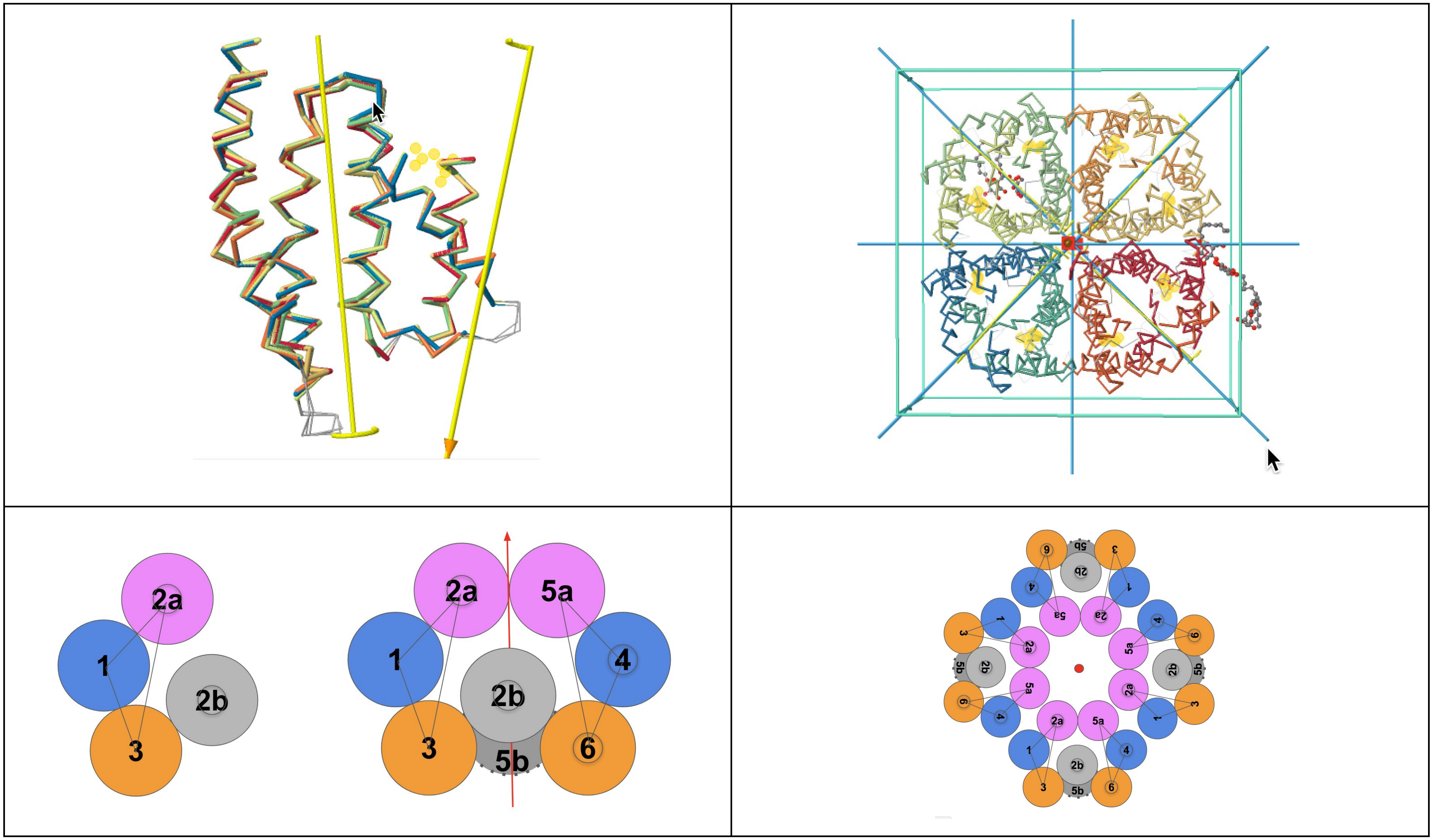

**Figure S10 - Protodomains, Domain Pseudo-symmetry and Quaternary symmetry -** AQP1 Protodomain topology inverted C2 312cc/a22a [forms a tetramer quaternary structure, FocA uses the same protodomain and forms a pentamer]. This is the only example with an asymmetric (a) 22 interface, of a very peculiar and idiosyncratic nature: helix 2 splits in 2 forming 2a and 2b to with a helix 2a-2a antiparallel interface from each protodomain on one side and 2b-2b on the other where the latter stack on top of each other one going up the other down - [iCn3D 3D visualization]

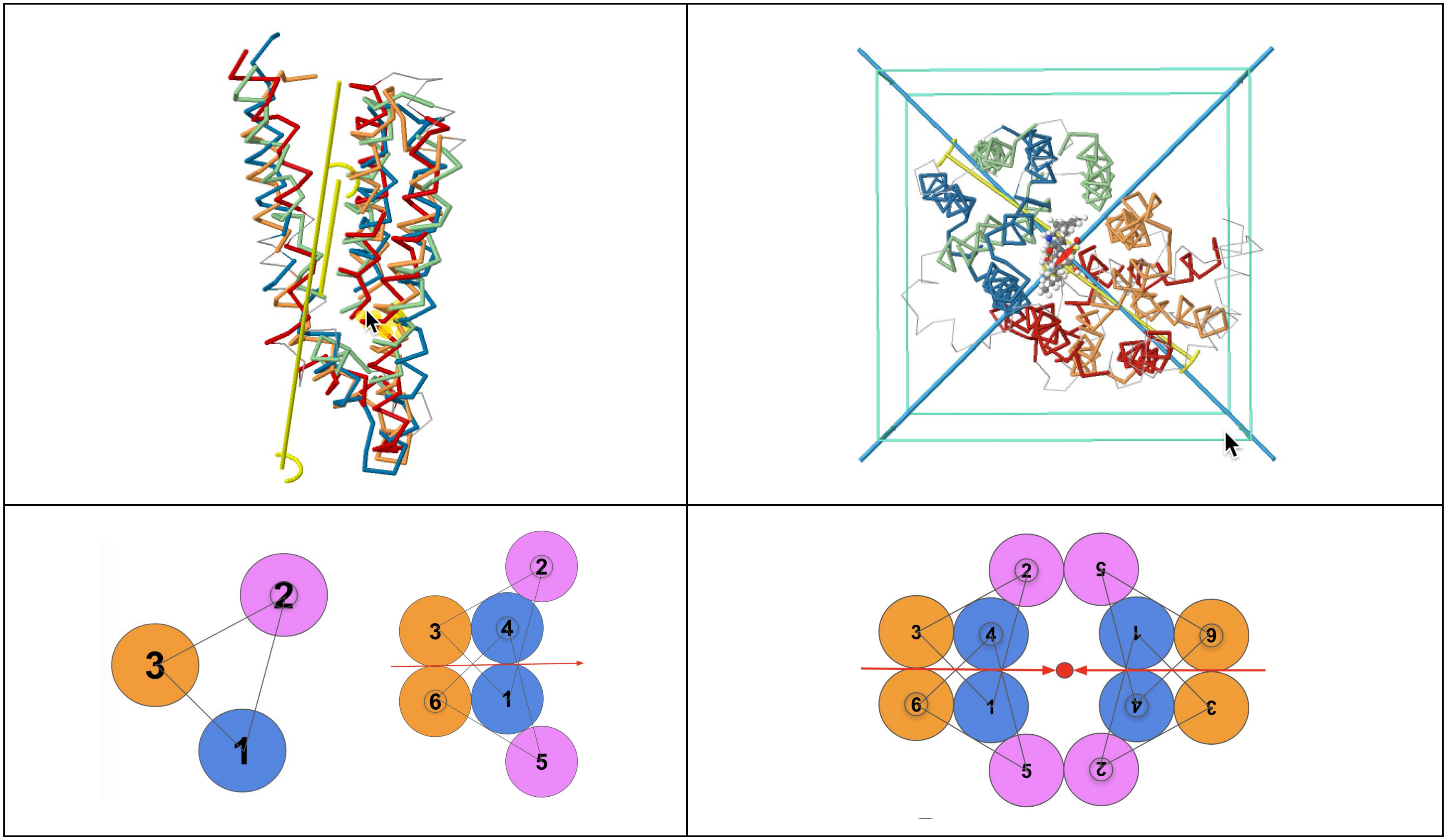

**Figure S11 - Protodomains, Domain Pseudo-symmetry and Quaternary symmetry -** MFS Protodomain topology inverted C2 interdigitated 132cc/s11a33a MFS - **A)** Inverted Interdigitated Protodomains forming a C2 symmetric domain. It possesses two locally symmetric interfaces TM1-TM1 and TM3-TM3 (s11a33a) **B)** C2 symmetric domain packing through a double (symmetry related) TM2-TM2 interface (2*s22a) formed by the second helix of each of the 4 protodomains. The 2-domains/4-protodomains MFS protein has an overall D2 symmetry. (PDB: 5EQI) since the internal symmetry axes of each domain colinearize and are orthogonal to the central axis (perpendicular to the membrane planes).

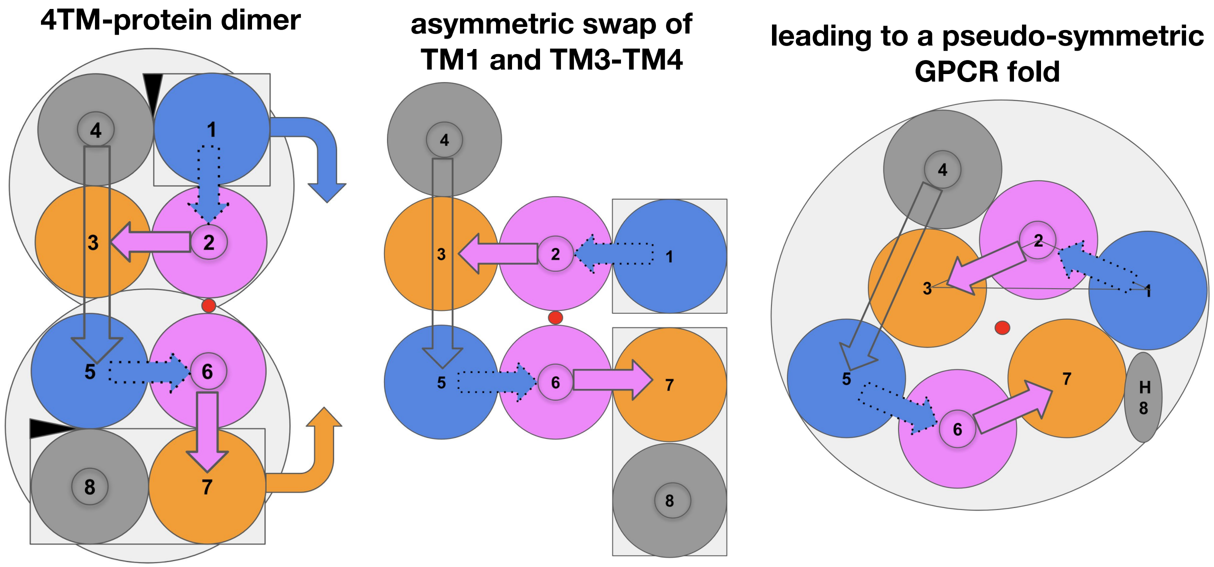

**Figure S12 - Concerted asymmetric subdomain swap TM3-TM4 vs. TM1: A rearrangement scenario** involving a rigid swap at the interface TM1-TM4 of both 4TMH domains forming a dimer under a TM4(1)-TM1(2) linker constraint. This would involve a conformational change and transition from a dimer of 4TM-protein binding G-proteins to a GPCR configuration through an asymmetric swap of one helix TM1 on one monomer vs. a two helices TM3-TM4 on the second to obtain a symmetric GPCR arrangement of TM1-2-3 vs TM5-6-7.

**6CMO_R:** chimera protein of Soluble cytochrome b562 and Rhodopsin
**1GZM_A:** RHODOPSIN

```
                    10        20        30        40        50        60
               ....*....|....*....|....*....|....*....|....*....|....*....|
6CMO_R 142 CgtegpnfyvpFSNAtgvvrspfeypqyYLAEPWQFSmlaaymfllivlgfpinfltlyv 201
1GZM_A   3 NgtegpnfyvpFSNKtgvvrspfeapqyYLAEPWQFSmlaaymfllimlgfpinfltlyv 62

                    70        80        90       100       110       120
               ....*....|....*....|....*....|....*....|....*....|....*....|
6CMO_R 202 tvqhkklrtplnyILLNLAVADLFMVLGGFTSTLYTSlhgyfvfgptgcnlqGFFATLGG 261
1GZM_A  63 tvqhkklrtplnyILLNLAVADLFMVFGGFTTTLYTSlhgyfvfgptgcnleGFFATLGG 122

                   130       140       150       160       170       180
               ....*....|....*....|....*....|....*....|....*....|....*....|
6CMO_R 262 EIALWSLVVLAIERYVVvckpmsnfrfGENHAIMGVAFTWvmalacaapplagwsryipe 321
1GZM_A 123 EIALWSLVVLAIERYVVvckpmsnfrfGENHAIMGVAFTWvmalacaapplvgwsryipe 182

                   190       200       210       220       230       240
               ....*....|....*....|....*....|....*....|....*....|....*....|
6CMO_R 322 glqcscgidyytlkpevnnesfviymfvvhftIPMIIIFFCYGQLVFTVKEaaaqqqesa 381
1GZM_A 183 gmqcscgidyytpheetnnesfviymfvvhfiIPLIVIFFCYGQLVFTVKEaaaqqqesa 242

                   250       260       270       280       290       300
               ....*....|....*....|....*....|....*....|....*....|....*....|
6CMO_R 382 ttqkaekevtrmviiyviaflicwvpyasvafyifthqgsCFgpifmTIPAFFAKsaaiy 441
1GZM_A 243 ttqkaekevtrmviimviaflicwlpyagvafyifthqgsDFgpifmTIPAFFAKtsavy 302

               ....*....
6CMO_R 442 npviyimmN 450
1GZM_A 303 npviyimmN 311
```

**Figure S13** - VAST+ Invariant substructure alignment between the active (6CMO, human) vs, inactive (1GZM bovine) conformations of Rhodopsin - RMSD = 1.70 Å for 106 residues aligned - 86% id . 3D visualization: iCn3D.

| Aquaporin | TM1 | TM2 a | TM2 b | TM3 | Protodomain TM1-TM2-TM3 | TM4 | TM5 a | TM5 b | TM6 | Protodomain TM4-TM5-TM6 |
|---|---|---|---|---|---|---|---|---|---|---|
| EC | 40 | 44 | - | 42 | 42 | 45 | 36 | 57 | 57 | 49 |
| IC | 46 | 44 | 57 | 48 | 49 | 42 | 51 | - | 45 | 46 |
| Full | 43 | 44 | 57 | 45 | 47 | 43 | 42 | 57 | 50 | 48 |

| Foca | TM1 | TM2 a | TM2 b | TM3 | Protodomain TM1-TM2-TM3 | TM4 | TM5 a | TM5 b | TM6 | Protodomain TM4-TM5-TM6 |
|---|---|---|---|---|---|---|---|---|---|---|
| EC | 28 | 38 | - | 36 | 34 | 39 | 34 | 39 | 33 | 36 |
| IC | 40 | 52 | 41 | 43 | 44 | 44 | 42 | - | 52 | 46 |
| Full | 34 | 44 | 41 | 40 | 40 | 42 | 38 | 39 | 43 | 41 |

| PnuC | TM0 | TM1 | TM2 | TM3 | Protodomain TM1-TM2-TM3 | TM4 | TM5 | TM6 | TM7 | Protodomain TM5-TM6-TM7 |
|---|---|---|---|---|---|---|---|---|---|---|
| EC | 28 | 50 | 48 | 34 | 44 | 32 | 46 | 46 | 33 | 42 |
| IC | 28 | 43 | 58 | 46 | 49 | 32 | 57 | 61 | 31 | 50 |
| Full | 28 | 46 | 51 | 39 | 46 | 32 | 50 | 51 | 32 | 44 |

| Tric | TM1 | TM2 | TM3 | Protodomain TM1-TM2-TM3 | TM4 | TM5 | TM6 | TM7 | Protodomain TM4-TM5-TM6 |
|---|---|---|---|---|---|---|---|---|---|
| EC | 38 | 57 | 40 | 45 | 41 | 47 | 36 | 28 | 41 |
| IC | 41 | 48 | 68 | 52 | 40 | 42 | 44 | 27 | 42 |
| Full | 40 | 53 | 54 | 49 | 41 | 44 | 39 | 28 | 41 |

| MFS | TM1 | TM2 | TM3 | TM4 | TM5 | TM6 | TM7 | TM8 | TM9 | TM10 | TM11 | TM12 |
|---|---|---|---|---|---|---|---|---|---|---|---|---|
| EC | 24 | 25 | 28 | 35 | 29 | 31 | 25 | 18 | 23 | 21 | 24 | 20 |
| IC | 22 | 41 | 35 | 34 | 28 | 30 | 22 | 26 | 21 | 16 | 21 | 23 |
| Full | 23 | 33 | 31 | 35 | 29 | 30 | 24 | 22 | 22 | 18 | 23 | 22 |
| | Proto 1 | | 29 | Proto 2 | | 31 | Proto 3 | | 23 | Proto 4 | | 21 |

**Figure S14: Sequence Divergence of TMH Families -** Sequence similarity score (see Methods section for details) of the aligned EC half, IC half, and Full TM sequences for each of the TMs **TMH families Aquaporin, Foca, PnuC, Tric, and MFS.** Protodomain 1 and 2 scores are given along with those for EC-facing and IC-facing halves. Higher numbers mean high sequence similarity (or higher conservation), where a maximum score of 100 would mean identical sequences or two sequences with similar residues at each position in the sequence alignment. Protodomains are specific to each protein family, shown in Table II in the main text and are discussed in the main text, and corresponding Supplement Figures. The list of proteins and PDB ids used for each family is provided in the Supplement File SF1. We use here a common 3TMH protodomain decomposition, including for PnuC (in the main text we used 4TMH)
