## Supplementary material for "7-Transmembrane Helical (7TMH) Proteins: Pseudo-Symmetry and Conformational Plasticity"

**GPCRs**

| PDB | Uniprot | use |
|---|---|---|
| 4PHU | O14842 | + |
| 4ZJC | O43613 | + |
| 2RH1 | P07550 | + |
| 4ZWJ | P08100 | + |
| 3UON | P08172 | + |
| 5DSG | P08173 | + |
| 5XCV | P11229 | + |
| 4XES | P20789 | + |
| 3V2Y | P21453 | + |
| 5U09 | P21554 | + |
| 5GLH | P24530 | + |
| 2LNL | P25024 | + |
| 3VW7 | P25116 | + |
| 4IAR | P28222 | + |
| 5EML | P29274 | + |
| 5UEN | P30542 | + |
| 4K5Y | P34998 | + |
| 3RZE | P35367 | + |
| 3PBL | P35462 | + |
| 4N6H | P41143 | + |
| 4DJH | P41145 | + |
| 4EA3 | P41146 | + |
| 4OO9 | P41594 | + |
| 5T1A | P41597 | + |
| 5VEW | P43220 | + |
| 5EE7 | P47871 | + |
| 4XNV | P47900 | + |
| 5UNG | P50052 | + |
| 4MBS | P51681 | + |
| 5LWE | P51686 | + |
| 5NDD | P55085 | + |
| 3ODU | P61073 | + |
| 4OR2 | Q13255 | + |
| 4Z34 | Q92633 | + |
| 4JKV | Q99835 | + |
| 4PXZ | Q9H244 | + |

**CDD Link**

NONE, S2Table.csv was used from (Cvicek et al. 2016):
http://www.csun.edu/gpcrs/datas/gross_paper/S2Table.csv

**Aquaporins**

| PDB | Uniprot | Use | gi # |
|---|---|---|---|
| 1FQY | P29972 | - | 14278358 |
| 1FX8 | P0AER0 | - | 21466058 |
| 1H6I | P29972 | + | 14278358 |
| 1IH5 | P29972 | - | 14278358 |
| 1J4N | P47865 | - | NONE |
| 1LDA | P0AER0 | - | 21466058 |
| 1LDF | P0AER0 | + | 21466058 |
| 1LDI | P0AER0 | - | 21466058 |
| 1RC2 | P60844 | - | NONE |
| 1SOR | Q6J8I9 | - | NONE |
| 1YMG | P06624 | - | NONE |
| 1Z98 | Q41372 | - | NONE |
| 2ABM | P60844 | - | NONE |
| 2B5F | Q41372 | - | NONE |
| 2B6O | Q6J8I9 | + | NONE |
| 2B6P | Q6J8I9 | - | NONE |
| 2C32 | P06624 | - | NONE |
| 2D57 | P47863 | - | NONE |
| 2EVU | Q9C4Z5 | - | NONE |
| 2F2B | Q9C4Z5 | + | NONE |
| 2O9D | P60844 | - | NONE |
| 2O9E | P60844 | - | NONE |
| 2O9F | P60844 | - | NONE |
| 2O9G | P60844 | + | NONE |
| 2W1P | F2QVG4 | - | NONE |
| 2W2E | F2QVG4 | - | NONE |
| 2ZZ9 | P47863 | - | NONE |
| 3C02 | Q8WPZ6 | + | NONE |
| 3CLL | Q41372 | - | NONE |
| 3CN5 | Q41372 | + | NONE |
| 3CN6 | Q41372 | - | NONE |
| 3D9S | P55064 | + | NONE |
| 3GD8 | P55087 | + | NONE |
| 3IYZ | P47863 | - | NONE |
| 3J41 | Q6J8I9 | - | NONE |
| 3LLQ | Q8UJW4 | + | NONE |
| 3M9I | Q6J8I9 | - | NONE |
| 3NE2 | O28846 | + | NONE |
| 3NK5 | P60844 | - | NONE |
| 3NKA | P60844 | - | NONE |
| 3NKC | P60844 | - | NONE |
| 3ZOJ | F2QVG4 | + | NONE |
| 4CSK | P29972 | - | 14278358 |
| 4IA4 | Q41372 | - | NONE |
| 4JC6 | Q41372 | - | NONE |

| | | | |
|---|---|---|---|
| 4NEF | P41181 | + | 618855042 |
| 4OJ2 | P41181 | - | 618855042 |
| 5BN2 | F2QVG4 | - | NONE |
| 5C5X | P55064 | - | NONE |
| 5DYE | P55064 | - | NONE |
| 5I32 | Q41951 | + | NONE |
| NONE | P23645 | + | 71153495 |
| NONE | P23900 | + | 1706896 |
| NONE | P06624 | + | 85544350 |
| NONE | Q08451 | + | 586102 |
| NONE | P43286 | + | 1175013 |
| NONE | P25818 | + | 138560 |
| NONE | P26587 | + | 135858 |
| NONE | P08995 | + | 1352509 |
| NONE | O24389 | + | 1518057 |

**CDD Link:**
https://www.ncbi.nlm.nih.gov/Structure/cdd/cddsrv.cgi?uid=321252

**FOCA**

| **PDB** | **Uniprot** | **Use** | **gi#** |
|---|---|---|---|
| 4FC4 | E8XEH9 | + | NONE |
| 3TDP | Q186B7 | + | NONE |
| 3Q7K | Q7CQU0 | + | NONE |
| 3KLY | Q9KRE7 | + | NONE |
| 3KCU | P0AC25 | + | NONE |
| NONE | P38750 | + | 731592 |
| NONE | Q8XCN1 | + | 156354448 |
| NONE | 0A1Q4GXT | + | 15927981 |
| NONE | Q8DPM4 | + | 15901077 |
| NONE | Q8ZNA4 | + | 16765720 |
| NONE | P37327 | + | 586613 |
| NONE | Q92E59 | + | 16799677 |
| NONE | W8U1Z5 | + | 15926006 |

**CDD Link**

https://www.ncbi.nlm.nih.gov/Structure/cdd/cddsrv.cgi?uid=COG2116&islf=1

**PNuC**

| PDB | Uniprot | Use | gi # |
|---|---|---|---|
| 4QTN | D2ZZC1 | + | NONE |
| NONE | D6ZNI7 | + | 15901687 |
| NONE | Q8NU75 | + | 19551314 |
| NONE | Q8X953 | + | 15830033 |
| NONE | O25877 | + | 15645903 |
| NONE | Q9CH61 | + | 15672860 |
| NONE | Q9I2E6 | + | 15597154 |
| NONE | Q9CK00 | + | 15603703 |
| NONE | P24520 | + | 20141705 |
| NONE | D3QMT7 | + | 15800460 |
| NONE | Q9ZJT8 | + | 15612275 |
| NONE | P0AFK2 | + | 2507102 |

CDD Link
https://www.ncbi.nlm.nih.gov/Structure/cdd/cddsrv.cgi?uid=COG3201&islf=1

**TRiC**

| PDB | Uniprot | Use | gi # | |
|---|---|---|---|---|
| 5WTR | Q981D4 | + | NONE | |
| 5H35 | Q981D4 | - | NONE | Not used as better resolution structure avaliable. |
| 5EIK | Q9NA73 | + | 75023742 | |
| 5EGI | Q9NA75 | + | NONE | |
| NONE | A7SYB0 | + | 156354448 | |
| NONE | A1X7VA | - | 761906102 | |
| NONE | B4L9M1 | + | 195135689 | |
| NONE | B4LI23 | + | 968115624 | |
| NONE | C3XU22 | + | 260834779 | |
| NONE | C3XU25 | + | 260834785 | |
| NONE | W5LC18 | + | 597742168 | |
| NONE | W4YLG9 | + | 390366097 | |
| NONE | B4J0T3 | + | 195011963 | |

**CDD Link**

https://www.ncbi.nlm.nih.gov/Structure/cdd/cddsrv.cgi?uid=pfam05197&islf=1

**Semi-SWEET**

| PDB | Uniprot | Use | gi # |
|---|---|---|---|
| 5UHS | B0SR19 | - | NONE |
| 5UHQ | B0SR19 | - | NONE |
| 4X5N | P0DMV3 | - | NONE |
| 4X5M | P0DMV3 | + | NONE |
| 4RNG | B5YGD6 | + | NONE |
| 4QNC | B0SR19 | + | NONE |
| NONE | Q98RJ6 | + | 15828483 |
| NONE | Q57574 | + | 2495803 |
| NONE | C3PPD0 | + | 15892952 |
| NONE | Q8DR88 | + | 15900237 |
| NONE | Q8YYQ1 | + | 14195404 |
| NONE | Q9PRA3 | + | 14195403 |
| NONE | Q9PRA4 | + | 17228290 |

**CDD Link**

https://www.ncbi.nlm.nih.gov/Structure/cdd/cddsrv.cgi?uid=COG4095&islf=1

**SWEET**

| PDB | Uniprot | Use |
|---|---|---|
| 5CTG | Q5N8J1 | + |
| 5CTH | Q5N8J1 | - |
| NONE | Q8L9J7 | + |
| NONE | Q6L568 | + |
| NONE | Q19VE6 | + |
| NONE | Q2QR07 | + |
| NONE | Q9FPN0 | + |
| NONE | Q9BRV3 | + |
| NONE | Q9XX26 | + |
| NONE | O45102 | + |
| NONE | B4NMK1 | + |
| NONE | D0N2J4 | + |
| NONE | V9FTL5 | + |
| NONE | .0A075AWI | + |
| NONE | F4NY39 | + |
| NONE | A8HVE3 | + |

**CDD Link**

https://www.ncbi.nlm.nih.gov/Structure/cdd/cddsrv.cgi?uid=pfam03083&islf=1

MFS

| PDB | Uniprot | Use | gi # |
|---|---|---|---|
| 4PYP | P11166 | - | NONE |
| 5EGQ | P11166 | - | NONE |
| 5EQH | P11166 | + | NONE |
| 5EQI | P11166 | - | NONE |
| 4GBY | P0AGF4 | + | NONE |
| 4GBZ | P0AGF4 | - | NONE |
| 4GC0 | P0AGF4 | - | NONE |
| 4JA3 | P0AGF4 | - | NONE |
| 4JA4 | P0AGF4 | - | NONE |
| 4QIQ | P0AGF4 | - | NONE |
| 3Q7P | P11551 | - | NONE |
| 3O7Q | P11551 | + | NONE |
| 1PW4 | P08194 | + | 34810882 |
| 1PV6 | P02920 | - | 34810676 |
| 2FCP | P02920 | - | 34810676 |
| 2CFQ | P02920 | + | 34810676 |
| 2V8N | P02920 | - | 34810676 |
| 2Y5Y | P02920 | - | 34810676 |
| 4ZYR | P02920 | - | 34810676 |
| 5GXB | P02920 | - | 34810676 |
| NONE | P76350 | + | 2500934 |
| NONE | P0AEX3 | + | 84029499 |
| NONE | Q5HRH0 | + | 81174989 |
| NONE | O51798 | + | 7387890 |
| NONE | P0C0L7 | + | 81171069 |
| NONE | P37643 | + | 586685 |
| NONE | P71369 | + | 7388456 |
| NONE | P94131 | + | 7387918 |

**CDD Link**

https://www.ncbi.nlm.nih.gov/Structure/cdd/cddsrv.cgi?uid=cd06174#seqalign

**ACHA7**

| PDB | Uniprot | Use | gi # |
|---|---|---|---|
| 2MAW | P36544 | - | NONE |
| 5AFH | P36544 | - | NONE |
| 5AFJ | P36544 | - | NONE |
| 5AFK | P36544 | - | NONE |
| 5AFL | P36544 | - | NONE |
| 5AFM | P36544 | - | NONE |
| 5AFN | P36544 | + | NONE |
| 4PIR | P23979 | + | 672885916 |
| NONE | P28476 | + | 410516956 |
| NONE | P24046 | + | 223590210 |
| NONE | P26714 | + | 120781 |
| NONE | P19019 | + | 120772 |
| NONE | P24045 | + | 120775 |
| NONE | P08220 | + | 120766 |
| NONE | P25123 | + | 635377460 |
| NONE | P22933 | + | 120792 |
| NONE | L8IEJ2 | + | 440904075 |

**CDD Link**

https://www.ncbi.nlm.nih.gov/Structure/cdd/cddsrv.cgi?uid=308533
