## Supplementary material for "7-Transmembrane Helical (7TMH) Proteins: Pseudo-Symmetry and Conformational Plasticity"

### ACHA7 TM1

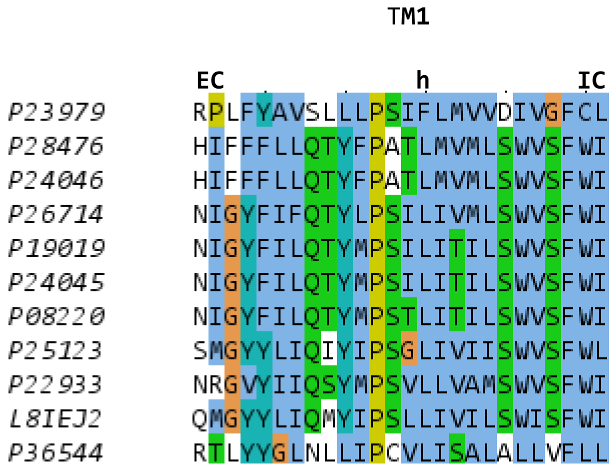
TM1
EC
h
IC
P23979 RPLFYAVSLLLPSIFLMVVDIVGFCL
P28476 HIFFFLLQTYFPATLMVMLSWVSFWI
P24046 HIFFFLLQTYFPATLMVMLSWVSFWI
P26714 NIGYFIFQTYLPSILIVMLSWVSFWI
P19019 NIGYFILQTYMPSILITILSWVSFWI
P24045 NIGYFILQTYMPSILITILSWVSFWI
P08220 NIGYFILQTYMPSTLITILSWVSFWI
P25123 SMGYYLIQIYIPSGLIVIISWVSFWL
P22933 NRGVYIIQSYMPSVLLVAMSWVSFWI
L8IEJ2 QMGYYLIQMYIPSLLIVILSWISFWI
P36544 RTLYYGLNLLIPCVLISALALLVFLL

### ACHA7 TM2

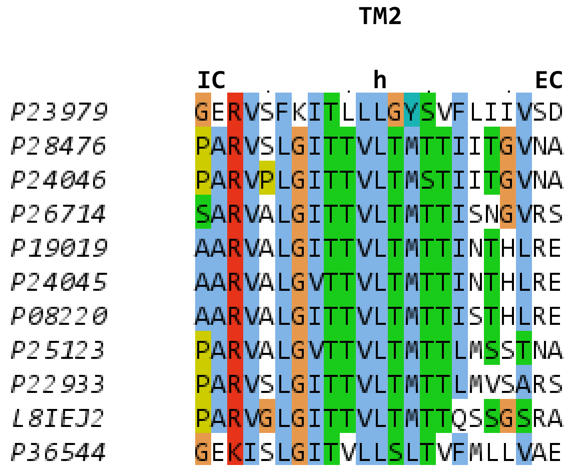
TM2
IC
h
EC
P23979 GERVSFKITLLLGYSVFLIIVSD
P28476 PARVSLGITTVLTMTTIITGVNA
P24046 PARVPLGITTVLTMSTIITGVNA
P26714 SARVALGITTVLTMTTISNGVRS
P19019 AARVALGITTVLTMTTINTHLRE
P24045 AARVALGVTTVLTMTTINTHLRE
P08220 AARVALGITTVLTMTTISTHLRE
P25123 PARVALGVTTVLTMTTLMSSTNA
P22933 PARVSLGITTVLTMTTLMVSARS
L8IEJ2 PARVGLGITTVLTMTTQSSGSRA
P36544 GEKISLGITVLLSLTVFMLLVAE

### ACHA7 TM3

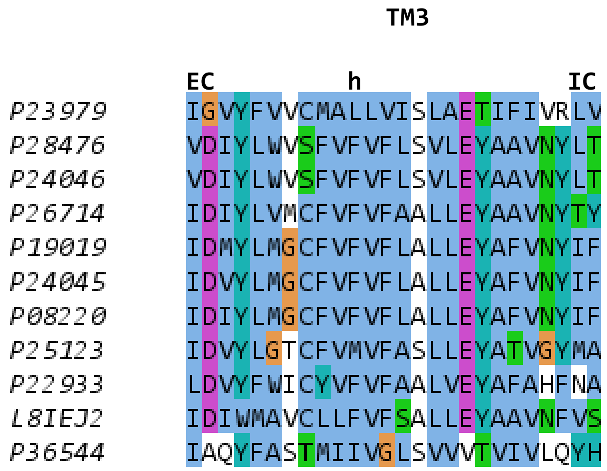
TM3
EC
h
IC
P23979 IGVYFVVCMALLVISLAETIFIVRLV
P28476 VDIYLWVSFVFVFLSVLEYAAVNYLT
P24046 VDIYLWVSFVFVFLSVLEYAAVNYLT
P26714 IDIYLVMCFVFVFAALLEYAAVNYTY
P19019 IDMYLMGCFVFVFLALLEYAFVNYIF
P24045 IDVYLMGCFVFVFLALLEYAFVNYIF
P08220 IDIYLMGCFVFVFLALLEYAFVNYIF
P25123 IDVYLGTCFVMVFASLLEYATVGYMA
P22933 LDVYFWICYVFVFAALVEYAFAHFNA
L8IEJ2 IDIWMAVCLLFVFSALLEYAAVNFVS
P36544 IAQYFASTMIIVGLSVVVTVIVLQYH

### ACHA7 TM4

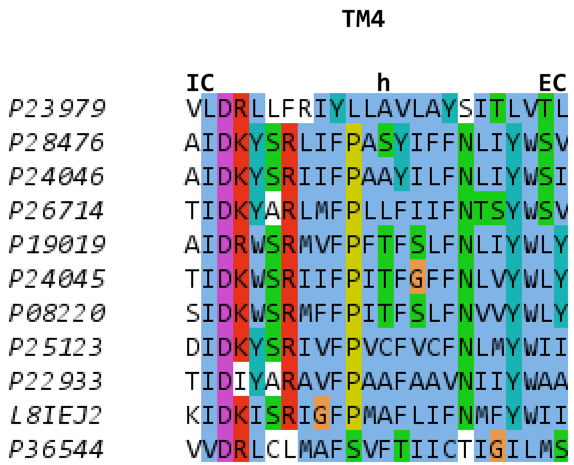
TM4
IC
h
EC
P23979 VLDRLLFRIYLLAVLAYSITLVTL
P28476 AIDKYSRLIFPASYIFFNLIYWSV
P24046 AIDKYSRIIFPAAYILFNLIYWSI
P26714 TIDKYARLMFPLLFIIFNTSYWSV
P19019 AIDRWSRMVFPFTFSLFNLIYWLY
P24045 TIDKWSRIIFPITFGFFNLVYWLY
P08220 SIDKWSRMFFPITFSLFNVVYWLY
P25123 DIDKYSRIVFPVCFVCFNLMYWII
P22933 TIDIYARAVFPAAFAAVNIIYWAA
L8IEJ2 KIDKISRIGFPMAFLIFNMFYWII
P36544 VVDRLCLMAFSVFTIICTIGILMS

### Aquaporin TM1

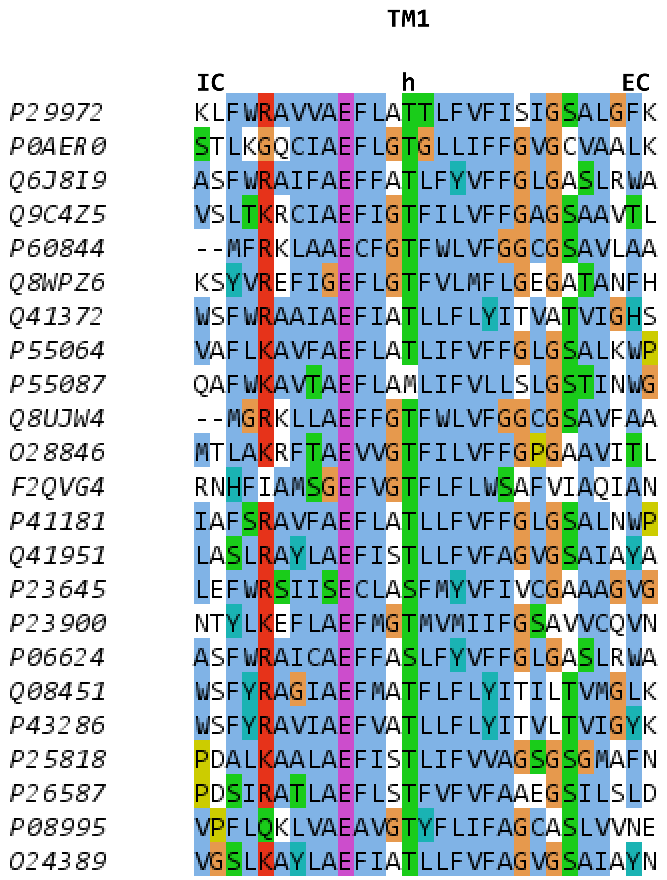

### Aquaporin TM2

```
                 TM2

         EC         h          IC
P29972   DNVKVSLAFGLSIATLAQSVGHIS
P0AER0   GQWEISVIWGLGVAMAIYLTAGVS
Q6J8I9   HVLQVALAFGLALATLVQAVGHIS
Q9C4Z5   DWVAIGLAFGFAIAASIYALGNIS
P60844   GFAGVALAFGLTVLTMAFAVGHIS
Q8WPZ6   DWYKLCLGWGLAVFFGILVSAKLS
Q41372   GLLGIAWAFGGMIFVLVYCTAGIS
P55064   TILQIALAFGLAIGTLAQALGPVS
P55087   DMVLISLCFGLSIATMVQCFGHIS
Q8UJW4   GFTGVALAFGLTVLTMAYAVGGIS
O28846   DWFAIGMAFALAIAAVIYSLGRIS
F2QVG4   QLIMISFGFGFGVMVGVFITYRVS
P41181   SVLQIAMAFGLGIGTLVQALGHIS
Q41951   GLVAIAVCHGFALFVAVAIGANIS
P23645   VLLATALASGLAMATLTQCFLHIS
P23900   TFDDVALGWAAAVVMGYFCAGGSA
P06624   HVLQVALAFGLALATLVQAVGHIS
Q08451   GIQGVAWAFGGMIFALVYCTAGIS
P43286   GILGIAWAFGGMIFILVYCTAGIS
P25818   GLVAAAVAHAFGLFVAVSVGANIS
P26587   GLILVALAHAFALFAAVSAAINVS
P08995   TFPGIAIVWGLVLTVLVYTVGHIS
O24389   GLVAIAVAHAFALFVGVSMAANIS
```

### Aquaporin TM2B

| | TM2B |
|---|---|
| | IC |
| P29972 | PAVTLGLLLSC |
| P0AER0 | PAVTIALWLFA |
| Q6J8I9 | PAVTFAFLVGS |
| Q9C4Z5 | PAVTIGLWSVK |
| P60844 | PAVTIGLWAGG |
| Q8WPZ6 | LAVSIGLSSIN |
| Q41372 | PAVTFGLFLAR |
| P55064 | PAITLALLVGN |
| P55087 | PAVTVAMVCTR |
| Q8UJW4 | PAVSVGLTVAG |
| O28846 | PAVTIALWSIG |
| F2QVG4 | PAVTLALVLAR |
| P41181 | PAVTVACLVGC |
| Q41951 | PAVTFGLAVGG |
| P23645 | PAVTLALCVVR |
| P23900 | PSITLANLVYR |
| P06624 | PAVTFAFLVGS |
| Q08451 | PAVTFGLFLAR |
| P43286 | PAVTFGLFLAR |
| P25818 | PAVTFGAFIGG |
| P26587 | PAVTFGALVGG |
| P08995 | PAVTIAFASTR |
| O24389 | PAVTLGLAVGG |

### Aquaporin TM3

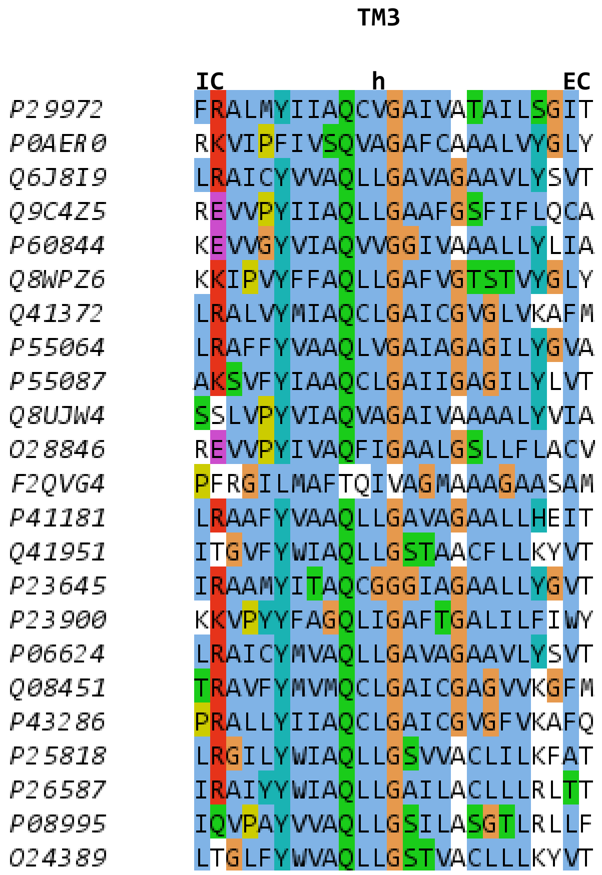

### Aquaporin TM4

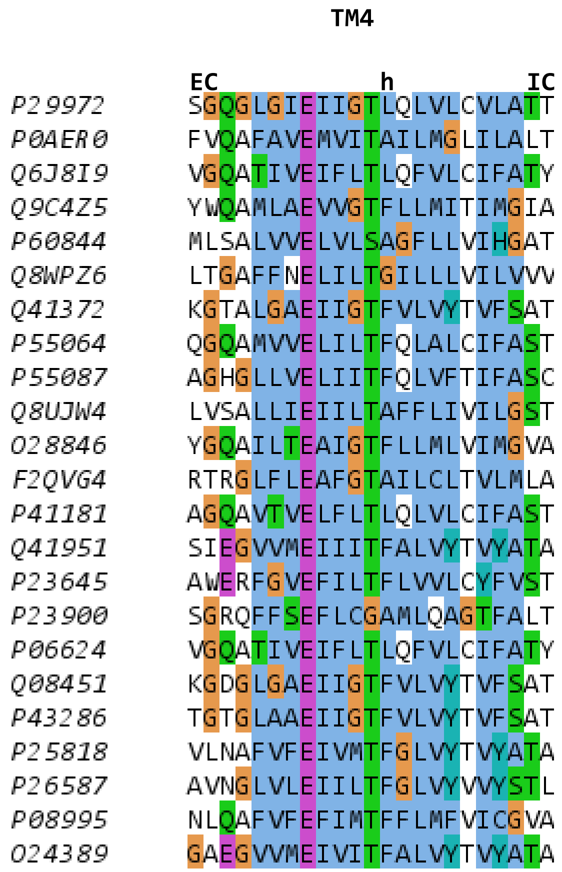

### Aquaporin TM5

```
                TM5

          IC      h          EC
P29972    SAPLAIGLSVALGHLLAIDYT
P0AER0    LAPLLIGLLIAVIGASMGPLT
Q6J8I9    SVALAVGFSLTLGHLFGMYYT
Q9C4Z5    FAGIIIGLTVAGIITTLGNIS
P60844    FAPIAIGLALTLIHLISIPVT
Q8WPZ6    KLSSVVGLIILCIGITFGGNT
Q41372    LAPLPIGFAVFMVHLATIPIT
P55064    SPALSIGLSVTLGHLVGIYFT
P55087    SIALAIGFSVAIGHLFAINYT
Q8UJW4    FAPIAIGLALTLIHLISIPVT
O28846    FAGLVIGLTVGGIITTIGNIT
F2QVG4    FAPFVIGIALLIAHLICIYYT
P41181    TPALSIGFSVALGHLLGIHYT
Q41951    IAPLAIGLIVGANILAAGPFS
P23645    NSAASIGCAYSACCFVSMPYL
P23900    VFPLMMFILIFIINASMAYQT
P06624    SVALAVGFSLTLGHLFGMYYT
Q08451    LAPLPIGFAVFLVHLATIPIT
P43286    LAPLPIGFAVFMVHLATIPIT
P25818    IAPIAIGFIVGANILAGGAFS
P26587    IAPLAIGLIVGANILVGGPFS
P08995    FAGIAIGSTLLLNVIIGGPVT
O24389    IAPIAIGFIVGANILAAGPFS
```

### Aquaporin TM5B

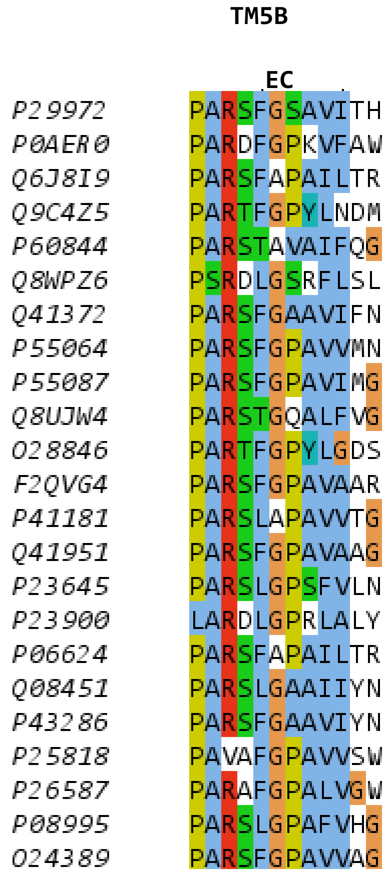

### Aquaporin TM6

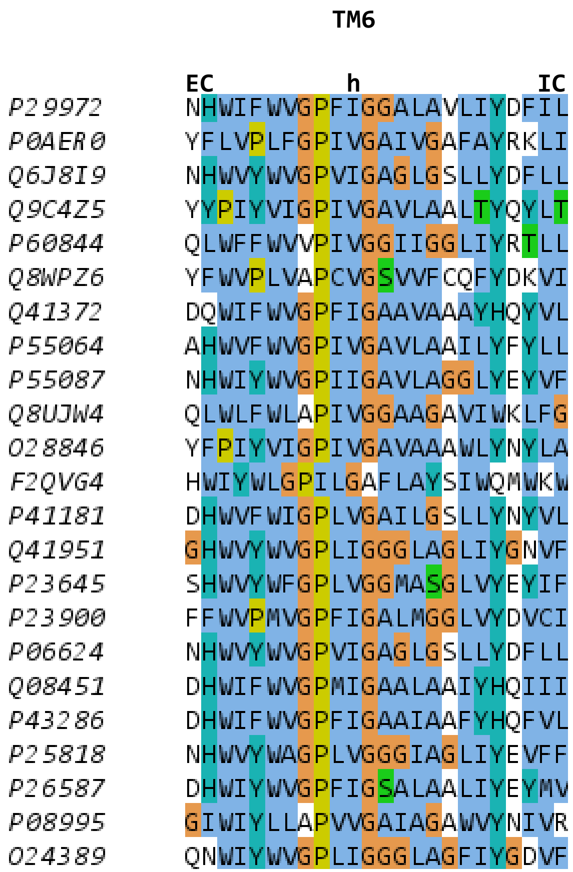

### Foca TM1

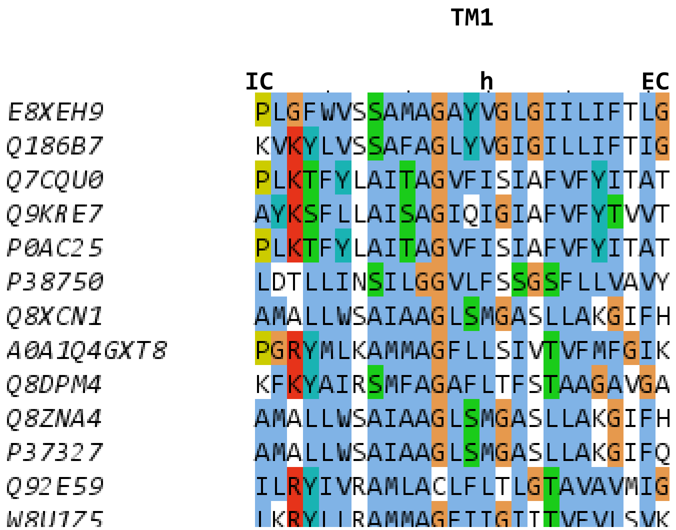
TM1
IC
h
EC
E8XEH9 PLGFWVSSAMAGAYVGLGIILIFTLG
Q186B7 KVKYLVSSAFAGLYVGIGILLIFTIG
Q7CQU0 PLKTFYLAITAGVFISIAFVFYITAT
Q9KRE7 AYKSFLLAISAGIQIGIAFVFYTVVT
P0AC25 PLKTFYLAITAGVFISIAFVFYITAT
P38750 LDTLLINSILGGVLFSSGSFLLVAVY
Q8XCN1 AMALLWSAIAAGLSMGASLLAKGIFH
A0A1Q4GXT8 PGRYMLKAMMAGFLLSIVTVFMFGIK
Q8DPM4 KFKYAIRSMFAGAFLTFSTAAGAVGA
Q8ZNA4 AMALLWSAIAAGLSMGASLLAKGIFH
P37327 AMALLWSAIAAGLSMGASLLAKGIFQ
Q92E59 ILRYIVRAMLACLFLTLGTAVAVMIG

### Foca TM2

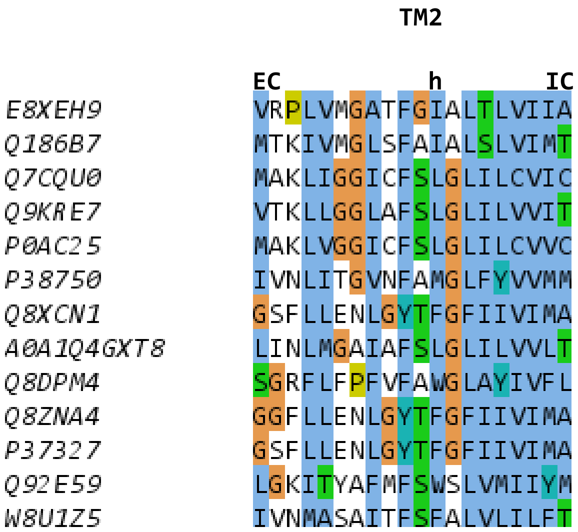
TM2
EC
h
IC
E8XEH9 VRPLVMGATFGIALTLVIIA
Q186B7 MTKIVMGLSFAIALSLVIMT
Q7CQU0 MAKLIGGICFSLGLILCVIC
Q9KRE7 VTKLLGGLAFSLGLILVVIT
P0AC25 MAKLVGGICFSLGLILCVVC
P38750 IVNLITGVNFAMGLFYVVMM
Q8XCN1 GSFLLENLGYTFGFIIVIMA
A0A1Q4GXT8 LINLMGAIAFSLGLILVVLT
Q8DPM4 SGRFLFPFVFAWGLAYIVFL
Q8ZNA4 GGFLLENLGYTFGFIIVIMA
P37327 GSFLLENLGYTFGFIIVIMA
Q92E59 LGKITYAFMFSWSLVMIIYM
W8U1Z5 IVNMASAITFSFALVLILFT

### Foca TM2B

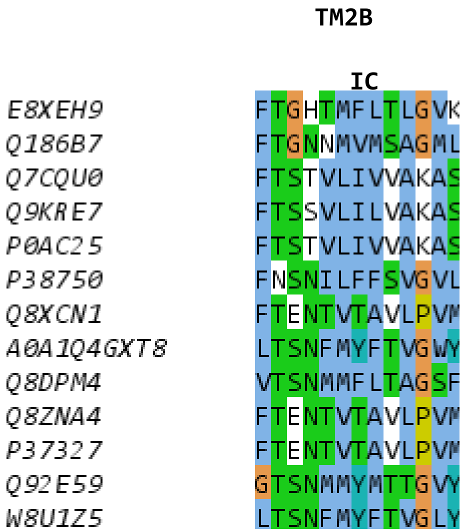

TM2B
IC
E8XEH9 FTGHTMFLTLGVK
Q186B7 FTGNNMVMSAGML
Q7CQU0 FTSTVLIVVAKAS
Q9KRE7 FTSSVLILVAKAS
P0AC25 FTSTVLIVVAKAS
P38750 FNSNILFFSVGVL
Q8XCN1 FTENTVTAVLPVM
A0A1Q4GXT8 LTSNFMYFTVGWY
Q8DPM4 VTSNMMFLTAGSF
Q8ZNA4 FTENTVTAVLPVM
P37327 FTENTVTAVLPVM
Q92E59 GTSNMMYMTTGVY
W8U1Z5 LTSNFMYFTVGLY

### Foca TM3

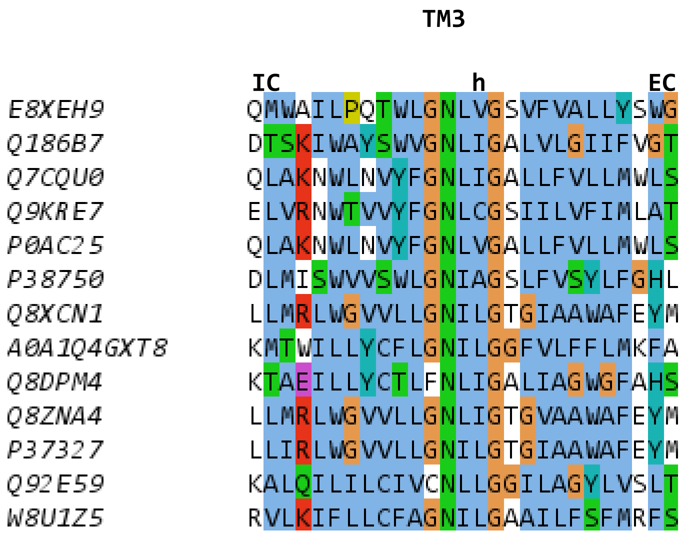
TM3
IC
h
EC
E8XEH9 QMWAILPQTWLGNLVGSVFVALLYSWG
Q186B7 DTSKIWAYSWVGNLIGALVLGIIFVGT
Q7CQU0 QLAKNWLNVYFGNLIGALLFVLLMWLS
Q9KRE7 ELVRNWTVVYFGNLCGSIILVFIMLAT
P0AC25 QLAKNWLNVYFGNLVGALLFVLLMWLS
P38750 DLMISWVVSWLGNIAGSLFVSYLFGHL
Q8XCN1 LLMRLWGVVLLGNILGTGIAAWAFEYM
A0A1Q4GXT8 KMTWILLYCFLGNILGGFVLFFLMKFA
Q8DPM4 KTAEILLYCTLFNLIGALIAGWGFAHS
Q8ZNA4 LLMRLWGVVLLGNLIGTGVAAWAFEYM
P37327 LLIRLWGVVLLGNILGTGIAAWAFEYM
Q92E59 KALQILILCIVCNLLGGILAGYLVSLT
W8U1Z5 RVLKIFLLCFAGNILGAAILFSFMRFS

### Foca TM4

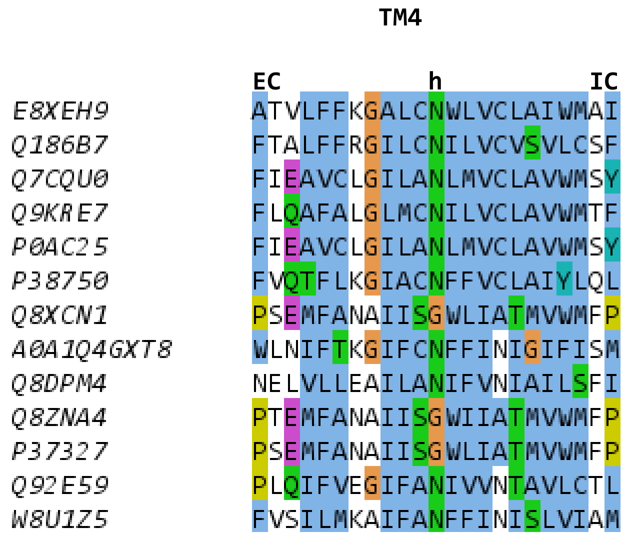
TM4
EC
h
IC
E8XEH9 ATVLFFKGALCNWLVCLAIWMAI
Q186B7 FTALFFRGILCNILVCVSVLCSF
Q7CQU0 FIEAVCLGILANLMVCLAVWMSY
Q9KRE7 FLQAFALGLMCNILVCLAVWMTF
P0AC25 FIEAVCLGILANLMVCLAVWMSY
P38750 FVQTFLKGIACNFFVCLAIYLQL
Q8XCN1 PSEMFANAIISGWLIATMVWMFP
A0A1Q4GXT8 WLNIFTKGIFCNFFINIGIFISM
Q8DPM4 NELVLLEAILANIFVNIAILSFI
Q8ZNA4 PTEMFANAIISGWIIATMVWMFP
P37327 PSEMFANAIISGWLIATMVWMFP
Q92E59 PLQIFVEGIFANIVVNTAVLCTL
W8U1Z5 FVSILMKAIFANFFINISLVIAM

### Foca TM5

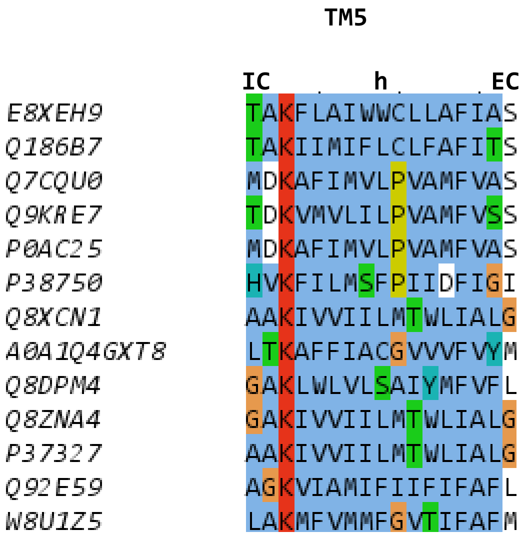
TM5
IC
h
EC
E8XEH9 TAKFLAIWWCLLAFIAS
Q186B7 TAKIIMIFLCLFAFITS
Q7CQU0 MDKAFIMVLPVAMFVAS
Q9KRE7 TDKVMVLILPVAMFVSS
P0AC25 MDKAFIMVLPVAMFVAS
P38750 HVKFILMSFPIIDFIGI
Q8XCN1 AAKIVVIILMTWLIALG
A0A1Q4GXT8 LTKAFFIACGVVVFVYM
Q8DPM4 GAKLWLVLSAIYMFVFL
Q8ZNA4 GAKIVVIILMTWLIALG
P37327 AAKIVVIILMTWLIALG
Q92E59 AGKVIAMIFIIFIFAFL
W8U1Z5 LAKMFVMMFGVTIFAFM

### Foca TM5B

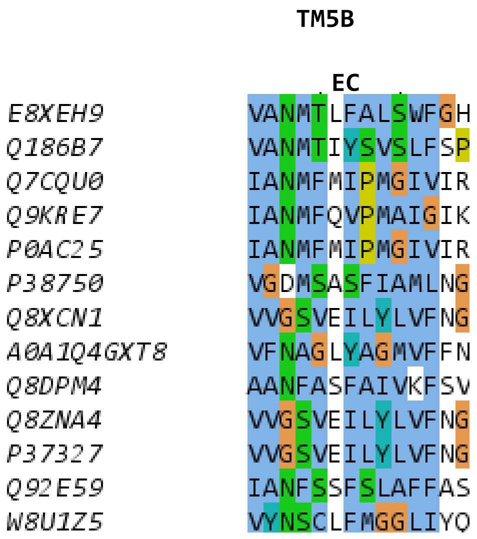
TM5B
EC
E8XEH9 VANMTLFALSWFGH
Q186B7 VANMTIYSVSLFSP
Q7CQU0 IANMFMIPMGIVIR
Q9KRE7 IANMFQVPMAIGIK
P0AC25 IANMFMIPMGIVIR
P38750 VGDMSASFIAMLNG
Q8XCN1 VVGSVEILYLVFNG
A0A1Q4GXT8 VFNAGLYAGMVFFN
Q8DPM4 AANFASFAIVKFSV
Q8ZNA4 VVGSVEILYLVFNG
P37327 VVGSVEILYLVFNG
Q92E59 IANFSSFSLAFFAS
W8U1Z5 VYNSCLFMGGLIYQ

### Foca TM6

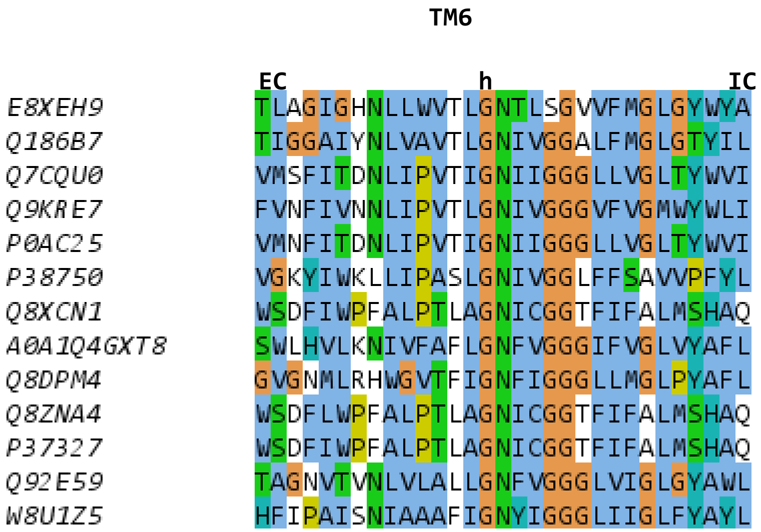

TM6
EC
h
IC
E8XEH9 TLAGIGHNLLWVTLGNTLSGVVFMGLGYWYA
Q186B7 TIGGAIYNLVAVTLGNIVGGALFMGLGTYIL
Q7CQU0 VMSFITDNLIPVTIGNIIGGGLLVGLTYWVI
Q9KRE7 FVNFIVNNLIPVTLGNIVGGGVFVGMWYWLI
P0AC25 VMNFITDNLIPVTIGNIIGGGLLVGLTYWVI
P38750 VGKYIWKLLIPASLGNIVGGLFFSAVVPFYL
Q8XCN1 WSDFIWPFALPTLAGNICGGTFIFALMSHAQ
A0A1Q4GXT8 SWLHVLKNIVFAFLGNFVGGGIFVGLVYAFL
Q8DPM4 GVGNMLRHWGVTFIGNFIGGGLLMGLPYAFL
Q8ZNA4 WSDFLWPFALPTLAGNICGGTFIFALMSHAQ
P37327 WSDFIWPFALPTLAGNICGGTFIFALMSHAQ
Q92E59 TAGNVTVNLVLALLGNFVGGGLVIGLGYAWL
W8U1Z5 HFIPAISNIAAAFIGNYIGGGLIIGLFYAYL

### GPCR TM1

```
                 TM1
        EC             h          IC
O14842  LPPQLSFGLYVAAFALGFPLNVLAIRGATA
O43613  QYEWVLIAAYVAVFVVALVGNTLVCLAVWR
P07550  VWVVGMGIVMSLIVLAIVFGNVLVITAIAK
P08100  WQFSMLAAYMFLLIVLGFPINFLTLYVTVQ
P08172  FEVVFIVLVAGSLSLVTIIGNILVMVSIKV
P08173  VEMVFIATVTGSLSLVTVVGNILVMLSIKV
P11229  WQVAFIGITTGLLSLATVTGNLLVLISFKV
P21453  NSIKLTSVVFILICCFIILENIFVLLTIWK
P21554  SQQLAIAVLSLTLGTFTVLENLLVLCVILH
P24530  TFKYINTVVSCLVFVLGIIGNSTLLRIIYK
P25024  LNKYVVIIAYALVFLLSLLGNSLVMLVILY
P25116  WLTLFVPSVYTGVFVVSLPLNIMAIVVFIL
P28222  PWKVLLVMLLALITLATTLSNAFVIATVYR
P29274  MGSSVYITVELAIAVLAILGNVLVCWAVWL
P30542  AFQAAYIGIEVLIALVSVPGNVLVIWAVKV
P35367  PQLMPLVVVLSTICLVTVGLNLLVLYAVRS
P35462  RPHAYYALSYCALILAIVFGNGLVCMAVLK
P41143  ALAIAITALYSAVCAVGLLGNVLVMFGIVR
P41145  AIPVIITAVYSVVFVVGLVGNSLVMFVIIR
P41146  GLKVTIVGLYLAVCVGGLLGNCLVMYVILR
P41597  IGAQLLPPLYSLVFIFGFVGNMLVVLILIN
P47900  FQFYYLPAVYILVFIIGFLGNSVAIWMFVF
P50052  KHLDAIPILYYIIFVIGFLVNIVVVTLFCC
P51681  IAARLLPPLYSLVFIFGFVGNMLVILILIN
P51686  FASHFLPPLYWLVFIVGALGNSLVILVYWY
P55085  LTTVFLPIVYTIVFVVGLPSNGMALWVFLF
P61073  FNKIFLPTIYSIIFLTGIVGNGLVILVMGY
Q92633  TVSKLVMGLGITVCIFIMLANLLVMVAIYV
Q9H244  ITQVLFPLLYTVLFFVGLITNGLAMRIFFQ
P20789  YSKVLVTAIYLALFVVGTVGNSVTAFTLAR
P34998  VHYHVAVIINYLGHCISLVALLVAFVLFLR
P43220  EQLLFLYIIYTVGYALSFSALVIASAILLG
P47871  KMYSSFQVMYTVGYSLSLGALLLALAILGG
P41594  RWGDPEPIAAVVFACLGLLATLFVTVVFII
Q13255  EWSNIESIIAIAFSCLGILVTLFVTLIFVL
Q99835  AEHQDMHSYIAAFGAVTGLCTLFTLATFVA
```

### GPCR TM2

```
                    TM2

           IC              h              EC
O14842     PSLVYALNLGCSDLLLTVSLPLKAVEALA
O43613     VTNYFIVNLSLADVLVTAICLPASLLVDI
P07550     VTNYFITSLACADLVMGLAVVPFGAAHIL
P08100     PLNYILLNLAVADLFMVLGGFTSTLYTSL
P08172     VNNYFLFSLACADLIIGVFSMNLYTLYTV
P08173     VNNYFLFSLACADLIIGAFSMNLYTVYII
P11229     VNNYFLLSLACADLIIGTFSMNLYTTYLL
P21453     PMYYFIGNLALSDLLAGVAYTANLLLSGA
P21554     PSYHFIGSLAVADLLGSVIFVYSFIDFHV
P24530     GPNILIASLALGDLLHIVIDIPINVYKLL
P25024     VTDVYLLNLALADLLFALTLPIWAASKVN
P25116     PAVVYMLHLATADVLFVSVLPFKISYYFS
P28222     PANYLIASLAVTDLLVSILVMPISTMYTV
P29274     VTNYFVVSLAAADIAVGVLAIPFAITIST
P30542     ATFCFIVSLAVADVAVGALVIPLAILINI
P35367     VGNLYIVSLSVADLIVGAVVMPMNILYLL
P35462     TTNYLVVSLAVADLLVATLVMPWVVYLEV
P41143     ATNIYIFNLALADALATSTLPFQSAKYLM
P41145     ATNIYIFNLALADALVTTTMPFQSTVYLM
P41146     ATNIYIFNLALADTLVLLTLPFQGTDILL
P41597     LTDIYLLNLAISDLLFLITLPLWAHSAAN
P47900     GISVYMFNLALADFLYVLTLPALIFYYFN
P50052     VSSIYIFNLAVADLLLLATLPLWATYYSY
P51681     MTDIYLLNLAISDLFFLLTVPFWAHYAAA
P51686     MTDMFLLNLAIADLLFLVTLPFWAIAAAD
P55085     PAVIYMANLALADLLSVIWFPLKIAYHIH
P61073     MTDKYRLHLSVADLLFVITLPFWAVDAVA
Q92633     PIYYLMANLAAADFFAGLAYFYLMFNTGP
Q9H244     NFIIFLKNTVISDLLMILTFPFKILSDAK
P20789     TVHYHLGSLALSDLLILLLAMPVELYNFI
P34998     LRNIIHWNLISAFILRNATWFVVQLTMSP
P43220     TRNYIHLNLFASFILRALSVFIKDAALKW
P47871     TRNAIHANLFASFVLKASSVLVIDGLLRT
P41594     SSRELCYIILAGICLGYLCTFCLIAKPK-
Q13255     SSRELCYIILAGIFLGYVCPFTLIAKPT-
Q99835     YPAVILFYVNACFFVGSIGWLAQFMDGAR
```

### GPCR TM3

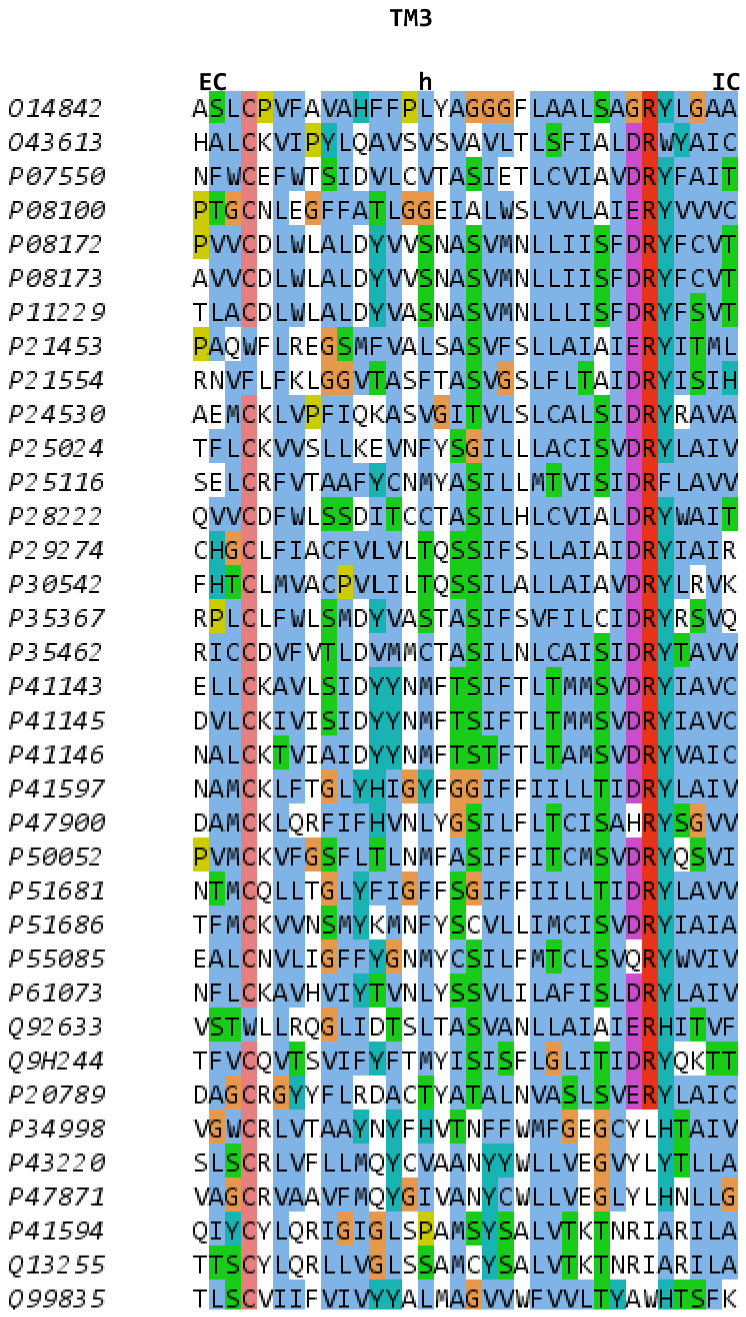
TM3
EC
h
IC
O14842 ASLCPVFAVAHFFPLYAGGGFLAALSAGRYLGAA
O43613 HALCKVIPYLQAVSVSVAVLTLSFIALDRWYAIC
P07550 NFWCEFWTSIDVLCVTASIETLCVIAVDRYFAIT
P08100 PTGCNLEGFFATLGGEIALWSLVVLAIERYVVVC
P08172 PVVCDLWLALDYVVSNASVMNLLIISFDRYFCVT
P08173 AVVCDLWLALDYVVSNASVMNLLIISFDRYFCVT
P11229 TLACDLWLALDYVASNASVMNLLLISFDRYFSVT
P21453 PAQWFLREGSMFVALSASVFSLLAIAIERYITML
P21554 RNVFLFKLGGVTASFTASVGSLFLTAIDRYISIH
P24530 AEMCKLVPFIQKASVGITVLSLCALSIDRYRAVA
P25024 TFLCKVVSLLKEVNFYSGILLLACISVDRYLAIV
P25116 SELCRFVTAAFYCNMYASILLMTVISIDRFLAVV
P28222 QVVCDFWLSSDITCCTASILHLCVIALDRYWAIT
P29274 CHGCLFIACFVLVLTQSSIFSLLAIAIDRYIAIR
P30542 FHTCLMVACPVLILTQSSILALLAIAVDRYLRVK
P35367 RPLCLFWLSMDYVASTASIFSVFILCIDRYRSVQ
P35462 RICCDVFVTLDVMMCTASILNLCAISIDRYTAVV
P41143 ELLCKAVLSIDYYNMFTSIFTLTMMSVDRYIAVC
P41145 DVLCKIVISIDYYNMFTSIFTLTMMSVDRYIAVC
P41146 NALCKTVIAIDYYNMFTSTFTLTAMSVDRYVAIC
P41597 NAMCKLFTGLYHIGYFGGIFFIILLTIDRYLAIV
P47900 DAMCKLQRFIFHVNLYGSILFLTCISAHRYSGVV
P50052 PVMCKVFGSFLTLNMFASIFFITCMSVDRYQSVI
P51681 NTMCQLLTGLYFIGFFSGIFFIILLTIDRYLAVV
P51686 TFMCKVVNSMYKMNFYSCVLLIMCISVDRYIAIA
P55085 EALCNVLIGFFYGNMYCSILFMTCLSVQRYWVIV
P61073 NFLCKAVHVIYTVNLYSSVLILAFISLDRYLAIV
Q92633 VSTWLLRQGLIDTSLTASVANLLAIAIERHITVF
Q9H244 TFVCQVTSVIFYFTMYISISFLGLITIDRYQKTT
P20789 DAGCRGYYFLRDACTYATALNVASLSVERYLAIC
P34998 VGWCRLVTAAYNYFHVTNFFWMFGEGCYLHTAIV
P43220 SLSCRLVFLLMQYCVAANYYWLLVEGVYLYTLLA
P47871 VAGCRVAAVFMQYGIVANYCWLLVEGLYLHNLLG
P41594 QIYCYLQRIGIGLSPAMSYSALVTKTNRIARILA
Q13255 TTSCYLQRLLVGLSSAMCYSALVTKTNRIARILA
Q99835 TLSCVIIFVIVYYALMAGVVWFVVLTYAWHTSFK

### GPCR TM4

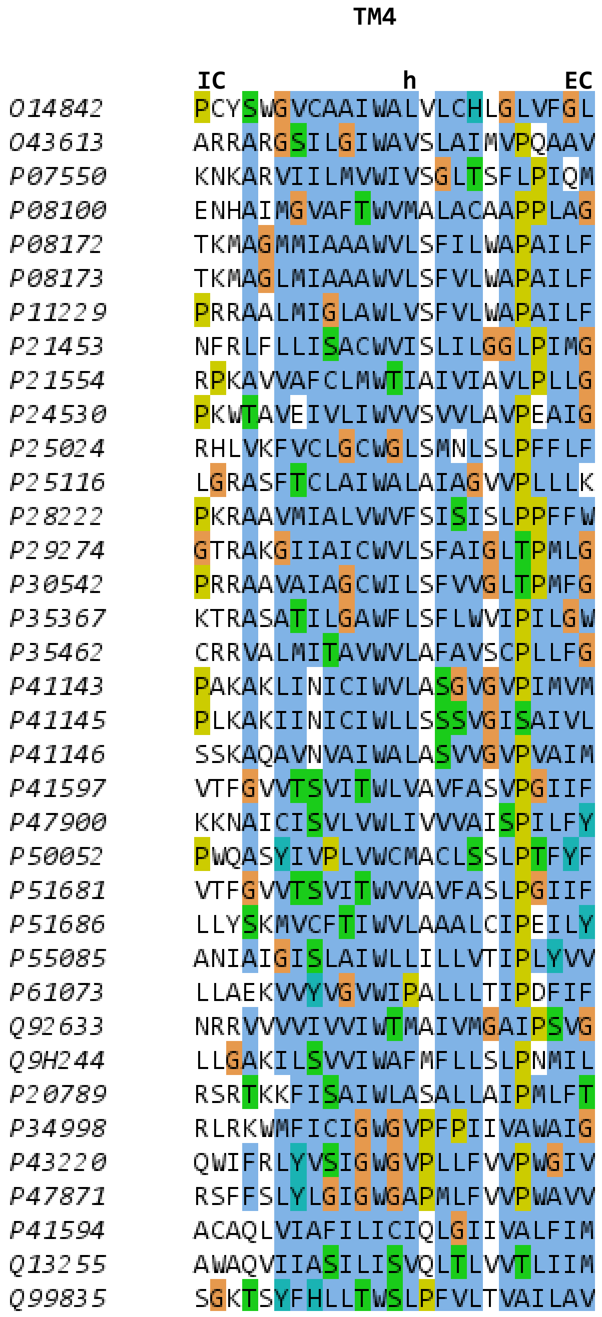
TM4
IC
h
EC
O14842 PCYSWGVCAAIWALVLCHLGLVFGL
O43613 ARRARGSILGIWAVSLAIMVPQAAV
P07550 KNKARVIILMVWIVSGLTSFLPIQM
P08100 ENHAIMGVAFTWVMALACAAPPLAG
P08172 TKMAGMMIAAAWVLSFILWAPAILF
P08173 TKMAGLMIAAAWVLSFVLWAPAILF
P11229 PRRAALMIGLAWLVSFVLWAPAILF
P21453 NFRLFLLISACWVISLILGGLPIMG
P21554 RPKAVVAFCLMWTIAIVIAVLPLLG
P24530 PKWTAVEIVLIWVVSVVLAVPEAIG
P25024 RHLVKFVCLGCWGLSMNLSLPFFLF
P25116 LGRASFTCLAIWALAIAGVVPLLLK
P28222 PKRAAVMIALVWVFSISISLPPFFW
P29274 GTRAKGIIAICWVLSFAIGLTPMLG
P30542 PRRAAVAIAGCWILSFVVGLTPMFG
P35367 KTRASATILGAWFLSFLWVIPILGW
P35462 CRRVALMITAVWVLAFAVSCPLLFG
P41143 PAKAKLINICIWVLASGVGVPIMVM
P41145 PLKAKIINICIWLLSSSVGISAIVL
P41146 SSKAQAVNVAIWALASVVGVPVAIM
P41597 VTFGVVTSVITWLVAVFASVPGIIF
P47900 KKNAICISVLVWLIVVVAISPILFY
P50052 PWQASYIVPLVWCMACLSSLPTFYF
P51681 VTFGVVTSVITWVVAVFASLPGIIF
P51686 LLYSKMVCFTIWVLAAALCIPEILY
P55085 ANIAIGISLAIWLLILLVTIPLYVV
P61073 LLAEKVVYVGVWIPALLLTIPDFIF
Q92633 NRRVVVVIVVIWTMAIVMGAIPSVG
Q9H244 LLGAKILSVVIWAFMFLLSLPNMIL
P20789 RSRTKKFISAIWLASALLAIPMLFT
P34998 RLRKWMFICIGWGVPFPIIVAWAIG
P43220 QWIFRLYVSIGWGVPLLFVVPWGIV
P47871 RSFFSLYLGIGWGAPMLFVVPWAVV
P41594 ACAQLVIAFILICIQLGIIVALFIM
Q13255 AWAQVIIASILISVQLTLVVTLIIM
Q99835 SGKTSYFHLLTWSLPFVLTVAILAV

### GPCR TM5

```
                 TM5
        EC                 h               IC
O14842  SAGPARFSLSLLLFFLPLAITAFCYVGCLRALAR
O43613  YPKIYHSCFFIVTYLAPLGLMAMAYFQIFRKLWG
P07550  TNQAYAIASSIVSFYVPLVIMVFVYSRVFQEAKR
P08100  NNESFVIYMFVVHFTIPMIIIFFCYGQLVFTVKE
P08172  SNAAVTFGTAIAAFYLPVIIMTVLYWHISRASKS
P08173  SNPAVTFGTAIAAFYLPVVIMTVLYIHISLASRS
P11229  SQPIITFGTAMAAFYLPVTVMCTLYWRIYRETEN
P21453  LYHKHYILFCTTVFTLLLLSIVILYCRIYSLVRT
P21554  HIDETYLMFWIGVTSVLLLFIVYAYMYILWKAHS
P24530  YKTAKDWWLFSFYFCLPLAITAFFYTLMTCEMLR
P25024  WRMVLRILPHTFGFIVPLFVMLFCYGFTLRTLFK
P25116  YYAYYFSAFSAVFFFVPLIISTVCYVSIIRCLSS
P28222  DHILYTVYSTVGAFYFPTLLLIALYGRIYVEARS
P29274  PMNYMVYFNFFACVLVPLLLMLGVYLRIFLAARR
P30542  SMEYMVYFNFFVWVLPPLLLMVLIYLEVFYLIRK
P35367  DVTWFKVMTAIINFYLPTLLMLWFYAKIYKAVRQ
P35462  SNPDFVIYSSVVSFYLPFGVTVLVYARIYVVLKQ
P41143  WDTVTKICVFLFAFVVPILIITVCYGLMLLRLRS
P41145  WDLFMKICVFIFAFVIPVLIIIVCYTLMILRLKS
P41146  WGPVFAICIFLFSFIVPVLVISVCYSLMIRRLRG
P41597  WNNFHTIMRNILGLVLPLLIMVICYSGILKTLLR
P47900  SYFIYSMCTTVAMFCVPLVLILGCYGLIVRALIY
P50052  WSAGIALMKNILGFIIPLIFIATCYFGIRKHLLK
P51681  WKNFQTLKIVILGLVLPLLVMVICYSGILKTLLR
P51686  LKSAVLTLKVILGFFLPFVVMACCYTIIIHTLIQ
P55085  DMFNYFLSLAIGVFLFPAFLTASAYVLMIRMLRS
P61073  WVVVFQFQHIMVGLILPGIVILSCYCIIISKLSH
Q92633  LYSDSYLVFWAIFNLVTFVVMVVLYAHIFGYVRQ
Q9H244  VWHEIVNYICQVIFWINFLIVIVCYTLITKELYR
P20789  TVKVVIQVNTFMSFLFPMLVISILNTVIANKLTV
P34998  RPGVYTDYIYQGPMILVLLINFIFLFNIVRILMT
P43220  NSNMNYWLIIRLPILFAIGVNFLIFVRVICIVVS
P47871  NDNMGFWWILRFPVFLAILINFFIFVRIVQLLVA
P41594  NTTNLGVVTPLGYNGLLILSCTFYAFKTRNVPA-
Q13255  NTSNLGVVAPLGYNGLLIMSCTYYAFKTRNVPA-
Q99835  KNYRYRAGFVLAPIGLVLIVGGYFLIRGVMTLFS
```

### GPCR TM6

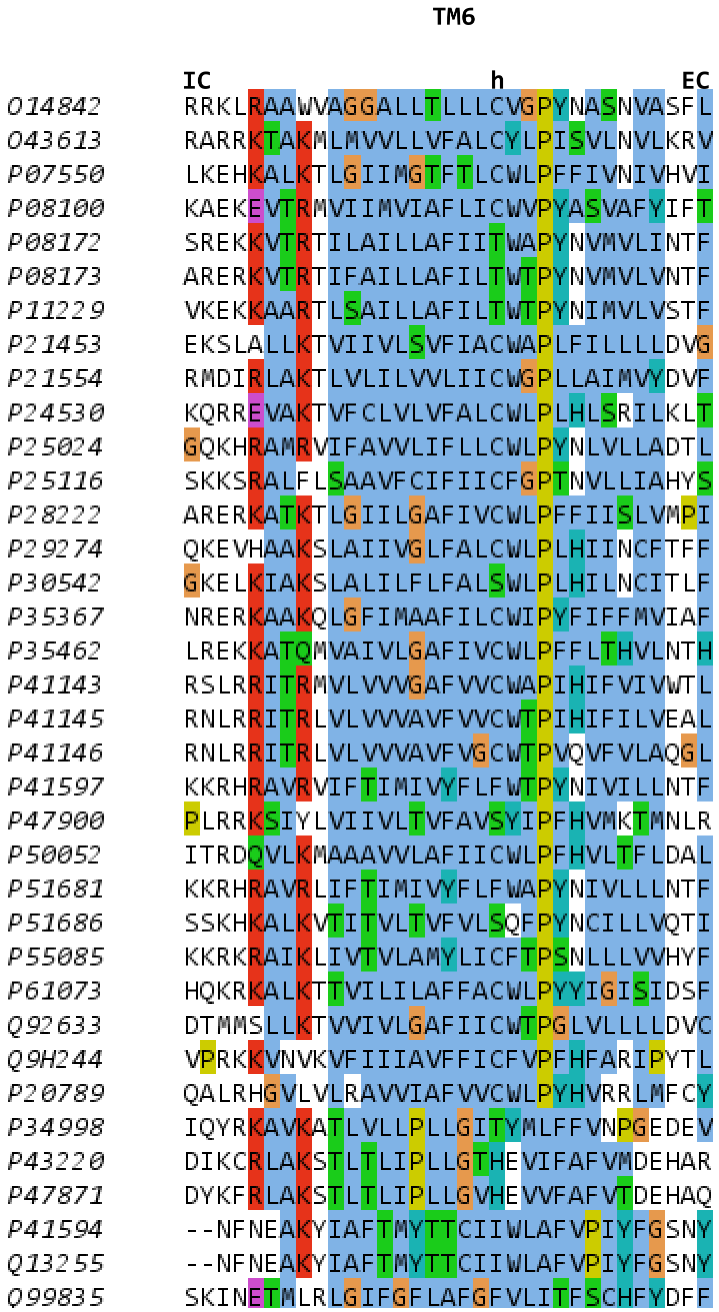

TM6
IC
h
EC
O14842 RRKLRAAWVAGGALLTLLLCVGPYNASNVASFL
O43613 RARRKTAKMLMVVLLVFALCYLPISVLNVLKRV
P07550 LKEHKALKTLGIIMGTFTLCWLPFFIVNIVHVI
P08100 KAEKEVTRMVIIMVIAFLICWVPYASVAFYIFT
P08172 SREKKVTRTILAILLAFIITWAPYNVMVLINTF
P08173 ARERKVTRTIFAILLAFILTWTPYNVMVLVNTF
P11229 VKEKKAARTLSAILLAFILTWTPYNIMVLVSTF
P21453 EKSLALLKTVIIVLSVFIACWAPLFILLLLDVG
P21554 RMDIRLAKTLVLILVVLIICWGPLLAIMVYDVF
P24530 KQRREVAKTVFCLVLVFALCWLPLHLSRILKLT
P25024 GQKHRAMRVIFAVVLIFLLCWLPYNLVLLADTL
P25116 SKKSRALFLSAAVFCIFIICFGPTNVLLIAHYS
P28222 ARERKATKTLGIILGAFIVCWLPFFIISLVMPI
P29274 QKEVHAAKSLAIIVGLFALCWLPLHIINCFTFF
P30542 GKELKIAKSLALILFLFALSWLPLHILNCITLF
P35367 NRERKAAKQLGFIMAAFILCWIPYFIFFMVIAF
P35462 LREKKATQMVAIVLGAFIVCWLPFFLTHVLNTH
P41143 RSLRRITRMVLVVVGAFVVCWAPIHIFVIVWTL
P41145 RNLRRITRLVLVVVAVFVVCWTPIHIFILVEAL
P41146 RNLRRITRLVLVVVAVFVGCWTPVQVFVLAQGL
P41597 KKRHRAVRVIFTIMIVYFLFWTPYNIVILLNTF
P47900 PLRRKSIYLVIIVLTVFAVSYIPFHVMKTMNLR
P50052 ITRDQVLKMAAAVVLAFIICWLPFHVLTFLDAL
P51681 KKRHRAVRLIFTIMIVYFLFWAPYNIVLLLNTF
P51686 SSKHKALKVTITVLTVFVLSQFPYNCILLVQTI
P55085 KKRKRAIKLIVTVLAMYLICFTPSNLLLVVHYF
P61073 HQKRKALKTTVILILAFFACWLPYYIGISIDSF
Q92633 DTMMSLLKTVVIVLGAFIICWTPGLVLLLLDVC
Q9H244 VPRKKVNVKVFIIIAVFFICFVPFHFARIPYTL
P20789 QALRHGVLVLRAVVIAFVVCWLPYHVRRLMFCY
P34998 IQYRKAVKATLVLLPLLGITYMLFFVNPGEDEV
P43220 DIKCRLAKSTLTLIPLLGTHEVIFAFVMDEHAR
P47871 DYKFRLAKSTLTLIPLLGVHEVVFAFVTDEHAQ
P41594 --NFNEAKYIAFTMYTTCIIWLAFVPIYFGSNY
Q13255 --NFNEAKYIAFTMYTTCIIWLAFVPIYFGSNY
Q99835 SKINETMLRLGIFGFLAFGFVLITFSCHFYDFF

### GPCR TM7

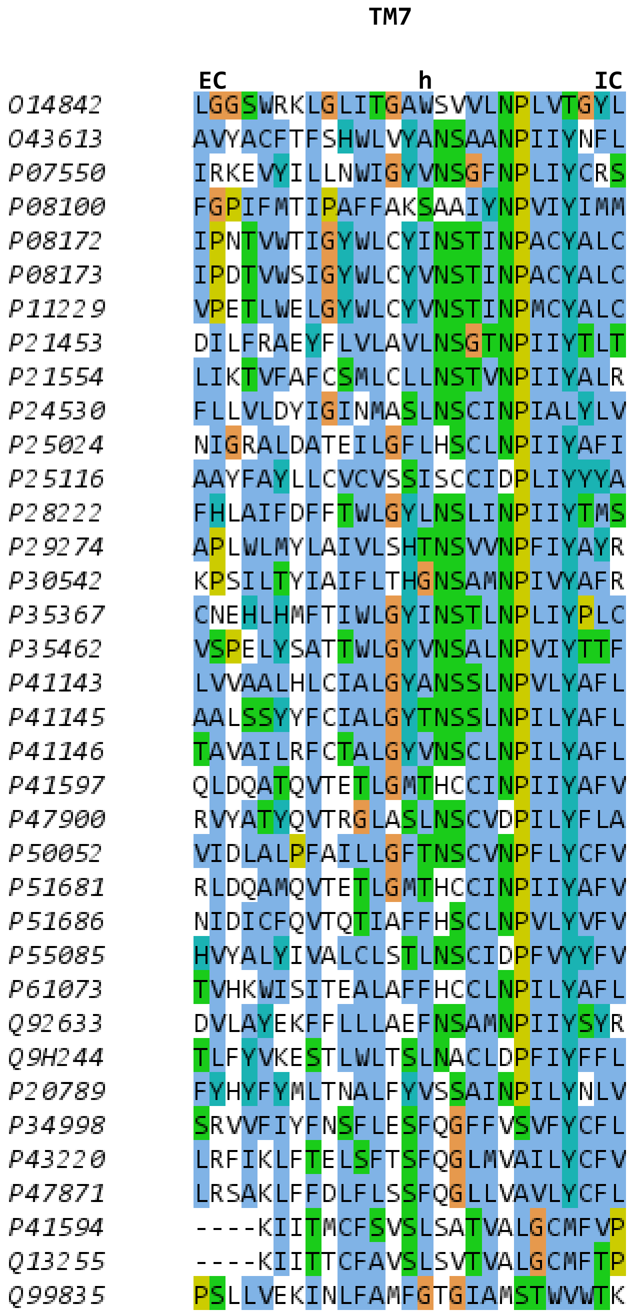
TM7
EC
h
IC
O14842 LGGSWRKLGLITGAWSVVLNPLVTGYL
O43613 AVYACFTFSHWLVYANSAANPIIYNFL
P07550 IRKEVYILLNWIGYVNSGFNPLIYCRS
P08100 FGPIFMTIPAFFAKSAAIYNPVIYIMM
P08172 IPNTVWTIGYWLCYINSTINPACYALC
P08173 IPDTVWSIGYWLCYVNSTINPACYALC
P11229 VPETLWELGYWLCYVNSTINPMCYALC
P21453 DILFRAEYFLVLAVLNSGTNPIIYTLT
P21554 LIKTVFAFCSMLCLLNSTVNPIIYALR
P24530 FLLVLDYIGINMASLNSCINPIALYLV
P25024 NIGRALDATEILGFLHSCLNPIIYAFI
P25116 AAYFAYLLCVCVSSISCCIDPLIYYYA
P28222 FHLAIFDFFTWLGYLNSLINPIIYTMS
P29274 APLWLMYLAIVLSHTNSVVNPFIYAYR
P30542 KPSILTYIAIFLTHGNSAMNPIVYAFR
P35367 CNEHLHMFTIWLGYINSTLNPLIYPLC
P35462 VSPELYSATTWLGYVNSALNPVIYTTF
P41143 LVVAALHLCIALGYANSSLNPVLYAFL
P41145 AALSSYYFCIALGYTNSSLNPILYAFL
P41146 TAVAILRFCTALGYVNSCLNPILYAFL
P41597 QLDQATQVTETLGMTHCCINPIIYAFV
P47900 RVYATYQVTRGLASLNSCVDPILYFLA
P50052 VIDLALPFAILLGFTNSCVNPFLYCFV
P51681 RLDQAMQVTETLGMTHCCINPIIYAFV
P51686 NIDICFQVTQTIAFFHSCLNPVLYVFV
P55085 HVYALYIVALCLSTLNSCIDPFVYYFV
P61073 TVHKWISITEALAFFHCCLNPILYAFL
Q92633 DVLAYEKFFLLLAEFNSAMNPIIYSYR
Q9H244 TLFYVKESTLWLTSLNACLDPFIYFFL
P20789 FYHYFYMLTNALFYVSSAINPILYNLV
P34998 SRVVFIYFNSFLESFQGFFVSVFYCFL
P43220 LRFIKLFTELSFTSFQGLMVAILYCFV
P47871 LRSAKLFFDLFLSSFQGLLVAVLYCFL
P41594 ----KIITMCFSVSLSATVALGCMFVP
Q13255 ----KIITTCFAVSLSVTVALGCMFTP
Q99835 PSLLVEKINLFAMFGTGIAMSTWVWTK

### MFS TM1

```
                                    TM1

                 IC                 h                EC
P08194           YRRLRWQIFLGIFFGYAAYYLVRKNFALAMPYLVEQG
P02920           MYYLKNTNFWMFGLFFFFYFFIMGAYFPFFPIWLHDI
P76350           SLSRARRAALGSFAGAVVDWYDFLLYGITAALVFNRE
P0AEX3           DTRRRIWAIVGASSGNLVEWFDFYVYSFCSLYFAHIF
Q5HRH0           DGNNAKKTVIATGIGNAMEWFDFGLYSYLAVIISKNF
O51798           GSPQQKKTFWACYSGWALDSFDMQMFSFLLPALTLTW
P0C0L7           DDGKLRKAITAASLGNAMEWFDFGVYGFVAYALGKVF
P37643           PINSRNKVLVASLIGTAIEFFDFYIYATAAVIVFPHI
P71369           VNSYGWKALIGSAVGYGMDGFDLLILGFMLSAISADL
P94131           GSHTWKIAFLFAFLALLVDGADLMLLSYSLNSIKAEF
P11166           VGGAVLGSLQFGYNTGVINAPQKVIEEFYNQTWVHRY
P0AGF4           NSSYIFSITLVATLGGLLFGYDTAVISGTVESLNTVF
P11551           GQSRSYIIPFALLCSLFFLWAVANNLNDILLPQFQQA
```

### MFS TM2

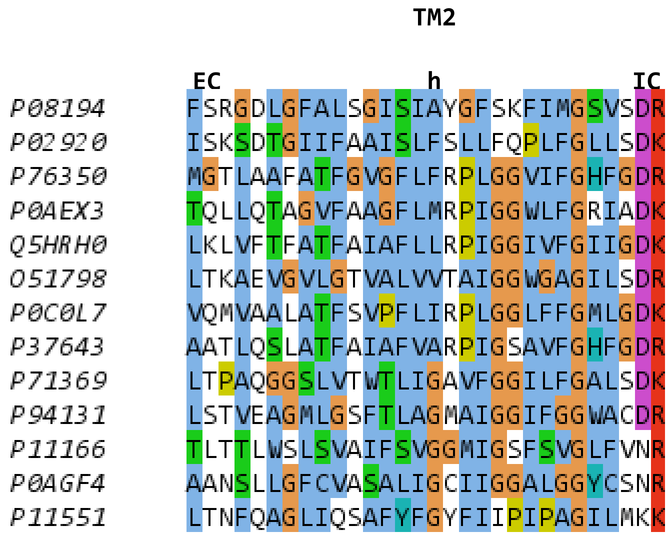

TM2
EC
h
IC
P08194 FSRGDLGFALSGISIAYGFSKFIMGSVSDR
P02920 ISKSDTGIIFAAISLFSLLFQPLFGLLSDK
P76350 MGTLAAFATFGVGFLFRPLGGVIFGHFGDR
P0AEX3 TQLLQTAGVFAAGFLMRPIGGWLFGRIADK
Q5HRH0 LKLVFTFATFAIAFLLRPIGGIVFGIIGDK
O51798 LTKAEVGVLGTVALVVTAIGGWGAGILSDR
P0C0L7 VQMVAALATFSVPFLIRPLGGLFFGMLGDK
P37643 AATLQSLATFAIAFVARPIGSAVFGHFGDR
P71369 LTPAQGGSLVTWTLIGAVFGGILFGALSDK
P94131 LSTVEAGMLGSFTLAGMAIGGIFGGWACDR
P11166 TLTTLWSLSVAIFSVGGMIGSFSVGLFVNR
P0AGF4 AANSLLGFCVASALIGCIIGGALGGYCSNR
P11551 LTNFQAGLIQSAFYFGYFIIPIPAGILMKK

### MFS TM3

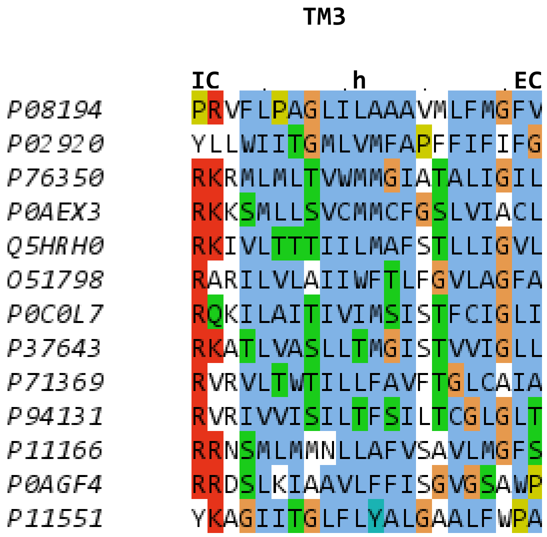
TM3
IC
h
EC
P08194 PRVFLPAGLILAAAVMLFMGFV
P02920 YLLWIITGMLVMFAPFFIFIFG
P76350 RKRMLMLTVWMMGIATALIGIL
P0AEX3 RKKSMLLSVCMMCFGSLVIACL
Q5HRH0 RKIVLTTTIILMAFSTLLIGVL
O51798 RARILVLAIIWFTLFGVLAGFA
P0C0L7 RQKILAITIVIMSISTFCIGLI
P37643 RKATLVASLLTMGISTVVIGLL
P71369 RVRVLTWTILLFAVFTGLCAIA
P94131 RVRIVVISILTFSILTCGLGLT
P11166 RRNSMLMMNLLAFVSAVLMGFS
P0AGF4 RRDSLKIAAVLFFISGVGSAWP
P11551 YKAGIITGLFLYALGAALFWPA

### MFS TM4

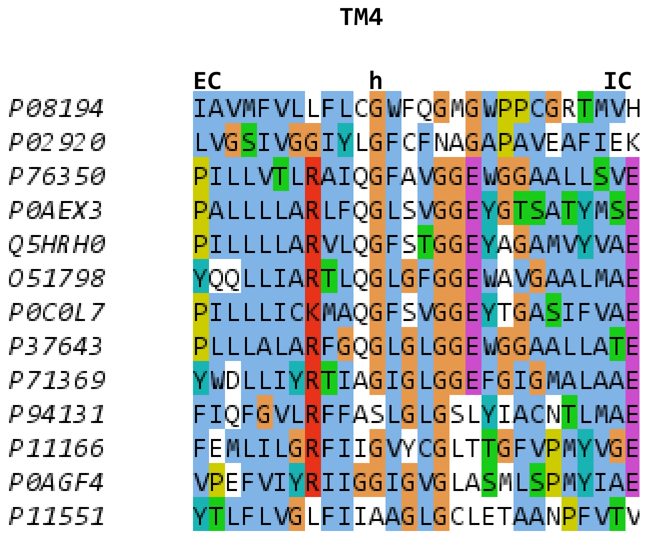

TM4
EC
h
IC
P08194 IAVMFVLLFLCGWFQGMGWPPCGRTMVH
P02920 LVGSIVGGIYLGFCFNAGAPAVEAFIEK
P76350 PILLVTLRAIQGFAVGGEWGGAALLSVE
P0AEX3 PALLLLARLFQGLSVGGEYGTSATYMSE
Q5HRH0 PILLLLARVLQGFSTGGEYAGAMVYVAE
O51798 YQQLLIARTLQGLGFGGEWAVGAALMAE
P0C0L7 PILLLICKMAQGFSVGGEYTGASIFVAE
P37643 PLLLALARFGQGLGLGGEWGGAALLATE
P71369 YWDLLIYRTIAGIGLGGEFGIGMALAAE
P94131 FIQFGVLRFFASLGLGSLYIACNTLMAE
P11166 FEMLILGRFIIGVYCGLTTGFVPMYVGE
P0AGF4 VPEFVIYRIIGGIGVGLASMLSPMYIAE
P11551 YTLFLVGLFIIAAGLGCLETAANPFVTV

### MFS TM5

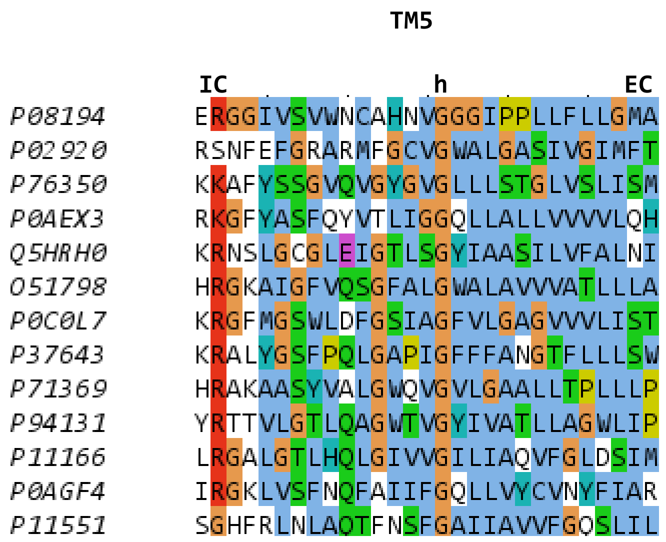
TM5
IC h EC
P08194 ERGGIVSVWNCAHNVGGGIPPLLFLLGMA
P02920 RSNFEFGRARMFGCVGWALGASIVGIMFT
P76350 KKAFYSSGVQVGYGVGLLLSTGLVSLISM
P0AEX3 RKGFYASFQYVTLIGGQLLALLVVVVLQH
Q5HRH0 KRNSLGCGLEIGTLSGYIAASILVFALNI
O51798 HRGKAIGFVQSGFALGWALAVVVATLLLA
P0C0L7 KRGFMGSWLDFGSIAGFVLGAGVVVLIST
P37643 KRALYGSFPQLGAPIGFFFANGTFLLLSW
P71369 HRAKAASYVALGWQVGVLGAALLTPLLLP
P94131 YRTTVLGTLQAGWTVGYIVATLLAGWLIP
P11166 LRGALGTLHQLGIVVGILIAQVFGLDSIM
P0AGF4 IRGKLVSFNQFAIIFGQLLVYCVNYFIAR
P11551 SGHFRLNLAQTFNSFGAIIAVVFGQSLIL

### MFS TM6

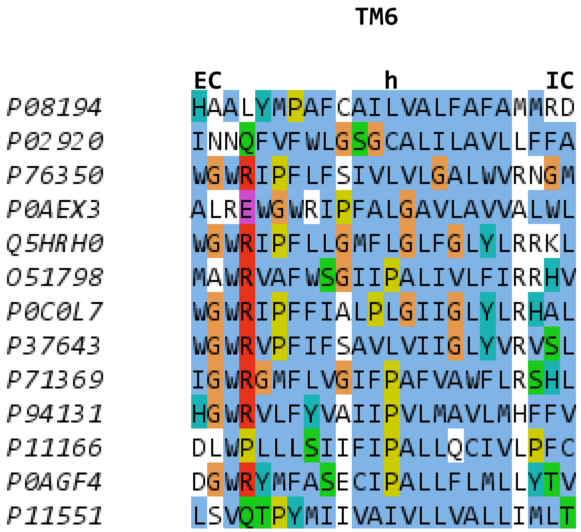
TM6
EC
h
IC
P08194 HAALYMPAFCAILVALFAFAMMRD
P02920 INNQFVFWLGSGCALILAVLLFFA
P76350 WGWRIPFLFSIVLVLGALWVRNGM
P0AEX3 ALREWGWRIPFALGAVLAVVALWL
Q5HRH0 WGWRIPFLLGMFLGLFGLYLRRKL
O51798 MAWRVAFWSGIIPALIVLFIRRHV
P0C0L7 WGWRIPFFIALPLGIIGLYLRHAL
P37643 WGWRVPFIFSAVLVIIGLYVRVSL
P71369 IGWRGMFLVGIFPAFVAWFLRSHL
P94131 HGWRVLFYVAIIPVLMAVLMHFFV
P11166 DLWPLLLSIIFIPALLQCIVLPFC
P0AGF4 DGWRYMFASECIPALLFLMLLYTV
P11551 LSVQTPYMIIVAIVLLVALLIMLT

### MFS TM7

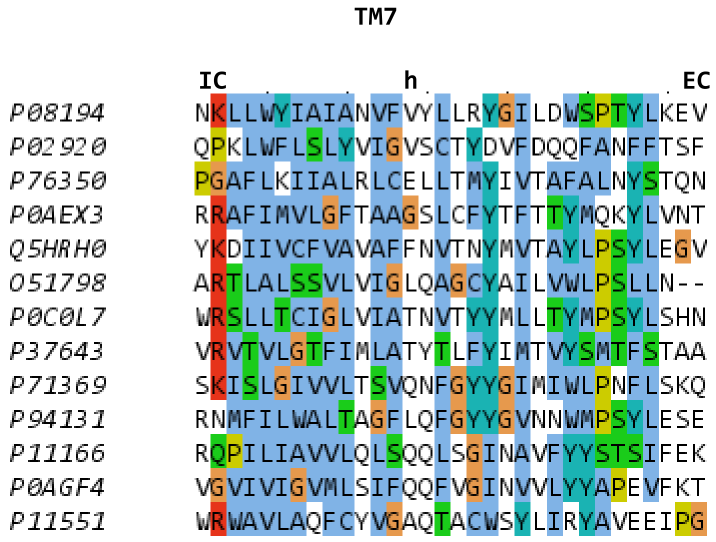
TM7
IC h EC
P08194 NKLLWYIAIANVFVYLLRYGILDWSPTYLKEV
P02920 QPKLWFLSLYVIGVSCTYDVFDQQFANFFTSF
P76350 PGAFLKIIALRLCELLTMYIVTAFALNYSTQN
P0AEX3 RRAFIMVLGFTAAGSLCFYTFTTYMQKYLVNT
Q5HRH0 YKDIIVCFVAVAFFNVTNYMVTAYLPSYLEGV
O51798 ARTLALSSVLVIGLQAGCYAILVWLPSLLN--
P0C0L7 WRSLLTCIGLVIATNVTYYMLLTYMPSYLSHN
P37643 VRVTVLGTFIMLATYTLFYIMTVYSMTFSTAA
P71369 SKISLGIVVLTSVQNFGYYGIMIWLPNFLSKQ
P94131 RNMFILWALTAGFLQFGYYGVNNWMPSYLESE
P11166 RQPILIAVVLQLSQQLSGINAVFYYSTSIFEK
P0AGF4 VGVIVIGVMLSIFQQFVGINVVLYYAPEVFKT
P11551 WRWAVLAQFCYVGAQTACWSYLIRYAVEEIPG

### MFS TM8

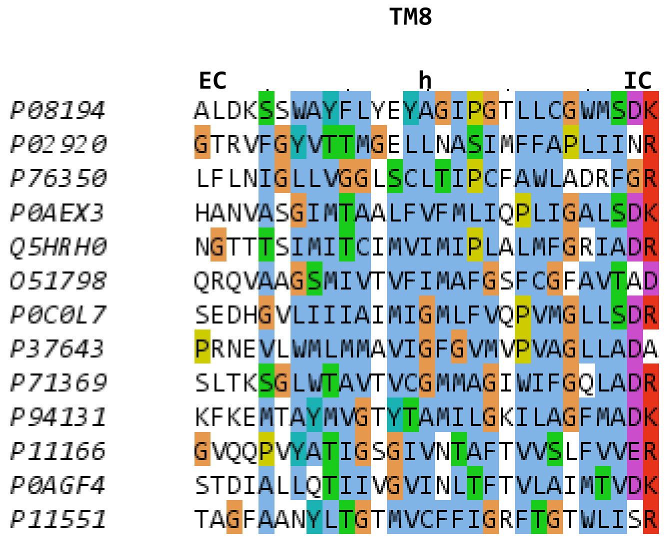

TM8
EC
h
IC
P08194 ALDKSSWAYFLYEYAGIPGTLLCGWMSDK
P02920 GTRVFGYVTTMGELLNASIMFFAPLIINR
P76350 LFLNIGLLVGGLSCLTIPCFAWLADRFGR
P0AEX3 HANVASGIMTAALFVFMLIQPLIGALSDK
Q5HRH0 NGTTTSIMITCIMVIMIPLALMFGRIADR
O51798 QRQVAAGSMIVTVFIMAFGSFCGFAVTAD
P0C0L7 SEDHGVLIIIAIMIGMLFVQPVMGLLSDR
P37643 PRNEVLWMLMMAVIGFGVMVPVAGLLADA
P71369 SLTKSGLWTAVTVCGMMAGIWIFGQLADR
P94131 KFKEMTAYMVGTYTAMILGKILAGFMADK
P11166 GVQQPVYATIGSGIVNTAFTVVSLFVVER
P0AGF4 STDIALLQTIIVGVINLTFTVLAIMTVDK
P11551 TAGFAANYLTGTMVCFFIGRFTGTWLISR

### MFS TM9

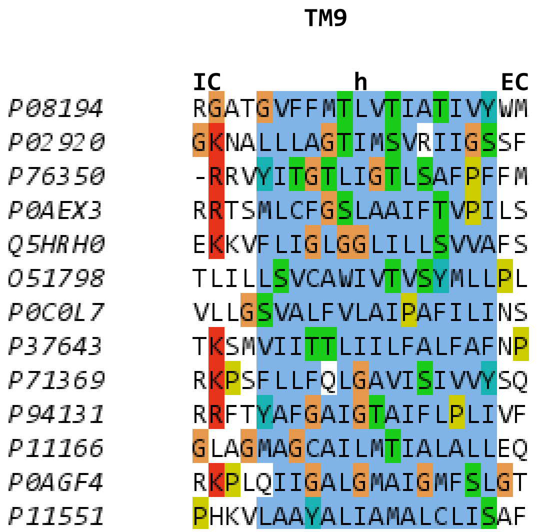

TM9
IC
h
EC
P08194 RGATGVFFMTLVTIATIVYWM
P02920 GKNALLLAGTIMSVRIIGSSF
P76350 -RRVYITGTLIGTLSAFPFFM
P0AEX3 RRTSMLCFGSLAAIFTVPILS
Q5HRH0 EKKVFLIGLGGLILLSVVAFS
O51798 TLILLSVCAWIVTVSYMLLPL
P0C0L7 VLLGSVALFVLAIPAFILINS
P37643 TKSMVIITTLIILFALFAFNP
P71369 RKPSFLLFQLGAVISIVVYSQ
P94131 RRFTYAFGAIGTAIFLPLIVF
P11166 GLAGMAGCAILMTIALALLEQ
P0AGF4 RKPLQIIGALGMAIGMFSLGT
P11551 PHKVLAAYALIAMALCLISAF

### MFS TM10

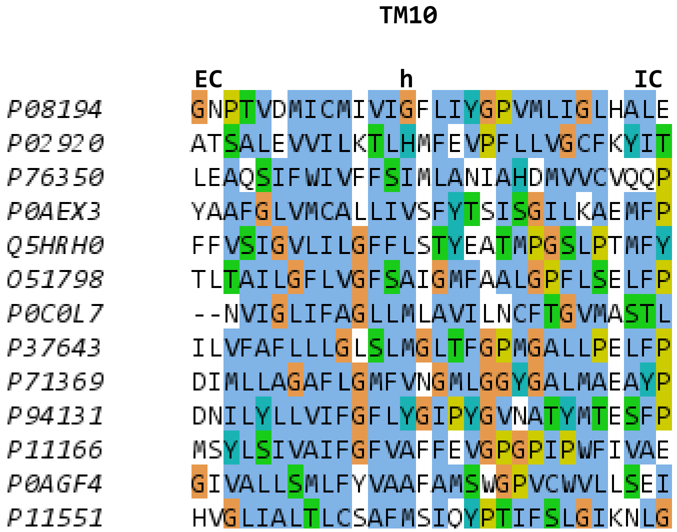

TM10
EC
h
IC
P08194 GNPTVDMICMIVIGFLIYGPVMLIGLHALE
P02920 ATSALEVVILKTLHMFEVPFLLVGCFKYIT
P76350 LEAQSIFWIVFFSIMLANIAHDMVVCVQQP
P0AEX3 YAAFGLVMCALLIVSFYTSISGILKAEMFP
Q5HRH0 FFVSIGVLILGFFLSTYEATMPGSLPTMFY
O51798 TLTAILGFLVGFSAIGMFAALGPFLSELFP
P0C0L7 --NVIGLIFAGLLMLAVILNCFTGVMASTL
P37643 ILVFAFLLLGLSLMGLTFGPMGALLPELFP
P71369 DIMLLAGAFLGMFVNGMLGGYGALMAEAYP
P94131 DNILYLLVIFGFLYGIPYGVNATYMTESFP
P11166 MSYLSIVAIFGFVAFFEVGPGPIPWFIVAE
P0AGF4 GIVALLSMLFYVAAFAMSWGPVCWVLLSEI
P11551 HVGLIALTLCSAFMSIQYPTIFSLGIKNLG

### MFS TM11

**TM11**

```
        IC            h            EC
P08194  AAGTAAGFTGLFGYLGGSVAASAIVGYTV
P02920  SATIYLVCFCFFKQLAMIFMSVLAGNMYE
P76350  RYSGAGVGYQVASVVGGGFTPFIAAALIT
P0AEX3  VRALGVGLSYAVANAIFGGSAEYVALSLK
Q5HRH0  RYRALAITFNVSVSLLGGTTPLIASYLVS
O51798  VRTTCMGFAYNVGKSIGAGSVVGVGVLST
P0C0L7  IRYSALAAAFNISVLVAGLTPTLAAWLVE
P37643  RYTGASFSYNVASILGASVAPYIAAWLQT
P71369  ARATAQNVLFNIGRAVGGFGPVVVGSVVL
P94131  IRGTAIGGAYNVGRLGAAIAPATIGFLAS
P11166  AAIAVAGFSNWTSNFIVGMCFQYVEQLCG
P0AGF4  IRGKALAIAVAAQWLANYFVSWTFPMMDK
P11551  QDTKYGSSFIVMTIIGGGIVTPVMGFVSD
```

### MFS TM12

**TM12**

```
                EC          h           IC
P08194          WDGGFMVMIGGSILAVILLIVVMIGEK
P02920          FQGAYLVLGLVALGFTLISVFTLSGPG
P76350          HSVAIYLLAGCLISAMTALLMKDSQRA
P0AEX3          ETAFFWYVTLMAVVAFLVSLMLHRKGK
Q5HRH0          PLTPAYYLTVISIIGFIVIALLHKSTA
O51798          ANAMGTFCLVAYAFAVFGIMLLPETRG
P0C0L7          LMMPAYYLMVVAVVGLITGVTMKETAN
P37643          LGAVGLYLAAMAGLTLIALLLTHETRH
P71369          FQTAIALLAIIYVIDMLATIFLIPELK
P94131          IGLGFVVMGAAYFICGVIPALFIKEKQ
P11166          --YVFIIFTVLLVLFFIFTYFKVPETK
P0AGF4          HFHNGFSYWIYGCMGVLAALFMWKFVP
P11551          IPTAELIPALCFAVIFIFARFRSQTAT
```

### PNuC TM0

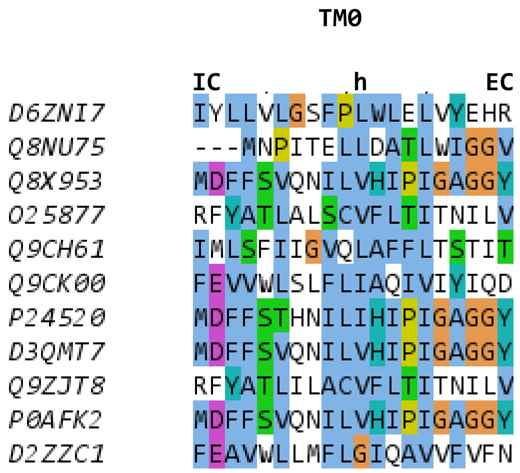
TM0
IC
h
EC
D6ZNI7 IYLLVLGSFPLWLELVYEHR
Q8NU75 ---MNPITELLDATLWIGGV
Q8X953 MDFFSVQNILVHIPIGAGGY
O25877 RFYATLALSCVFLTITNILV
Q9CH61 IMLSFIIGVQLAFFLTSTIT
Q9CK00 FEVVWLSLFLIAQIVIYIQD
P24520 MDFFSTHNILIHIPIGAGGY
D3QMT7 MDFFSVQNILVHIPIGAGGY
Q9ZJT8 RFYATLILACVFLTITNILV
P0AFK2 MDFFSVQNILVHIPIGAGGY
D2ZZC1 FEAVWLLMFLGIQAVVFVFN

### PNuC TM1

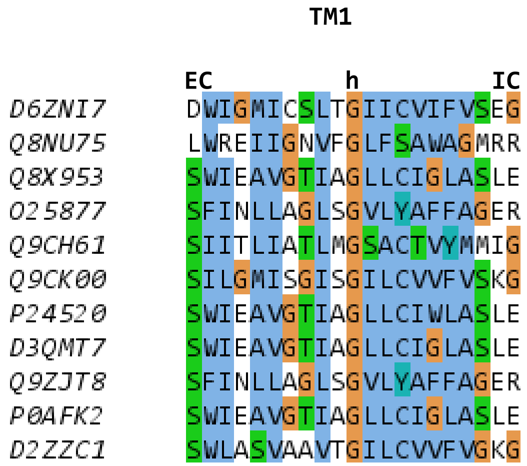
TM1
EC
h
IC
D6ZNI7 DWIGMICSLTGIICVIFVSEG
Q8NU75 LWREIIGNVFGLFSAWAGMRR
Q8X953 SWIEAVGTIAGLLCIGLASLE
O25877 SFINLLAGLSGVLYAFFAGER
Q9CH61 SIITLIATLMGSACTVYMMIG
Q9CK00 SILGMISGISGILCVVFVSKG
P24520 SWIEAVGTIAGLLCIWLASLE
D3QMT7 SWIEAVGTIAGLLCIGLASLE
Q9ZJT8 SFINLLAGLSGVLYAFFAGER
P0AFK2 SWIEAVGTIAGLLCIGLASLE
D2ZZC1 SWLASVAAVTGILCVVFVGKG

### PNuC TM2

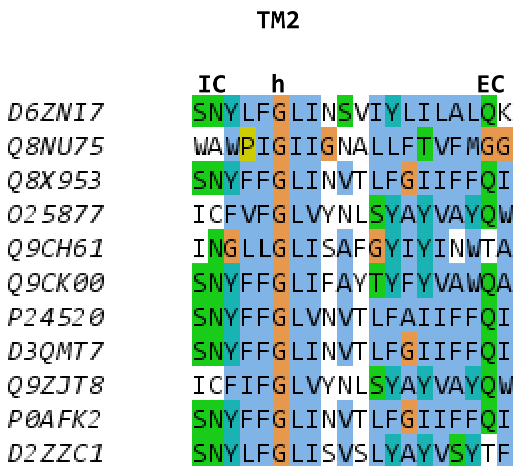

TM2
IC
h
EC
D6ZNI7 SNYLFGLINSVIYLILALQK
Q8NU75 WAWPIGIIGNALLFTVFMGG
Q8X953 SNYFFGLINVTLFGIIFFQI
O25877 ICFVFGLVYNLSYAYVAYQW
Q9CH61 INGLLGLISAFGYIYINWTA
Q9CK00 SNYFFGLIFAYTYFYVAWQA
P24520 SNYFFGLVNVTLFAIIFFQI
D3QMT7 SNYFFGLINVTLFGIIFFQI
Q9ZJT8 ICFIFGLVYNLSYAYVAYQW
P0AFK2 SNYFFGLINVTLFGIIFFQI
D2ZZC1 SNYLFGLISVSLYAYVSYTF

### PNuC TM3

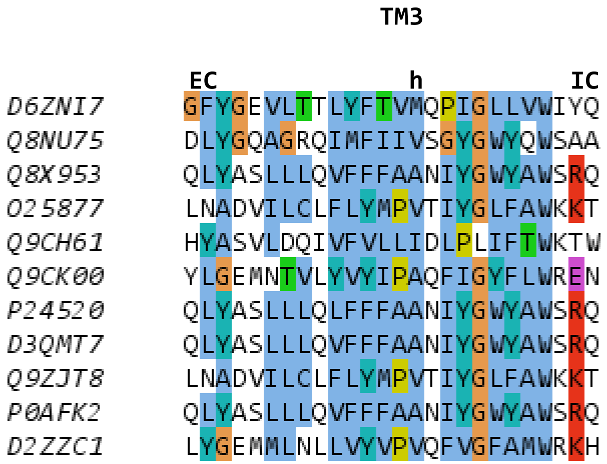

TM3
EC
h
IC
D6ZNI7 GFYGEVLTTLYFTVMQPIGLLVWIYQ
Q8NU75 DLYGQAGRQIMFIIVSGYGWYQWSAA
Q8X953 QLYASLLLQVFFFAANIYGWYAWSRQ
O25877 LNADVILCLFLYMPVTIYGLFAWKKT
Q9CH61 HYASVLDQIVFVLLIDLPLIFTWKTW
Q9CK00 YLGEMNTVLYVYIPAQFIGYFLWREN
P24520 QLYASLLLQLFFFAANIYGWYAWSRQ
D3QMT7 QLYASLLLQVFFFAANIYGWYAWSRQ
Q9ZJT8 LNADVILCLFLYMPVTIYGLFAWKKT
P0AFK2 QLYASLLLQVFFFAANIYGWYAWSRQ
D2ZZC1 LYGEMMLNLLVYVPVQFVGFAMWRKH

### PNuC TM4

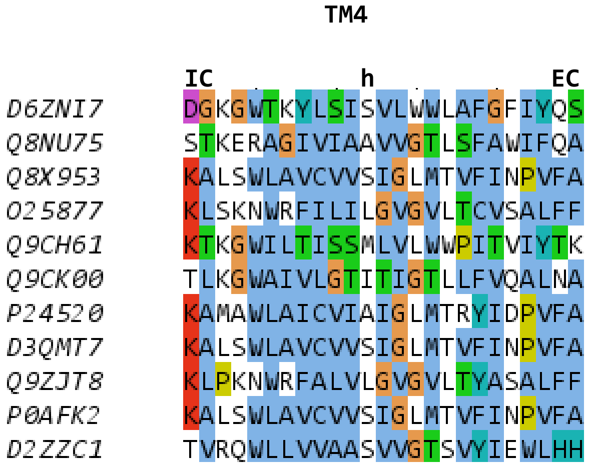

TM4
IC
h
EC
D6ZNI7 DGKGWTKYLSISVLWWLAFGFIYQS
Q8NU75 STKERAGIVIAAVVGTLSFAWIFQA
Q8X953 KALSWLAVCVVSIGLMTVFINPVFA
O25877 KLSKNWRFILILGVGVLTCVSALFF
Q9CH61 KTKGWILTISSMLVLWWPITVIYTK
Q9CK00 TLKGWAIVLGTITIGTLLFVQALNA
P24520 KAMAWLAICVIAIGLMTRYIDPVFA
D3QMT7 KALSWLAVCVVSIGLMTVFINPVFA
Q9ZJT8 KLPKNWRFALVLGVGVLTYASALFF
P0AFK2 KALSWLAVCVVSIGLMTVFINPVFA
D2ZZC1 TVRQWLLVVAASVVGTSVYIEWLHH

### PNuC TM5

```
                 TM5

        EC          h     IC
D6ZNI7  PYRDSITDATNGVGQILMTA
Q8NU75  PWADAWIFVGSILATYGMAR
Q8X953  PFWDSCMMVLSIVAMILMTR
O25877  LWAESFNFVIFIIAFILQVL
Q9CH61  PLWDAITLIIGAAASILVVR
Q9CK00  TGLDGLTTVIVVVAQLLMIL
P24520  PFWDSCMMVLSIVAMILMTR
D3QMT7  PFWDSCMMVLSIVAMILMTR
Q9ZJT8  LWAESFNFVIFIIAFILQVL
P0AFK2  PFWDSCMMVLSIVAMILMTR
D2ZZC1  PTLDGVTVVVSIVAQVLMIL
```

### PNuC TM6

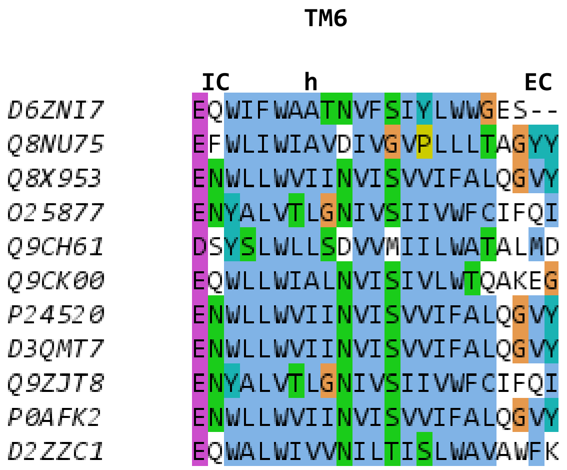
TM6
IC
h
EC
D6ZNI7 EQWIFWAATNVFSIYLWWGES--
Q8NU75 EFWLIWIAVDIVGVPLLLTAGYY
Q8X953 ENWLLWVIINVISVVIFALQGVY
O25877 ENYALVTLGNIVSIIVWFCIFQI
Q9CH61 DSYSLWLLSDVVMIILWATALMD
Q9CK00 EQWLLWIALNVISIVLWTQAKEG
P24520 ENWLLWVIINVISVVIFALQGVY
D3QMT7 ENWLLWVIINVISVVIFALQGVY
Q9ZJT8 ENYALVTLGNIVSIIVWFCIFQI
P0AFK2 ENWLLWVIINVISVVIFALQGVY
D2ZZC1 EQWALWIVVNILTISLWAVAWFK

### PNuC TM7

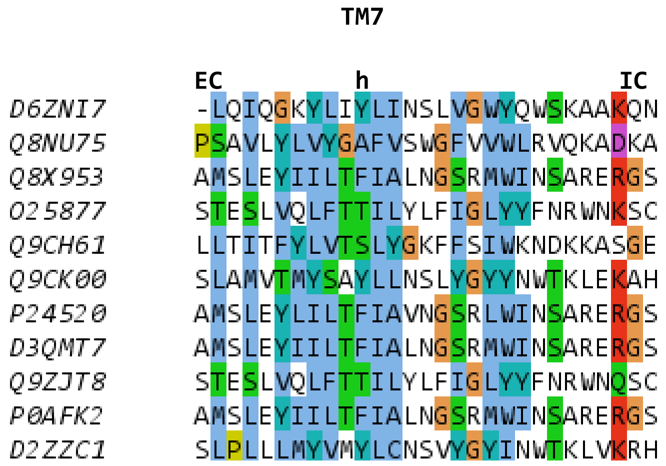
TM7
EC
h
IC
D6ZNI7 -LQIQGKYLIYLINSLVGWYQWSKAAKQN
Q8NU75 PSAVLYLVYGAFVSWGFVVWLRVQKADKA
Q8X953 AMSLEYIILTFIALNGSRMWINSARERGS
O25877 STESLVQLFTTILYLFIGLYYFNRWNKSC
Q9CH61 LLTITFYLVTSLYGKFFSIWKNDKKASGE
Q9CK00 SLAMVTMYSAYLLNSLYGYYNWTKLEKAH
P24520 AMSLEYLILTFIAVNGSRLWINSARERGS
D3QMT7 AMSLEYIILTFIALNGSRMWINSARERGS
Q9ZJT8 STESLVQLFTTILYLFIGLYYFNRWNQSC
P0AFK2 AMSLEYIILTFIALNGSRMWINSARERGS
D2ZZC1 SLPLLLMYVMYLCNSVYGYINWTKLVKRH

### SemiSweet TM1

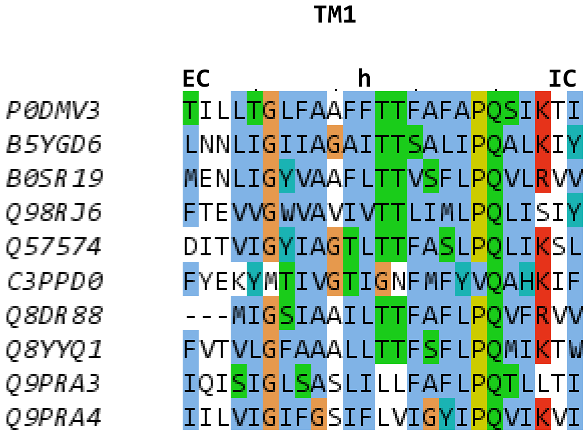

### SemiSweet TM2

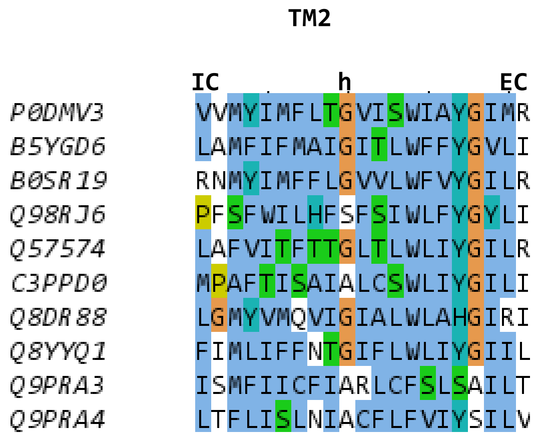

### SemiSweet TM3

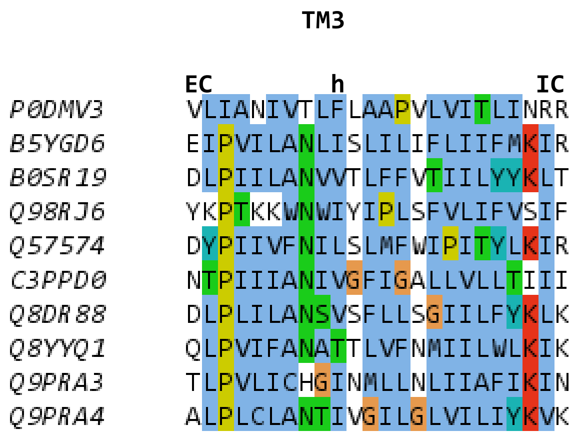

### Sweet TM1

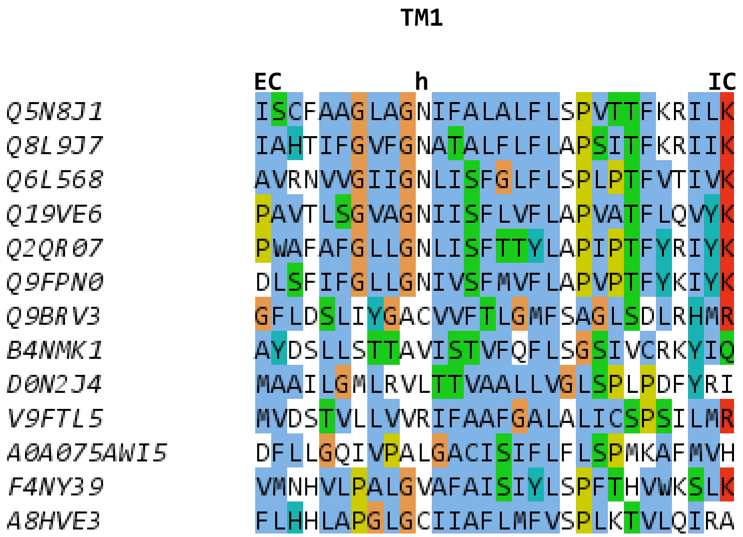
TM1
EC
h
IC
Q5N8J1 ISCFAAGLAGNIFALALFLSPVTTFKRILK
Q8L9J7 IAHTIFGVFGNATALFLFLAPSITFKRIIK
Q6L568 AVRNVVGIIGNLISFGLFLSPLPTFVTIVK
Q19VE6 PAVTLSGVAGNIISFLVFLAPVATFLQVYK
Q2QR07 PWAFAFGLLGNLISFTTYLAPIPTFYRIYK
Q9FPN0 DLSFIFGLLGNIVSFMVFLAPVPTFYKIYK
Q9BRV3 GFLDSLIYGACVVFTLGMFSAGLSDLRHMR
B4NMK1 AYDSLLSTTAVISTVFQFLSGSIVCRKYIQ
D0N2J4 MAAILGMLRVLTTVAALLVGLSPLPDFYRI
V9FTL5 MVDSTVLLVVRIFAAFGALALICSPSILMR
A0A075AWI5 DFLLGQIVPALGACISIFLFLSPMKAFMVH
F4NY39 VMNHVLPALGVAFAISIYLSPFTHVWKSLK
A8HVE3 FLHHLAPGLGCIIAFLMFVSPLKTVLQIRA

### Sweet TM2

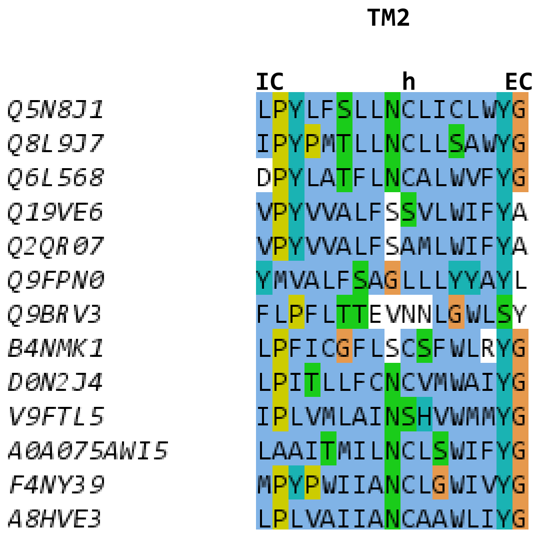
TM2
IC h EC
Q5N8J1 LPYLFSLLNCLICLWYG
Q8L9J7 IPYPMTLLNCLLSAWYG
Q6L568 DPYLATFLNCALWVFYG
Q19VE6 VPYVVALFSSVLWIFYA
Q2QR07 VPYVVALFSAMLWIFYA
Q9FPN0 YMVALFSAGLLLYYAYL
Q9BRV3 FLPFLTTEVNNLGWLSY
B4NMK1 LPFICGFLSCSFWLRYG
D0N2J4 LPITLLFCNCVMWAIYG
V9FTL5 IPLVMLAINSHVWMMYG
A0A075AWI5 LAAITMILNCLSWIFYG
F4NY39 MPYPWIIANCLGWIVYG
A8HVE3 LPLVAIIANCAAWLIYG

### Sweet TM3

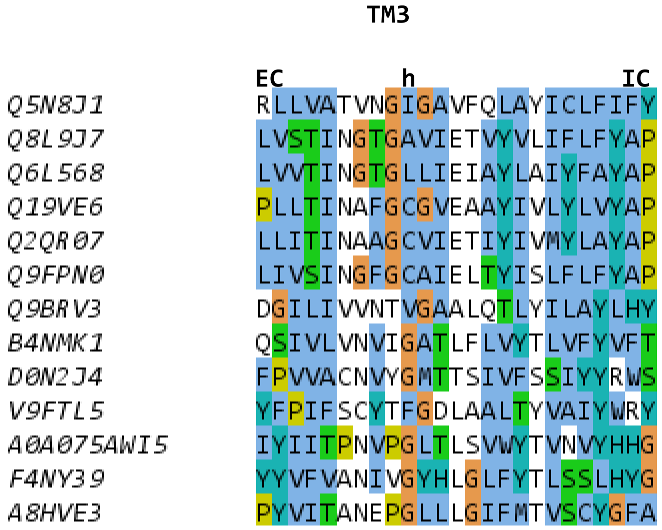

TM3
EC
h
IC
Q5N8J1 RLLVATVNGIGAVFQLAYICLFIFY
Q8L9J7 LVSTINGTGAVIETVYVLIFLFYAP
Q6L568 LVVTINGTGLLIEIAYLAIYFAYAP
Q19VE6 PLLTINAFGCGVEAAYIVLYLVYAP
Q2QR07 LLITINAAGCVIETIYIVMYLAYAP
Q9FPN0 LIVSINGFGCAIELTYISLFLFYAP
Q9BRV3 DGILIVVNTVGAALQTLYILAYLHY
B4NMK1 QSIVLVNVIGATLFLVYTLVFYVFT
D0N2J4 FPVVACNVYGMTTSIVFSSIYYRWS
V9FTL5 YFPIFSCYTFGDLAALTYVAIYWRY
A0A075AWI5 IYIITPNVPGLTLSVWYTVNVYHHG
F4NY39 YYVFVANIVGYHLGLFYTLSSLHYG
A8HVE3 PYVITANEPGLLLGIFMTVSCYGFA

### Sweet TM4

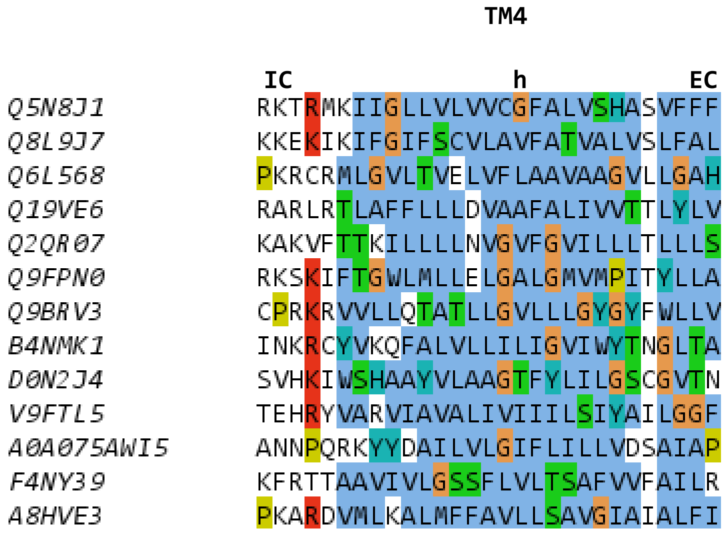
TM4
IC
h
EC
Q5N8J1 RKTRMKIIGLLVLVVCGFALVSHASVFFF
Q8L9J7 KKEKIKIFGIFSCVLAVFATVALVSLFAL
Q6L568 PKRCRMLGVLTVELVFLAAVAAGVLLGAH
Q19VE6 RARLRTLAFFLLLDVAAFALIVVTTLYLV
Q2QR07 KAKVFTTKILLLLNVGVFGVILLLTLLLS
Q9FPN0 RKSKIFTGWLMLLELGALGMVMPITYLLA
Q9BRV3 CPRKRVVLLQTATLLGVLLLGYGYFWLLV
B4NMK1 INKRCYVKQFALVLLILIGVIWYTNGLTA
D0N2J4 SVHKIWSHAAYVLAAGTFYLILGSCGVTN
V9FTL5 TEHRYVARVIAVALIVIIILSIYAILGGF
A0A075AWI5 ANNPQRKYYDAILVLGIFLILLVDSAIAP
F4NY39 KFRTTAAVIVLGSSFLVLTSAFVVFAILR
A8HVE3 PKARDVMLKALMFFAVLLSAVGIAIALFI

### Sweet TM5

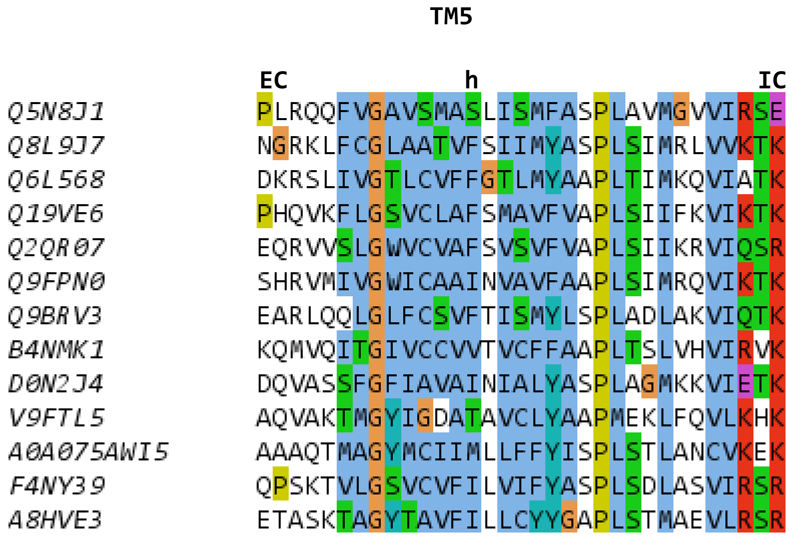
TM5
EC
h
IC
Q5N8J1 PLRQQFVGAVSMASLISMFASPLAVMGVVIRSE
Q8L9J7 NGRKLFCGLAATVFSIIMYASPLSIMRLVVKTK
Q6L568 DKRSLIVGTLCVFFGTLMYAAPLTIMKQVIATK
Q19VE6 PHQVKFLGSVCLAFSMAVFVAPLSIIFKVIKTK
Q2QR07 EQRVVSLGWVCVAFSVSVFVAPLSIIKRVIQSR
Q9FPN0 SHRVMIVGWICAAINVAVFAAPLSIMRQVIKTK
Q9BRV3 EARLQQLGLFCSVFTISMYLSPLADLAKVIQTK
B4NMK1 KQMVQITGIVCCVVTVCFFAAPLTSLVHVIRVK
D0N2J4 DQVASSFGFIAVAINIALYASPLAGMKKVIETK
V9FTL5 AQVAKTMGYIGDATAVCLYAAPMEKLFQVLKHK
A0A075AWI5 AAAQTMAGYMCIIMLLFFYISPLSTLANCVKEK
F4NY39 QPSKTVLGSVCVFILVIFYASPLSDLASVIRSR
A8HVE3 ETASKTAGYTAVFILLCYYGAPLSTMAEVLRSR

### Sweet TM6

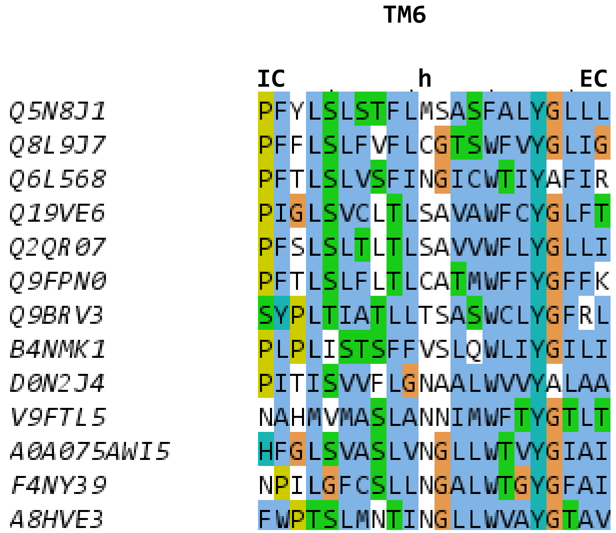
TM6
IC
h
EC
Q5N8J1 PFYLSLSTFLMSASFALYGLLL
Q8L9J7 PFFLSLFVFLCGTSWFVYGLIG
Q6L568 PFTLSLVSFINGICWTIYAFIR
Q19VE6 PIGLSVCLTLSAVAWFCYGLFT
Q2QR07 PFSLSLTLTLSAVVWFLYGLLI
Q9FPN0 PFTLSLFLTLCATMWFFYGFFK
Q9BRV3 SYPLTIATLLTSASWCLYGFRL
B4NMK1 PLPLISTSFFVSLQWLIYGILI
D0N2J4 PITISVVFLGNAALWVVYALAA
V9FTL5 NAHMVMASLANNIMWFTYGTLT
A0A075AWI5 HFGLSVASLVNGLLWTVYGIAI
F4NY39 NPILGFCSLLNGALWTGYGFAI
A8HVE3 FWPTSLMNTINGLLWVAYGTAV

### Sweet TM7

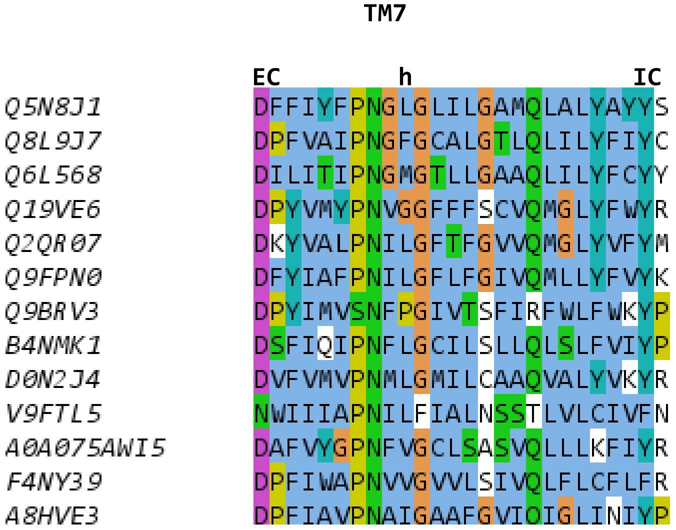
TM7
EC
h
IC
Q5N8J1 DFFIYFPNGLGLILGAMQLALYAYYS
Q8L9J7 DPFVAIPNGFGCALGTLQLILYFIYC
Q6L568 DILITIPNGMGTLLGAAQLILYFCYY
Q19VE6 DPYVMYPNVGGFFFSCVQMGLYFWYR
Q2QR07 DKYVALPNILGFTFGVVQMGLYVFYM
Q9FPN0 DFYIAFPNILGFLFGIVQMLLYFVYK
Q9BRV3 DPYIMVSNFPGIVTSFIRFWLFWKYP
B4NMK1 DSFIQIPNFLGCILSLLQLSLFVIYP
D0N2J4 DVFVMVPNMLGMILCAAQVALYVKYR
V9FTL5 NWIIIAPNILFIALNSSTLVLCIVFN
A0A075AWI5 DAFVYGPNFVGCLSASVQLLLKFIYR
F4NY39 DPFIWAPNVVGVVLSIVQLFLCFLFR
A8HVE3 DPFIAVPNAIGAAFGVIQIGLINIYP

### Tric TM1

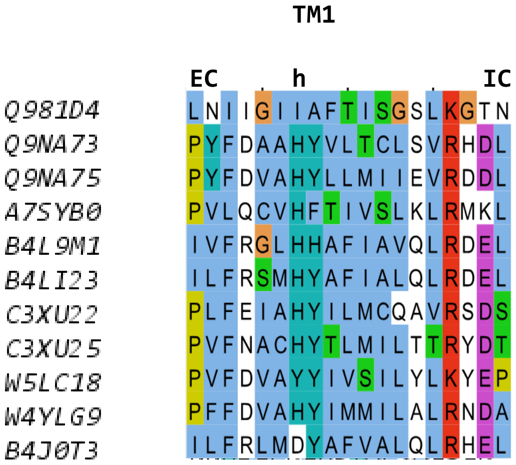
TM1
EC
h
IC
Q981D4 LNIIGIIAFTISGSLKGTN
Q9NA73 PYFDAAHYVLTCLSVRHDL
Q9NA75 PYFDVAHYLLMIIEVRDDL
A7SYB0 PVLQCVHFTIVSLKLRMKL
B4L9M1 IVFRGLHHAFIAVQLRDEL
B4LI23 ILFRSMHYAFIALQLRDEL
C3XU22 PLFEIAHYILMCQAVRSDS
C3XU25 PVFNACHYTLMILTTRYDT
W5LC18 PVFDVAYYIVSILYLKYEP
W4YLG9 PFFDVAHYIMMILALRNDA
B4J0T3 ILFRLMDYAFVALQLRHEL

### Tric TM2

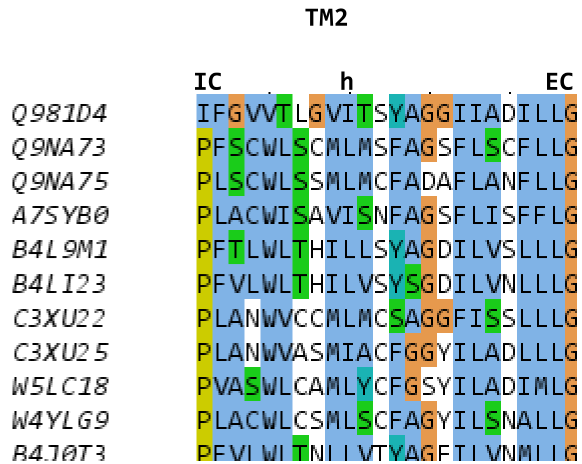
TM2
IC
h
EC
Q981D4 IFGVVTLGVITSYAGGIIADILLG
Q9NA73 PFSCWLSCMLMSFAGSFLSCFLLG
Q9NA75 PLSCWLSSMLMCFADAFLANFLLG
A7SYB0 PLACWISAVISNFAGSFLISFFLG
B4L9M1 PFTLWLTHILLSYAGDILVSLLLG
B4LI23 PFVLWLTHILVSYSGDILVNLLLG
C3XU22 PLANWVCCMLMCSAGGFISSLLLG
C3XU25 PLANWVASMIACFGGYILADLLLG
W5LC18 PVASWLCAMLYCFGSYILADIMLG
W4YLG9 PLACWLCSMLSCFAGYILSNALLG

### Tric TM3

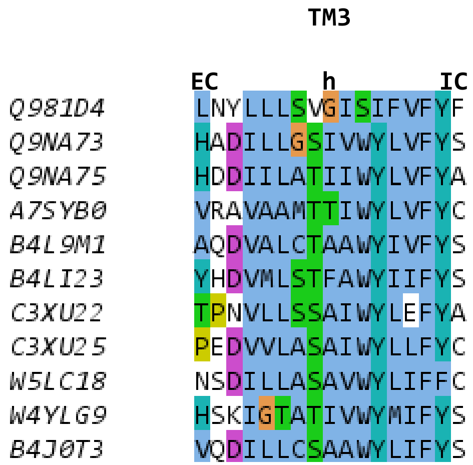

TM3
EC
h
IC
Q981D4 LNYLLLSVGISIFVFYF
Q9NA73 HADILLGSIVWYLVFYS
Q9NA75 HDDIILATIIWYLVFYA
A7SYB0 VRAVAAMTTIWYLVFYC
B4L9M1 AQDVALCTAAWYIVFYS
B4LI23 YHDVMLSTFAWYIIFYS
C3XU22 TPNVLLSSAIWYLEFYA
C3XU25 PEDVVLASAIWYLLFYC
W5LC18 NSDILLASAVWYLIFFC
W4YLG9 HSKIGTATIVWYMIFYS
B4J0T3 VQDILLCSAAWYLIFYS

### Tric TM4

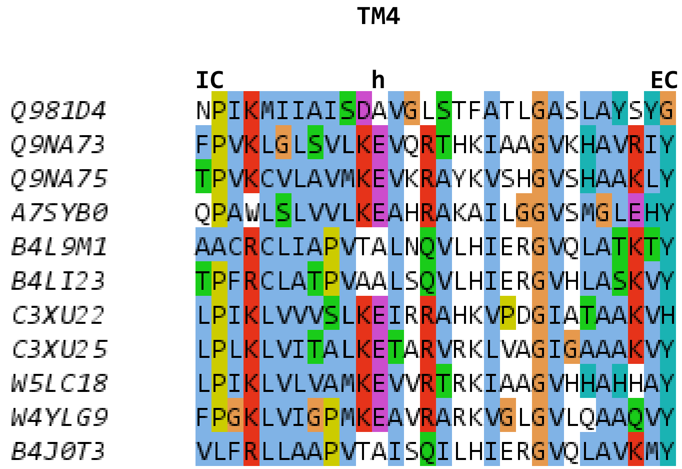
TM4
IC h EC
Q981D4 NPIKMIIAISDAVGLSTFATLGASLAYSYG
Q9NA73 FPVKLGLSVLKEVQRTHKIAAGVKHAVRIY
Q9NA75 TPVKCVLAVMKEVKRAYKVSHGVSHAAKLY
A7SYB0 QPAWLSLVVLKEAHRAKAILGGVSMGLEHY
B4L9M1 AACRCLIAPVTALNQVLHIERGVQLATKTY
B4LI23 TPFRCLATPVAALSQVLHIERGVHLASKVY
C3XU22 LPIKLVVVSLKEIRRAHKVPDGIATAAKVH
C3XU25 LPLKLVITALKETARVRKLVAGIGAAAKVY
W5LC18 LPIKLVLVAMKEVVRTRKIAAGVHHAHHAY
W4YLG9 FPGKLVIGPMKEAVRARKVGLGVLQAAQVY
B4J0T3 VLFRLLAAPVTAISQILHIERGVQLAVKMY

### Tric TM5

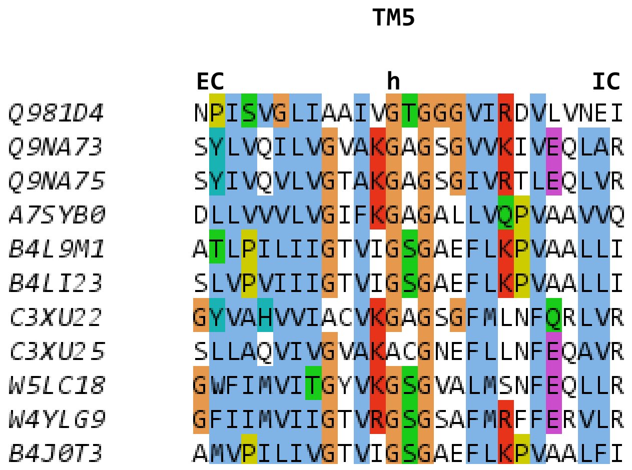
TM5
EC
h
IC
Q981D4 NPISVGLIAAIVGTGGGVIRDVLVNEI
Q9NA73 SYLVQILVGVAKGAGSGVVKIVEQLAR
Q9NA75 SYIVQVLVGTAKGAGSGIVRTLEQLVR
A7SYB0 DLLVVVLVGIFKGAGALLVQPVAAVVQ
B4L9M1 ATLPILIIGTVIGSGAEFLKPVAALLI
B4LI23 SLVPVIIIGTVIGSGAEFLKPVAALLI
C3XU22 GYVAHVVIACVKGAGSGFMLNFQRLVR
C3XU25 SLLAQVIVGVAKACGNEFLLNFEQAVR
W5LC18 GWFIMVITGYVKGSGVALMSNFEQLLR
W4YLG9 GFIIMVIIGTVRGSGSAFMRFFERVLR
B4J0T3 AMVPILIVGTVIGSGAEFLKPVAALFI

### Tric TM6

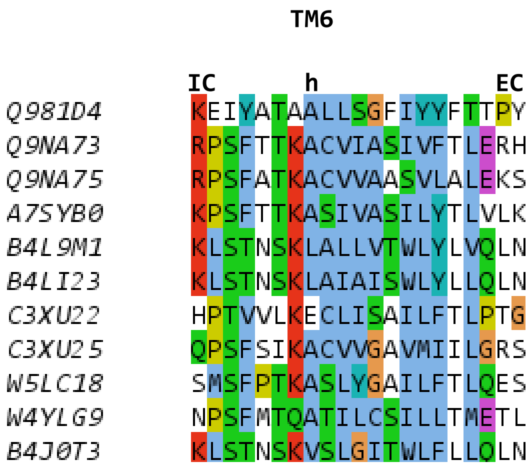
TM6
IC
h
EC
Q981D4 KEIYATAALLSGFIYYFTTPY
Q9NA73 RPSFTTKACVIASIVFTLERH
Q9NA75 RPSFATKACVVAASVLALEKS
A7SYB0 KPSFTTKASIVASILYTLVLK
B4L9M1 KLSTNSKLALLVTWLYLVQLN
B4LI23 KLSTNSKLAIAISWLYLLQLN
C3XU22 HPTVVLKECLISAILFTLPTG
C3XU25 QPSFSIKACVVGAVMIILGRS
W5LC18 SMSFPTKASLYGAILFTLQES
W4YLG9 NPSFMTQATILCSILLTMETL
B4J0T3 KLSTNSKVSLGITWLFLLQLN

### Tric TM7

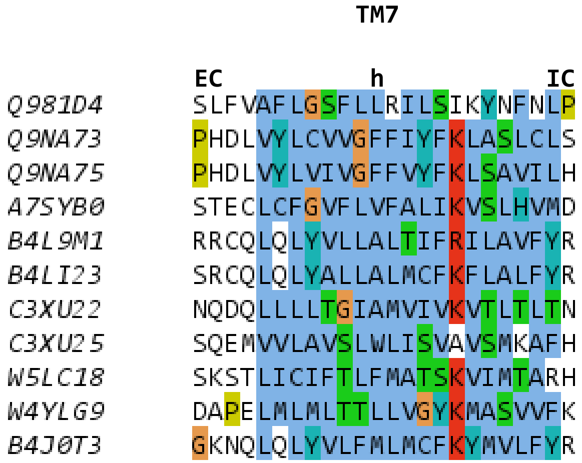

TM7
EC
h
IC
Q981D4 SLFVAFLGSFLLRILSIKYNFNLP
Q9NA73 PHDLVYLCVVGFFIYFKLASLCLS
Q9NA75 PHDLVYLVIVGFFVYFKLSAVILH
A7SYB0 STECLCFGVFLVFALIKVSLHVMD
B4L9M1 RRCQLQLYVLLALTIFRILAVFYR
B4LI23 SRCQLQLYALLALMCFKFLALFYR
C3XU22 NQDQLLLLTGIAMVIVKVTLTLTN
C3XU25 SQEMVVLAVSLWLISVAVSMKAFH
W5LC18 SKSTLICIFTLFMATSKVIMTARH
W4YLG9 DAPELMLMLTTLLVGYKMASVVFK
B4J0T3 GKNQLQLYVLFMLMCFKYMVLFYR
